# Genome mining leads to the identification of a stable and promiscuous Baeyer-Villiger monooxygenase from a thermophilic microorganism

**DOI:** 10.1101/2024.04.05.588253

**Authors:** Amir R. Bunyat-zada, Stephan E. Ducharme, Maria E. Cleveland, Esther R. Hoffman, Graeme W. Howe

## Abstract

Baeyer-Villiger monooxygenases are NAD(P)H-dependent flavoproteins that catalyze oxygen insertion reactions which convert ketones to valuable esters and lactones. While these enzymes offer an appealing alternative to traditional Baeyer-Villiger oxidations, these proteins tend to be either too unstable or exhibit too narrow of a substrate scope for implementation as industrial biocatalysts. Here, sequence similarity networks were used to search for novel Baeyer-Villiger monooxygenases that are both stable and substrate promiscuous. Our genome mining efforts led to the identification of an enzyme from *Chloroflexota* bacterium (strain G233) dubbed *ssn*BVMO that exhibits i) the highest melting temperature recorded to date for a naturally sourced Baeyer-Villiger monooxygenase, ii) a remarkable kinetic stability across a wide range of conditions, and iii) a broad substrate scope that includes linear aliphatic, aromatic, and sterically bulky ketones. Kinetic characterization of this enzyme was undertaken to identify the optimal conditions for *ssn*BVMO catalysis, and a subsequent quantitative assay using propiophenone as a substrate afforded more than 95% conversion. To spur the implementation of this enzyme as an oxidative biocatalyst, several fusion proteins were constructed that linked *ssn*BVMO to a thermostable phosphite dehydrogenase. These self-sufficient enzymes can recycle NADPH and permit oxidations to be run with sub-stoichiometric quantities of this expensive cofactor. Extensive characterization of these fusion enzymes permitted identification of PTDH-L1-*ssn*BVMO as the most promising oxidative biocatalyst. Results described herein demonstrate that this new monooxygenase has significant potential as a useful industrial biocatalyst for Baeyer-Villiger oxidations.

## Introduction

Baeyer Villiger (BV) reactions describe the oxidation of ketones to esters or lactones, typically using peroxides or peroxyacids as the oxidant.^1^ While these transformations are synthetically useful, the requirement for peroxide/peroxyacid oxidants poses significant limitations, as these reagents can be costly, shock-sensitive, and lead to the production of stoichiometric amounts of hazardous waste.^1^ Attempts to leverage hydrogen peroxide as an oxidant typically necessitate the use of metallo-or organo-catalysts.^2,3,4^ Unfortunately, these essential catalysts often require intricate syntheses, and the subsequent BV reactions still fail to address the issue of stoichiometric waste production.^5^ Furthermore, although BV oxidations serve as a valuable tool for synthetic chemists, reports on asymmetric variants remain scarce, and those that are available demonstrate restricted substrate scopes, thereby limiting their applicability.^2,5^ Thus, despite several decades of advances, challenges persist in the current approach to BV oxidations.

Baeyer-Villiger monooxygenases (BVMOs) are enzymes that can facilitate greener alternatives to traditional BV oxidations.^5^ These enzymes catalyze BV reactions using molecular oxygen as the sole oxidant and generate water as the only by-product (Scheme 1).^5^ BVMOs are Class B flavoprotein monooxygenases (FMOs) that are generally proficient at oxidizing carbon atoms and some heteroatoms, with the BVMO-catalyzed oxidation of sulfur and nitrogen atoms garnering considerable interest.^6^ This protein family has evolved to allow the reaction of O2 with carbon on organic compounds via a concerted reaction that is formally spin-forbidden.^6^ This is proposed to be achieved by a one-electron transfer from the reduced form of the flavin cofactor to the molecular oxygen to form a superoxide-flavin radical pair (Scheme 1).^7^ The subsequent spin-inversion results in the formation of reduced oxygen.^7^ BVMOs, like other FMOs, usually maintain the reduced oxygen in a deprotonated state, enabling this group to conduct nucleophilic oxygenation.^8^ The generally accepted mechanism of the BVMO-catalyzed transformation is shown in Scheme 1.

**Scheme 1.**
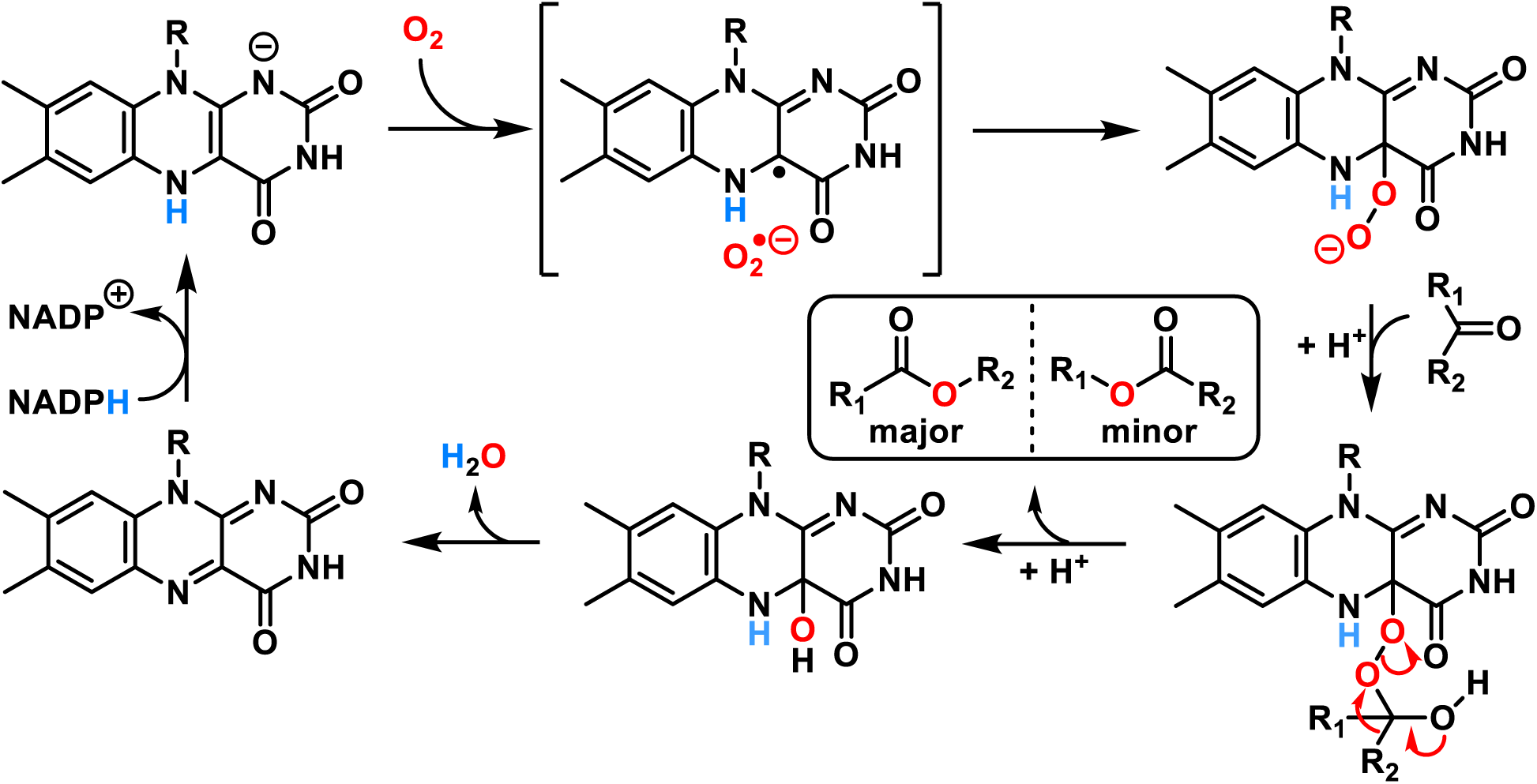
The proposed catalytic cycle for BVMOs. The regiochemistry of BVMO-catalyzed reactions is governed by i) the relative stability of the positive charge that develops on the migrating group,^9^ and ii) the orientation of each carbon-carbon bond relative to the labile peroxy group, with the anti-periplanar C-C bond preferentially migrating.^5,10^

Despite the enormous potential environmental benefits, BVMOs have not yet seen widespread industrial implementation. While the viability of utilizing both isolated BVMOs and whole cells expressing BVMOs as industrial biocatalysts has been demonstrated, several common obstacles may render these biocatalytic approaches less economically competitive.^11,12^ These include, but are not limited to, poor stability and/or solubility of the enzyme, low expression levels, dependence on stoichiometric quantities of expensive cofactors (NADPH), oxygen availability, and inhibition by substrates and/or products.^5,13–18^ In many instances, directed evolution has been used to alter natural BVMOs into novel biocatalysts that address some subset of these obstacles. For instance, Codexis subjected a cyclohexanone monooxygenase (CHMO) from *Acinetobacter* sp. strain NCIB 9871 to 19 rounds of directed evolution before the resulting variant was implemented in their chemoenzymatic synthesis of esomeprazole.^19^ This heavily mutagenized CHMO carried out the desired BV reaction with a 140,000-fold increase in productivity, vastly improved enantioselectivity (>99% ee), and a total turnover number (TTN) of approximately 9000.^19^

Since their initial discovery, BVMOs have been consistently challenged by poor thermostability. While cyclohexanone monooxygenase (*Ac*CHMO; UniProt ID: P12015) was first isolated from *Acinetobacter calcoaceticus* NCIMB 9871 in 1976, structural information was not obtained until 2019 with a variant *Ac*CHMO bearing 8 stabilizing mutations.^20,21^ Despite the instability of the wildtype enzyme, *Ac*CHMO has attracted significant attention for its’ broad substrate scope and is now considered a prototypical type 1 BVMO.^5^ Naturally thermostable BVMOs also exist, with phenylacetone monooxygenase (PAMO; UniProt ID: Q47PU3) from *Thermobifida fusca* serving as an exemplary system.^22^ These two archetypal enzymes serve as the foundational elements for a dichotomy among naturally occurring BVMOs: in general, BVMOs are *either* stable (PAMO) *or* promiscuous (*Ac*CHMO).^5^ We hypothesized that Nature had likely evolved enzymes that exhibit both the stability of PAMOs and the promiscuity of *Ac*CHMOs. Given that mutations introduced over the course of directed evolution to improve a desired activity of an enzyme are often accompanied by compensatory decreases in protein stability,^23,24^ we hypothesized that a naturally thermostable, promiscuous BVMO could serve as an excellent starting point for further protein engineering efforts. To this end, we employed sequence similarity networks (SSNs) to identify proteins from thermophilic host organisms that bear significant similarity to *Ac*CHMOs. Here, we describe the identification and subsequent characterization of one such BVMO from *Chloroflexota* bacterium strain G233 dubbed *ssn*BVMO.^25^

Following the identification of *ssn*BVMO, we sought to address another major hurdle for the large-scale implementation of BVMOs: the requirement for stoichiometric quantities of the expensive NADPH cofactor. Previous work has overcome this issue by fusing BVMOs to coenzyme regenerating enzymes (CREs).^26,27^ To maximize the potential utility of our novel BVMO, several fusions were designed that linked *ssn*BVMO to a thermostable phosphite dehydrogenase (17X-PTDH).^28^ This enzyme converts NAD(P)^+^ to NAD(P)H while oxidizing phosphite (a cheap, sacrificial substrate) to phosphate.^28,29^ Here, we demonstrate that the resulting fusion proteins are self-sufficient BVMOs that can convert ketones to esters with sub-stoichiometric quantities of NADPH. Collectively, this work demonstrates that *ssn*BVMO exhibits i) a higher melting temperature (*T*_m_) than any known naturally occurring BVMO, ii) an exceptional kinetic stability across a wide range of conditions, iii) a relatively broad substrate scope, and iv) can be fused to CREs to generate self-sufficient oxidative biocatalysts.

## Results & Discussion

### Using sequence similarity networks to identify a thermostable BVMO variant

Both PAMO and *Ac*CHMO belong to the FAD/NAD(P)-binding domain superfamily, but a significant portion of the N-terminus of PAMO is annotated as a “pyridine nucleotide-disulphide oxidoreductase” (PF13738), whereas the N-terminus of *Ac*CHMO is described as a “flavin-binding monooxygenase-like” protein (PF00743). To ensure that members of both families were represented in the bioinformatic analysis, the Enzyme Function Initiative’s Enzyme Similarity Tool (EFI-EST) was used to carry out an all-by-all BLAST of all sequences found within the UniRef90 database that belong to either Pfam.

An initial SSN was constructed with an alignment score (AS) of 140, corresponding to a %ID cut-off of approximately 35% (Figs S1 – S2). As the largest cluster in this network contained both the PAMO and *Ac*CHMO sequences, a daughter network was constructed from this cluster, and the AS of the resulting network was gradually increased until these sequences partitioned into two separate clusters. While this partitioning occurred with an AS of 175 (∼39 %ID cut-off), the resulting *Ac*CHMO-containing cluster did not contain any sequences from thermophilic organisms and only contained a few unannotated genes from thermotolerant organisms (i.e., *Aspergillus thermomutatus*). To identify BVMOs that exhibit the stability of PAMO and the promiscuity of *Ac*CHMO, the AS was reduced to 170 (∼38 %ID cut-off) to give rise to one cluster that contained PAMO, *Ac*CHMO, and nearly 3,500 functionally unannotated sequences that are similar to both of these BVMOs (Fig S3). Sequences from this cluster were then prioritized based on the reported growth temperatures of the host organisms. While several sequences from thermotolerant (i.e., *Minwuia thermotolerans, Bacillus thermotolerans*) and moderately thermophilic organisms (i.e., *Pseudonocardia thermophila*, *Amycolatopsis thermoflava*, *Fontimonas thermophila*) were identified, the presence of approximately 100 sequences from *Chloroflexia* was taken to be notable due to their close phylogenetic relationship with the generally thermophilic *Thermomicrobia*. One sequence of particular interest (UniProt ID: A0A2A9HFE7) was identified from *Chloroflexota* bacterium (strain G233). Since this organism grows optimally at a temperature of 75 °C, we tentatively identified the protein encoded by A0A2A9HFE7 as a thermostable BVMO that we dubbed *ssn*BVMO.^25,30^ This sequence was 59% and 41% identical to PAMO and *Ac*CHMO, respectively. Furthermore, the *ssn*BVMO sequence bears two Rossman fold motifs (GxGxx[G/A]) that flank the two consensus sequences associated with BVMOs ([A/G]GxWxxxx[F/Y]P[G/M]xxxD and FxGxxxHxxxW[P/D]; Figs S4 – S5).^31–33^ The presence of these motifs within the sequence of *ssn*BVMO strongly suggested that the identified enzyme would catalyze Baeyer-Villiger oxidations.

### Phylogenetic analysis of ssnBVMO

To further establish the relationship between our target *ssn*BVMO and the previously characterized PAMO and *Ac*CHMO, a phylogenetic tree was constructed. A multiple sequence alignment (MSA) was performed with the sequence of *ssn*BVMO and the sequences found in the cladogram analysis of BVMOs from Furst *et. al*.^5^ This alignment of 45 sequences was then used to construct a maximum likelihood (ML) phylogeny (Fig S6). Using a Class D flavoprotein (Q6Q272) as an outgroup, 12 subgroups were identified based on tree topology and generally high supporting bootstrap values. As expected, *ssn*BVMO clusters with the subgroups containing type I BVMOs. Interestingly, *ssn*BVMO is located in subgroup 2, which also contains PAMO, while *Ac*CHMO is located in the neighbouring subgroup 6.

### *In silico* structural analysis of *ssn*BVMO

The structure of *ssn*BVMO was predicted using AlphaFold (Fig S7)^34^ and compared to the crystal structure of PAMO (PDB ID: 1W4X) using the Needleman-Wunsch alignment algorithm and the BLOSUM-62 similarity matrix (Fig 1).^22^ With a root-mean-square deviation (RMSD) between 434 pruned pairs of 0.970 Å, the overall folds of PAMO and *ssn*BVMO are very similar.

**Figure 1.**
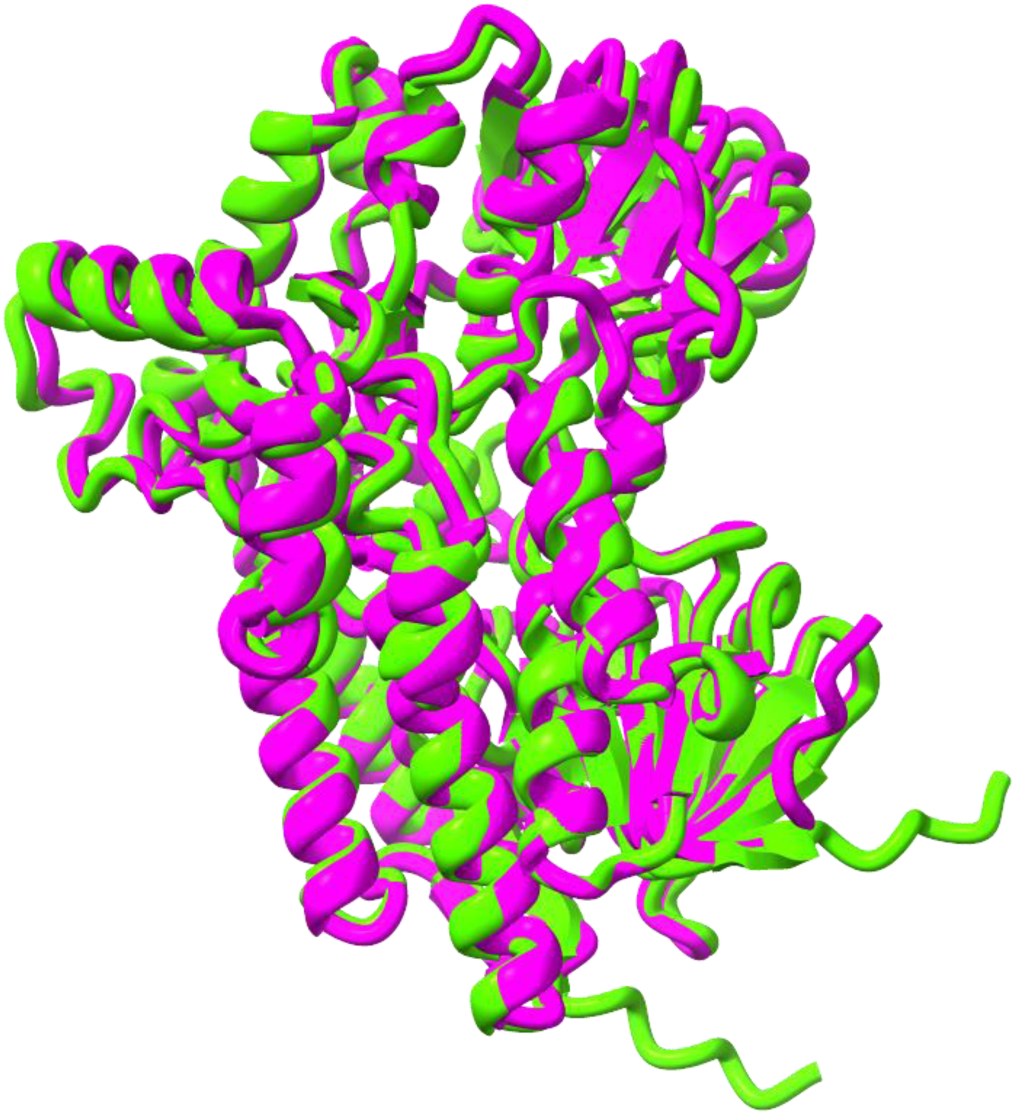
The predicted structure of ssnBVMO (green) overlaid with the experimentally determined structure of PAMO (pink).

The predicted structure of *ssn*BVMO was then aligned with the crystal structure of the Cys65Asp PAMO mutant that has both FAD and NADP^+^ bound in the active site (PDB ID: 4D03).^35^ This alignment suggested tentative binding sites for both FAD and the hydride donor in *ssn*BVMO. Further analysis using an MSA with previously reported BVMOs and CAVER allowed for the tentative assignment of the active site of *ssn*BVMO (Fig 2).^5,36^ Based on extensive mechanistic analysis of existing BVMOs and the structural predictions for *ssn*BVMO, residues likely important for *ssn*BVMO catalysis are Pro434, Ser435, Val436, Leu437, Trp495, Arg330, and Asp589.^5^ It is proposed that the Asp58 modulates the basicity of the active site Arg330 through hydrogen bonding, and that this Arg330 ensures that the C4a-peroxyflavin is kept in the anionic state.^5^ Trp495 is proposed to form a hydrogen bond to the 3ʹ-OH of the NADP^+^ ribose ring without significantly perturbing its electronics, since substitution with other aromatic residues in place of Trp495 is generally tolerated by BVMOs.^5^ Pro434, Ser435, Val436, and Leu437 likely comprise a part of the active site insertion loop, termed the “bulge,” that is thought to be responsible for the proper loading of substrates.^5^ Moreover, CAVER analysis reveals three inner tunnels within *ssn*BVMO: one aligning with the tentative active site residues (pink tunnel), another with the position of FAD (orange tunnel), and a third tunnel aligns with the position of NADP^+^ (cyan) (Fig 2).^36^ It is interesting to note that the proposed substrate tunnel narrows significantly at the “bulge” just before the active site pocket, indicating that dynamic motion of the “bulge” might be needed to accommodate the substrate within the active site.

**Figure 2.**
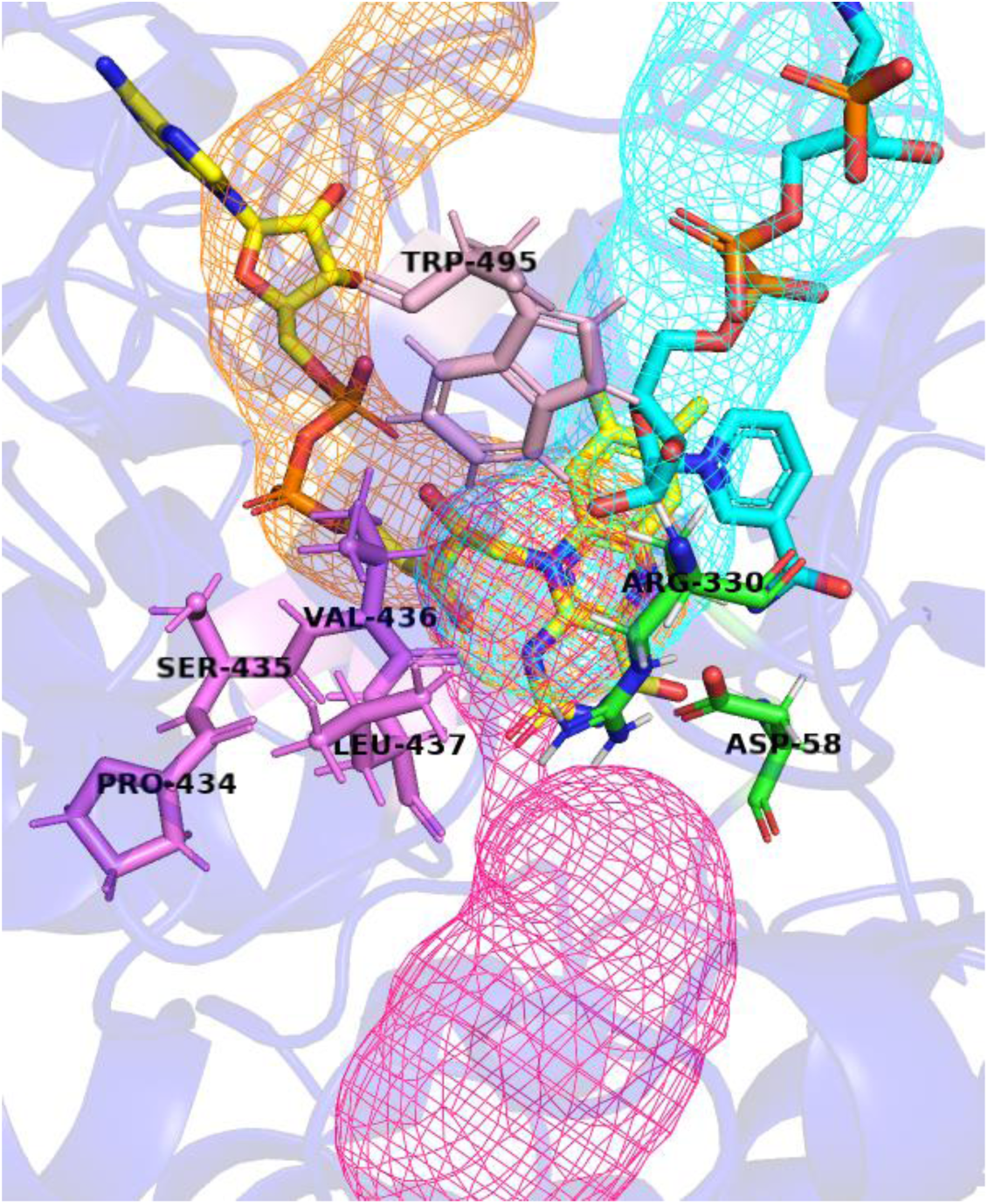
Putative active site residues highlighted based on MSA with known BVMOs and CAVER analysis of the predicted structure of *ssn*BVMO. The positioning of the FAD (yellow) and NADPH (cyan) within the *ssn*BVMO tunnels was determined by structural alignment with the crystal structure of the PAMO Cys65Asp mutant. Important residues for catalysis are labelled and are coloured in green, and the results of the CAVER analysis are shown by mesh tunnels. Tentatively assigned tunnels for FAD, NADPH, and the substrate are shown in orange, cyan, and pink, respectively.

### FAD supplementation and uptake by *ssn*BVMO

Previous work on BVMOs has demonstrated that these proteins are not necessarily purified from *Escherichia coli* in the FAD-bound state.^37^ To bypass possible batch-to-batch variations that this might introduce, BVMOs are commonly supplemented with exogenous FAD following purification from *E. coli*.^38,39^ The uptake of FAD by purified stocks of *ssn*BVMO was clearly demonstrated by comparing absorbance spectra collected immediately after FAD supplementation and approximately 10 minutes after FAD supplementation (Fig S8). Previous experiments have determined that FAD bound to BVMOs has a λ_max_ at 430 nm, while free FAD has a λ_max_ near 450 nm. Thus, the blue shift observed following the incubation of 3 μM of purified *ssn*BVMO with 15 μM FAD can be attributed to FAD uptake by the *apo*-*ssn*BVMO (Fig S8). To further reinforce the importance of FAD for *ssn*BVMO, freshly purified *ssn*BVMO stocks that were subjected to 0.2% SDS treatment demonstrated a clear red shift, with the λ_max_ shifting from 430 nm to 450 nm as the protein denatured and FAD was released (Fig S9). Following this SDS treatment, the molar extinction coefficient of the enzyme-bound FAD was determined to be ε_430_ = 14.2 mM^-1^ cm^-1^. This increase in molar absorptivity agrees with previous studies on the uptake of FAD by BVMOs.^37^

### Determination of the optimal reaction conditions for *ssn*BVMO

To determine the optimal conditions for oxidation by *ssn*BVMO, initial rates of reaction were recorded for the enzyme-catalyzed oxidation of 5 mM 2-octanone with 0.1 mM NADPH at pH 7.5 (100 mM Tris) across a series of different temperatures (Fig S10). As temperature was increased, the initial rates of oxidation increased until 60 °C. At temperatures higher than 60 °C, slow increases in absorbance at 340 nm were observed after a few minutes. This increase in absorbance has been tentatively assigned to precipitation of the enzyme at these elevated temperatures, consistent with the *T*_m_ value observed for *ssn*BVMO (see below). The thermal instability of the NADPH cofactor seemed likely to further confound interpretation of apparent initial rates collected at temperatures ≥ 60 °C. As such, all subsequent kinetic characterization was carried out at 50 °C to minimize any confounding effects associated with the denaturation of the BVMOs or non-enzymatic oxidation of the NADPH cofactor.

A pH-rate profile was then constructed for the BVMO-catalyzed oxidation of 2-octanone at 50 °C. Initial rates of oxidation were measured between pH 5.5 and pH 10.0 using a series of Good’s buffers. The bell-shaped dependence of *k*_cat_ on pH suggested that the protonation states of two ionizable groups (p*K*_a1_ = 6.2 ± 0.2; p*K*_a2_ = 9.0 ± 0.2) are important for catalysis by *ssn*BVMO (Fig S11). Initially, the simplified kinetic scheme presented in Scheme S1 was used to analyze the pH-rate profile with the assumption that enzymatic activity is negligible at both high and low pH (*k*ʹʹ_cat_ = 0 in Scheme S1). While the results of fitting the data to Eqn. S1 with *k*ʹʹ_cat_ = 0 are presented in Table S3 and Figure S11, repeated experiments suggested that enzymatic activity *did not* drop to zero at higher pH values. This suggested a potential alternative in which the Michaelis complexes formed under both neutral (EH•S) and basic (E**^−^**•S) conditions are productive (Scheme S1 with *k*ʹʹ_cat_ ≠ 0). While the pH-rate data seems to be fit more accurately by Eqn. S1 with *k*ʹʹ_cat_ ≠ 0, additional experiments are necessary to confirm or refute the relevance of this reaction pathway to *ssn*BVMO catalysis under basic conditions.

While assignment of the apparent p*K*a values to specific ionizable groups is outside of the scope of this study, the peroxy moiety of the C4a-hydroperoxyflavin intermediate has a reported p*K*a of 8.4 within the context of catalysis by CHMO.^40^ This is similar to the apparent p*K*a^2^ reported here. However, the acidity of this compound can vary widely with the specific monooxygenase under study. Fraaije, for instance, has shown that this same species likely has a p*K*_a_ >> 9 within the active site of PAMO, while Chaiyen has suggested a p*K*_a_ ≈ 10 for an FMN-dependent monooxygenase.^41,42^ While experiments to elucidate the groups responsible for the observed pH dependence are underway, the data in Figure S11 was sufficient to recognize that *ssn*BVMO is maximally active under neutral conditions, such that subsequent kinetic characterization was undertaken at pH = 7.5.

The stability of *ssn*BVMO and was also assayed between pH 5.5 and pH 10.0 by incubating aliquots at different pHs and 4 °C for 24 h. Subsequent measurement of the residual activities of these aliquots using the initial rates of 2-octanone oxidation showed only negligible losses in enzymatic activity following overnight storage of *ssn*BVMO across all tested conditions (Fig S12).

### Determination of the optimal PTDH-BVMO linker peptide

The predicted structure of *ssn*BVMO indicated that both termini of the protein were disordered, such that fusions could be attempted at either site. Initial efforts to fuse 17X-PTDH to the N-terminus of *ssn*BVMO were facilitated by short linker peptides (denoted by the letter L) and constructed using Gibson Assembly. The length and composition of linker peptides has been shown to affect both activity and stability of the resulting fusions. Thus, five different linker sequences (Table S4) were employed to assess the influence of these segments.^38^ To complete this assessment, the pH and temperature-varied activity tests performed on *ssn*BVMO were repeated for each PTDH-L-*ssn*BVMO variant (Fig S13A and S13B). Gratifyingly, all the PTDH-L-*ssn*BVMO fusions operate optimally under very similar conditions to those that are optimal for *ssn*BVMO catalysis. While most fusions behaved nearly identically to the parent *ssn*BVMO, PTDH-L1-*ssn*BVMO was found to remain active at elevated temperatures (>60 °C) that inactivated all the other fusion proteins, suggesting an enhanced thermostability for this fusion. Additionally, PTDH-L1-*ssn*BVMO significantly outperforms all other PTDH-L-*ssn*BVMO fusions in slightly acidic conditions. Following this analysis, PTDH-L1-*ssn*BVMO was selected as the most promising fusion and was taken forward for more detailed characterisation. This characterization included FAD-supplementation assays, determination of optimal reaction conditions for this self-sufficient BVMO, and evaluation of the stability of the fusion at different acidities (Fig S14 and S15).

### Thermal and kinetic stability assays

Using the ThermoFAD method,^43^ *T*_m_ values of 62.5 °C and 60.8 °C were measured for *ssn*BVMO and PTDH-L1-*ssn*BVMO, respectively (Figs 3A, S16 and S17). The effect of ionic strength on the *T_m_* of *ssn*BVMO was also evaluated briefly, and it was found that the addition of 1 M NaCl only slightly increased the *T_m_* to 64.0 °C (Fig S18). A *T*_m_ of 62.5 °C for *ssn*BVMO represents the highest *T*_m_ value of any naturally occurring BVMO, although the archetypically thermostable PAMO exhibits a similar *T*_m_ of 61 °C.^44^ Aside from PAMO, few naturally thermostable BVMOs have been reported. One such BVMO from the eukaryote *Cyanidioschyzon merolae* (*Cm*BVMO) has a reported *T*_m_ of 56 °C, while a cyclohexanone monooxygenase from *Thermocrispum municipale* (*Tm*CHMO) exhibited a *T*_m_ of 52 °C.^45,46^ Similar to *ssn*BVMO, the polycyclic ketone monooxygenase from *Thermothelomyces thermophila* (PockeMO) was also fused to a PTDH. The PockeMO-PTDH fusion and parent PockeMO proteins displayed the same, moderately high *T*_m_ values of 47 °C.^47^ This mirrors the similar *T*_m_ values observed for *ssn*BVMO and PTDH-L1-*ssn*BVMO.

**Figure 3A.**
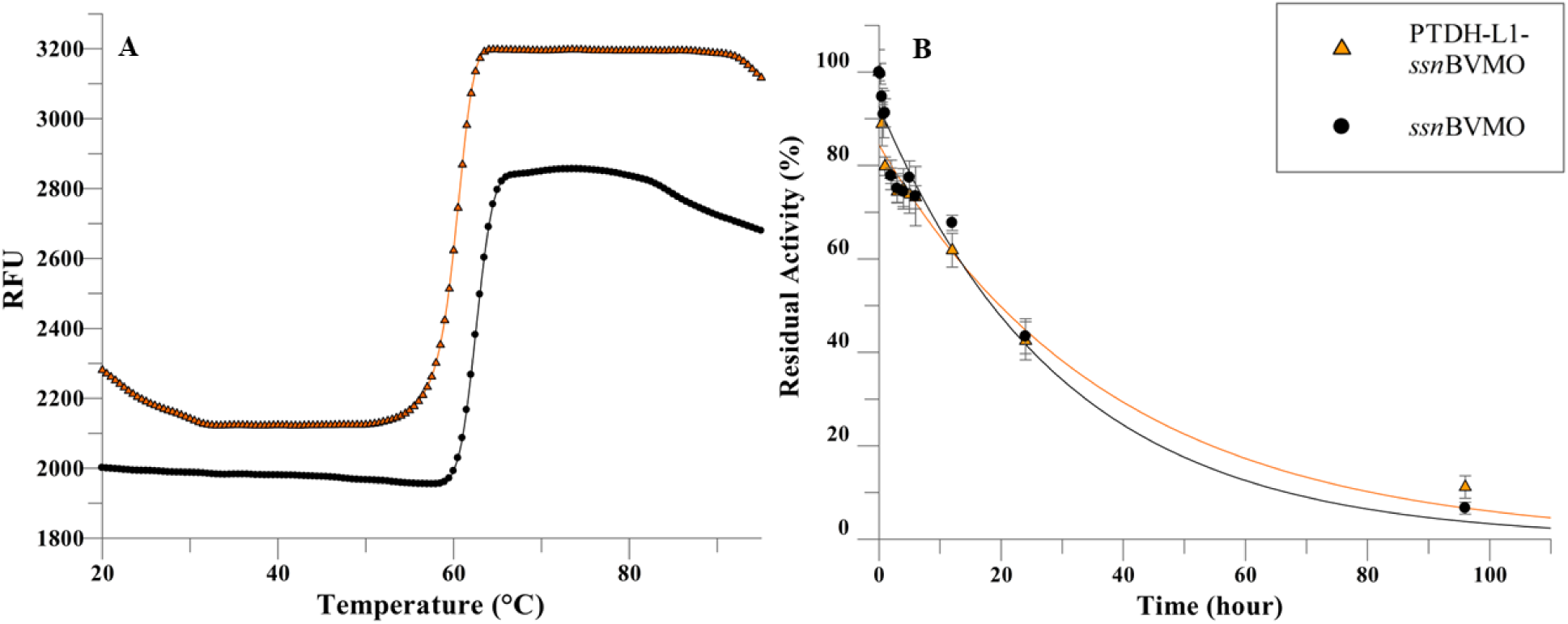
Melt curves for *ssn*BVMO (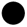) and PTDH-L1-*ssn*BVMO (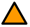) obtained from the relative fluorescence (RFU) of bound and unbound FAD. **B.** Time-dependent loss of activity after incubation of *ssn*BVMO (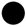) and PTDH-L1-*ssn*BVMO (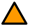) at 50 °C. The residual activity data is fit to Eqn. 1 describing a first-order exponential decay (see methods).

Although a high *T*_m_ is a useful trait for a biocatalyst, the length of time that the enzyme remains active at a given temperature is also a crucial parameter that will dictate the utility of said enzyme. To gauge this kinetic stability of our BVMOs, each enzyme was incubated in a 50 °C water bath, and the residual activity was recorded at various time points. The half-life of thermal inactivation (*t*_1/2_ (50 °C)) of *ssn*BVMO was determined to be *t*_1/2_ (50 °C) = 20 ± 2 h (Fig 3B). The *t*_1/2_ (50 °C) of PTDH-L1-*ssn*BVMO was found to be slightly elevated (*t*_1/2_ (50 °C) = 23 ± 1 h; Fig 3B), although further refinement of these values would be necessary to determine if this difference in *t*_1/2_ (50 °C) is statistically significant. The kinetic stabilities of ssnBVMO and the PTDH-fusion enzymes are particularly significant when recognizing that thermostability exhibited by other BVMOs does not necessarily translate into kinetic stability: *Cm*BVMO is reportedly inactivated by incubation temperatures above 45 °C, whereas the half-life for thermal inactivation (*t*_1/2_) of *Tm*BVMO was measured to be about 9 hr at 30 °C and about 1 – 2 hrs at 40 °C.^45,46^ PockeMO also exhibits limited kinetic stability, with a reported *t*_1/2_ of 24 hrs at 35 °C.^47^

PAMO, the archetypical thermostable BVMO, exhibits both a similar melting temperature (*T*_m_ ∼ 61 °C) and a similar half-life of inactivation at 50 °C (*t*_1/2_ (52 °C) ∼ 24 h) to *ssn*BVMO, and this indicates that the targeted genome mining approach reported here successfully enabled the identification of a novel BVMO that is both thermally and kinetically stable.^48^ Finally, while some BVMOs with high *T*_m_ values (including PAMO) have been reported to become more active after incubation at elevated temperatures, initial rates of reaction with *ssn*BVMO were not affected in this manner. This might indicate that the thermally activated transitions that apparently convert PAMO and other BVMOs to a more catalytically active state are not relevant to *ssn*BVMO catalysis. Along with the high *T*_m_ and *t*_1/2_ (50 °C) values, the finding that *ssn*BVMO does not require any pre-activation is yet another property consistent with a biocatalyst with potential industrial utility.

### Evaluation of the substrate scopes of *ssn*BVMO and PTDH-L1-*ssn*BVMO

Based on the panel of tested substrates using UV-Vis and mass spectrometry, *ssn*BVMO exhibits a marked affinity for linear aliphatic ketones while also processing aromatic ketones with varying patterns of substitution (Table 1). The ability of *ssn*BVMO to process both aromatic and aliphatic ketones is promising, as one of the most significant hurdles to the implementation of PAMO as a biocatalyst is “the limitation of the substrate scope to small aromatic ketones.”^5^ Furthermore, the observation that the alicyclic ketone norcamphor was oxidized by *ssn*BVMO was particularly intriguing, given that PAMO *does not* accept this class of substrate while *Ac*CHMO does.^46^ However, further exploration of alicyclic ketones as potential substrates revealed that these compounds are not generally reactive towards *ssn*BVMO-catalyzed oxidation, with neither estrone nor *β*-tetralone being processed by *ssn*BVMO. While several aromatic ketones were oxidized by *ssn*BVMO, *α*-tetralone was not converted to the corresponding lactone.

**Table 1.** The substrate scope for *ssn*BVMO and PTDH-L1-*ssn*BVMO. GC-MS traces used to confirm product structures are shown in Fig. S20 – S59, and Michaelis-Menten curves used to derive kinetic parameters are shown in Fig. S63 – S81.

| Substrate | Product structure | *ssnBVMO* |  |  | PTDH-L1-*ssnBVMO* |  |  |
| --- | --- | --- | --- | --- | --- | --- | --- |
| | | $k_{\text{cat}}$ (s<sup>-1</sup>) | $K_{\text{M}}$ (mM) | $k_{\text{cat}}/K_{\text{M}}$ (mM<sup>-1</sup>s<sup>-1</sup>) | $k_{\text{cat}}$ (s<sup>-1</sup>) | $K_{\text{M}}$ (mM) | $k_{\text{cat}}/K_{\text{M}}$ (mM<sup>-1</sup>s<sup>-1</sup>) |
| 2-heptanone |  | 0.455 ± 0.038 | 0.0037 ± 0.0015 | 123 ± 51 | 0.61 ± 0.11 | 0.0063 ± 0.0031 | 97 ± 51 |
| 2-octanone |  | 0.671 ± 0.076 | 0.0201 ± 0.0096 | 33 ± 16 | 0.536 ± 0.078 | 0.0257 ± 0.0025 | 21 ± 4 |
| 2-decanone |  | 0.537 ± 0.023 | 0.186 ± 0.057 | 2.88 ± 8.9 | 0.578 ± 0.099 | 0.0112 ± 0.0050 | 52 ± 25 |
| 4-phenyl-2-butanone |  | 0.76 ± 0.10 | 0.022 ± 0.012 | 34 ± 20 | nr<sup>a</sup> | nr<sup>a</sup> | nr<sup>a</sup> |
| propiophenone |  | 0.152 ± 0.003 | 0.373 ± 0.039 | 0.409 ± 0.044 | 0.371 ± 0.057 | 0.70 ± 0.14 | 0.53 ± 0.13 |
| acetophenone |  | 0.335 ± 0.018 | 2.52 ± 0.61 | 0.133 ± 0.033 | 0.459 ± 0.081 | 3.60 ± 0.79 | 0.127 ± 0.036 |
| norcamphor |  | 0.165 ± 0.006 | 25.7 ± 2.4 | 0.00639 ± 0.00063 | 0.096 ± 0.021 | 6.0 ± 1.9 | 0.0160 ± 0.0062 |
| progesterone<sup>b</sup> |  | 0.537 ± 0.058 | 0.274 ± 0.082 | 1.96 ± 0.62 | 0.312 ± 0.051 | 0.033 ± 0.011 | 9.4 ± 3.4 |
| 4-hydroxyacetophenone |  | 0.455 ± 0.026 | 1.28 ± 0.18 | 0.356 ± 0.055 | 0.301 ± 0.061 | 0.76 ± 0.12 | 0.39 ± 0.10 |
| 3-hydroxyacetophenone |  | 0.127 ± 0.038 | 2.5 ± 2.0 | 0.051 ± 0.043 | 0.120 ± 0.023 | 1.59 ± 0.27 | 0.075 ± 0.019 |

<sup>a</sup>nr = no reaction; <sup>b</sup>Product structure was determined by UPLC-MS and <sup>1</sup>H NMR (Fig. S60 – 62).

Like its parent enzyme, PTDH-L1-*ssn*BVMO displays a preference for linear ketones over bulkier aromatic ketones. Interestingly, the most efficiently processed aromatic ketone by *ssn*BVMO, 4-phenyl-2-butanone, is not processed by PTDH-L1-*ssn*BVMO. The bulkiness of 4-phenyl-2-butanone does not seem to be the limiting factor, as the oxidation of progesterone by the fusion enzyme was accompanied by a ∼380% increase in catalytic efficiency compared to *ssn*BVMO, giving it a comparable efficiency to that exhibited with linear ketones. Notably, there is also a significant increase in the catalytic efficiency for 2-decanone, stemming primarily from an apparent increased binding affinity for the substrate. Previous BVMO-PTDH fusions produced by Fürst^47^ and Torres Pazmiño^26^ showed similar, yet less pronounced increases in catalytic efficiency with model substrates compared to the parent BVMO. As seen for PTDH-L1*-ssn*BVMO, most of these increases in catalytic efficiency stem from increases in the apparent binding affinities. Efforts to crystallize both *ssn*BVMO and PTDH-L1-*ssn*BVMO are underway, with hopes to examine possible conformational changes induced by the fusion to 17X-PTDH. This structural information may permit a more precise explanation of the observed differences in substrate scope and catalytic efficiencies of *ssn*BVMO and PTDH-L1-*ssn*BVMO.

Following determination of the substrate scope for *ssn*BVMO catalysis, site directed mutagenesis was used to evaluate the identity of the putative active site. Specifically, the R330A mutant of *ssn*BVMO was generated, purified, and assayed against 2-heptanone at 50 °C and pH 7.5. While the wildtype enzyme was found to catalyze the oxidation of this compound, the observed inactivity of R330A-*ssn*BVMO with 2-heptanone demonstrates the importance of Arg330 for oxidation by *ssn*BVMO (Fig S19).

Several BVMOs catalyze “non-canonical” oxidations of functionalities other than ketones. Depending on the biocatalyst used and the regiochemistry of the oxidation, aldehydes can be converted into the corresponding carboxylates or the formate esters that can decompose to the corresponding alcohol. However, *ssn*BVMO and PTDH-L1-*ssn*BVMO exhibited no activity toward benzaldehyde. Similarly, neither BVMO was able to catalyze the non-canonical oxidation of *N*,*N*-dimethylaniline to the corresponding *N*-oxide nor of thioanisole into the corresponding sulfoxide/sulfone.

Intriguingly, *ssn*BVMO was also able to regiospecifically oxidize progesterone, with only the exocyclic ketone being converted to the corresponding ester. This suggests *ssn*BVMO may exhibit intermediate levels of promiscuity with steroidal substrates when compared with PAMO that does not accept these compounds and PockeMO that is able to oxidize both exocyclic ketones and alicyclic ketones found within either the A ring or the D ring of steroidal substrates. This also suggests that *ssn*BVMO is more tolerant of bulky substrates than the archetypically promiscuous *Ac*CHMO, as the latter enzyme is unable to process progesterone.^46^ PTDH-L1-*ssn*BVMO was also confirmed to oxidize progesterone through the use of UPLC-MS.

Collectively, these results suggest that *ssn*BVMO exhibits a broader substrate scope than PAMO, but a somewhat narrower scope than *Ac*CHMO. To the best of our knowledge, *ssn*BVMO is the most kinetically and thermally stable of the naturally occurring BVMOs known to date. Given this stability and the ability of this enzyme to catalyze the oxidation of linear aliphatic ketones, aromatic ketones, a cyclic ketone, and sterically bulky substrates, *ssn*BVMO has significant potential to enable industrial enzymatic oxidations. While *ssn*BVMO itself is less catalytically efficient than several highly engineered BVMOs, this novel enzyme represents a stable, promiscuous biocatalyst that would serve as a natural starting point for the directed evolution of increasingly efficient variants of *ssn*BVMO. While the few differences between the substrate scopes of *ssn*BVMO and the fusion protein are intriguing, results with PTDH-L1-*ssn*BVMO largely indicate that fusion of PTDH to *ssn*BVMO does not disrupt the ability of the BVMO to oxidize a wide range of substrates.

### Flexible docking studies

Computational analysis using the AutoDock suite^49^ demonstrated that substrate orientation, rather than binding energy, is crucial for catalysis by *ssn*BVMO. Specifically, carbonyl groups of compounds that were not processed by *ssn*BVMO did not hydrogen bond with the Arg330 residue that is essential for catalysis.^5^ This observation holds regardless of the apparent binding energy (see Fig 4). Notably, there was no significant difference in the lowest binding energies between compounds that were oxidized and those that were not, suggesting that factors other than binding energy influence the oxidation of ketones by *ssn*BVMO. Exceptions to this trend were observed with progesterone and cyclohexanone. The lowest energy binding orientation of progesterone did orient the substrate carbonyl group towards Arg330, but the distance between the hydrogen bond donor and acceptor was larger than with other processed substrates (4.2 Å). Conversely, cyclohexanone was bound in the proper orientation for hydrogen bond formation with Arg330, and yet showed no conversion. However, in a preliminary directed evolution campaign, a single round of error-prone PCR resulted in a variant *ssn*BVMO that was able to oxidize cyclohexanone (data not shown). This might suggest an inherent predisposition of *ssn*BVMO to process this substrate, consistent with the suggested H-bond with Arg330.

**Figure 4A.**
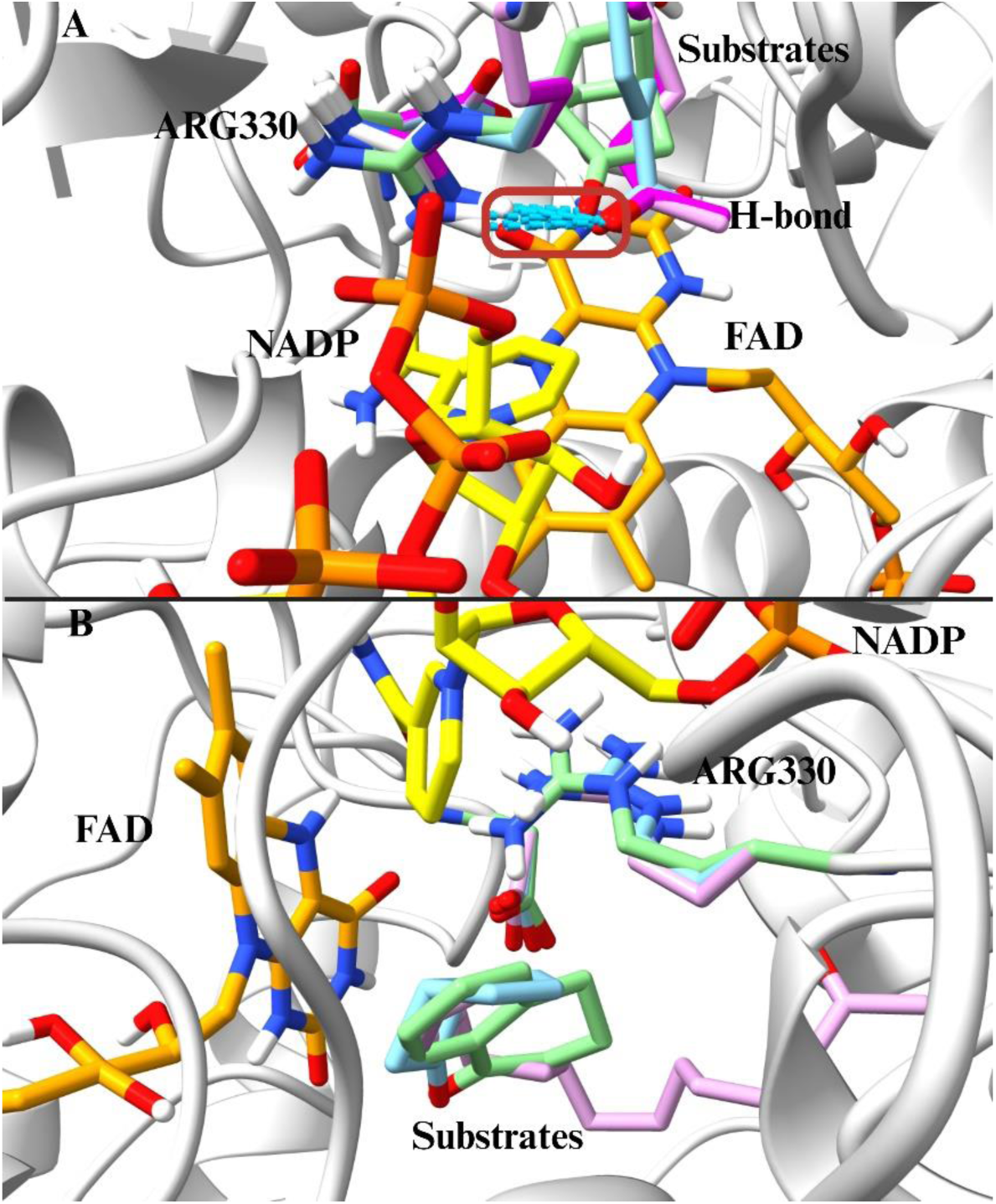
For substrates oxidized by *ssn*BVMO, including norcamphor (pale green), 3-hydroxyacetophenone (light blue), 2-heptanone (pink), and 4-phenyl-2-butanone (magenta), the lowest energy binding orientation involves forming a hydrogen bond with Arg330. **B.** For substrates that *ssn*BVMO did not process, such as α-tetralone (pale green), (±)-*cis*-bicyclo[3.2.0]hept-2-en-6-one (light blue), and 2-pentadecanone (pink), the lowest energy binding orientation was identified. During docking, Asp58 and Arg330 were set as flexible residues, and these residues are colored to match their corresponding substrate.

### Conversion assays using propiophenone

Although several engineered BVMOs are available that catalyze BV reactions more efficiently, we opted to evaluate the potential of *ssn*BVMO as an oxidative biocatalyst by quantifying the conversion of propiophenone to phenyl propanoate (Fig S82). These quantitative conversion experiments employed propiophenone as a substrate because the aromatic substrates and products could be extracted reproducibly via a simple solid phase extraction (SPE) that permitted accurate conversion determinations. Initial qualitative GC-MS analyses with several different substrates illustrated that the conditions employed for the *ssn*BVMO-catalyzed oxidations also led to hydrolysis of the expected esters into alcohols. Consequently, both phenyl propanoate and phenol were produced in the *ssn*BVMO-catalyzed oxidation of propiophenone. To quantify conversion, calibration curves were constructed with both products and propiophenone using toluene as an internal standard. After incubating propiophenone with *ssn*BVMO for 48 hrs at 35 °C, the phenyl propanoate peaks present in the chromatograms were not significantly greater than the noise associated with the instrument (Fig S82). As such, the final conversion of propiophenone to phenyl propionate was evaluated using the phenol peak integral and was determined to be 95 ± 1.8% with a turnover number (TON) of 179. We note that the reaction conditions employed were not optimized for conversion, so the observation that 95% of a 3 mM propiophenone stock could be turned over by *ssn*BVMO in 48 hrs serves as an illustration of the potential utility of this enzyme even before engineering efforts to enhance the catalytic efficiency of this enzyme. An analogous experiment evaluating the oxidation of propiophenone by PTDH-L1-*ssn*BVMO has yielded an increased TON of 299 when tested under similar conditions, which is consistent with the increased catalytic efficiency observed in Table 1 (Fig S83). Moreover, conversion experiments with PTDH-L1-*ssn*BVMO also demonstrate that the fusion of PTDH was successful since we have achieved a 62% conversion of 3 mM propiophenone using 0.25 mM of NADPH. It is important to acknowledge that variability in SPE extraction efficiencies of different compounds across different days likely renders comparisons of conversions measured without the use of internal standards misleading. Consequently, we exercise caution and abstain from making such comparisons.

### Conclusions & Future Work

In this study, targeted genome mining was used to find gene sequences sourced from thermophilic microorganisms that had high sequence similarity to promiscuous BVMOs. The resultant enzyme was found to be both thermostable and relatively substrate promiscuous. Given these properties, as well as the above-average kinetic stability of *ssn*BVMO, this enzyme has the potential to be a useful starting point for engineering a promiscuous and stable oxidative biocatalyst. Even before efforts to optimize the catalyst or reaction conditions, the PTDH-L1-*ssn*BVMO fusion protein was designed and found to be able to catalyze the near-complete conversion of propiophenone with sub-stoichiometric amounts of NADPH, further demonstrating that *ssn*BVMO is a good candidate for enzyme engineering. Moreover, combined with preliminary docking studies performed here, further investigation of the active site of *ssn*BVMO through additional computational and mechanistic studies may enhance our understanding of what has the potential to be a new class of naturally occurring BVMOs that are both substrate promiscuous and thermostable, challenging the classic paradigm that these enzymes are either substrate promiscuous or thermostable.

## Materials & Methodology

### Chemicals and materials

All compounds and solvent, unless otherwise indicated, were purchased from MilliporeSigma. All buffers used were purchased from BioBasic and used as received. Reduced nicotinamide adenine dinucleotide phosphate (NADPH), sodium chloride, Lysogeny Broth (LB), glycerol, Na_2_HPO_4_, KH_2_PO_4_, tryptone, yeast extract, glucose, and lactose were purchased from BioShop Canada.

### Bioinformatics

All sequence similarity networks (SSNs) were constructed using the Enzyme Function Initiative’s Enzyme Similarity Tool (EFI-EST) and were analyzed using Cytoscape 3.8.0. Protein family (PFam) assignments were done using the InterProScan webtool.^50–52^ *ssn*BVMO (A0A2A9HFE7), a Class D flavoprotein (Q6Q272), and 46 sequences found in the cladogram analysis from Fürst *et al.* were downloaded from Uniprot and aligned in AliView 1.1.^5,53^ Subsequently, a maximum-likelihood phylogenetic tree was estimated by RAxML version 8.2.12 using 47 sequences as inputs on the CIPRES gateway with automatic bootstrapping terminating at 1000 bootstrap replicates.^54,55^ The resulting phylogeny was visualized with FigTree.

### Gibson Assembly of pET-28a(+)-PTDH-L1-*ssn*BVMO

Six variations of the PTDH-L1-*ssn*BVMO plasmid were produced, differing solely by a short linker sequence separating the BVMO and 17X-PTDH regions. These plasmids were prepared using Gibson assembly, using pET-28a-*ssn*BVMO as the backbone, and 17X-PTDH as the insert. The varying linker sequences (Table S4) were introduced to each assembly by encoding them into the primers. The 17X-PTDH gene was annealed to the 5ʹ end of the *ssn*BVMO gene using Gibson assembly, with the linker sequence separating the two genes. Assembly was completed according to the New England Biolabs’ protocol. Assembled plasmids were transformed into *E. coli* DH10β using heat shock, and the resulting transformants were used to inoculate 5 mL overnight cultures. Plasmids were extracted from these cultures following the protocol outlined in the Monarch plasmid Miniprep kit. Successful assembly was determined through whole plasmid sequencing by Plasmidsaurus.

### Protein overexpression and purification

#### *ssn*BVMO

The N-terminally His_6_-tagged gene encoding *ssn*BVMO was codon optimized for expression in *E. coli*, synthesized, and cloned into a pET-28a (+) expression vector by BioBasic. Transformation of this plasmid into chemically competent *E. coli* BL21(DE3) was achieved *via* heat shock. A single colony was then used to inoculate an overnight culture. This starter culture was used to inoculate 1 L of LB media supplemented with 50 µg/mL kanamycin, which was subsequently grown to an OD_600_ of 0.6 before supplementation with 0.4 mM isopropyl *β*-D-1-thiogalactopyranoside (IPTG). The culture was then grown at 18 °C and 180 rotations per minute (RPM) for approximately 18 h. Cells were collected by centrifugation 4000 x *g* (10 min) and resuspended in lysis buffer (50 mM Tris, pH 7.50, 0.5 M NaCl, 10% (v/v) glycerol). Cells were then lysed by multiple passes through an Emulsiflex at ∼10,000 psi. Cellular debris were removed by centrifugation at 16,000 x *g* for 1 h, and the supernatant loaded onto a Ni^2+^-NTA agarose resin (Cytiva) that had been pre-equilibrated with lysis buffer. The column was washed with approximately 10 column volumes of wash buffer (50 mM Tris, pH 7.5, 0.1 M NaCl, 10 mM imidazole, 10% (v/v) glycerol) before the protein of interest was eluted using 50 mM Tris, pH 7.5, 0.1 M NaCl, 250 mM imidazole, 10% (v/v) glycerol. The collected protein was then buffer exchanged into storage buffer (50 mM Tris, pH 7.5, 0.1 M NaCl, 20% v/v glycerol). 10 mM flavin adenine dinucleotide (FAD) was then added, and the FAD-supplemented enzyme was kept at 4°C overnight before being flash-frozen and stored at -80 °C. Enzyme concentrations were determined using Bradford assays, and SDS-PAGE was used to evaluate the purity of the *ssn*BVMO (Fig S84). Typical purification yields were 75 – 105 mg from 1 L cultures.

#### PTDH-L-*ssn*BVMO

Each assembled pET-28a(+)-PTDH-L-*ssn*BVMO plasmid was separately transformed into *E. coli* BL21(DE3), and inoculated into starter cultures as was described above for *ssn*BVMO. Starter cultures were used to inoculate 1 L of autoinduction media (6 g/L Na_2_HPO_4_, 3 g/L KH_2_PO_4_, 20 g/L tryptone, 5 g/L yeast extract, 5 g/L NaCl, 6 mL/L glycerol, 0.5 g/L glucose, 2 g/L lactose), which was grown for approximately 18 h at 37 °C and 180 RPM. Cell lysis, purification, buffer exchange, FAD supplementation, concentration assays and SDS-PAGE were all completed as described in the above section (Fig S85-S88). Typical purification yields were 32-72 mg from 1 L cultures. It is important to note that preliminary activity tests with PTDH-L-*ssn*BVMO variants were performed without exogenous FAD supplementation.

### FAD uptake tests

For FAD uptake tests, 3 µM of *ssn*BVMO that had not been supplemented with FAD following purification by immobilized metal affinity chromatography (IMAC) was incubated with 15 µM FAD and spectral scans between 300 to 600 nm were taken over time to monitor for exogenous FAD uptake by the enzyme.

To determine the molar extinction coefficient of *ssn*BVMO-bound FAD, the absorbance of purified enzyme in 100 mM Tris, pH 7.5 was recorded between 300 nm and 600 nm. The enzyme was then denatured by the addition of 0.2% sodium dodecyl sulfate (SDS) to release the FAD, and the absorbance spectrum of this mixture was recorded over time until the observed λ_max_ remained constant with time. Together with the known extinction coefficient of FAD at 450 nm (ε_450_ = 11.3 mM^-1^ cm^-1^), the absorbance of this mixture at this wavelength was then used to determine [FAD]_released_. The concentration of FAD released upon SDS treatment was assumed to be equal to the concentration of catalytically active *ssn*BVMO. The absorbance spectrum of *ssn*BVMO in buffer was also recorded in triplicates and used to determine the molar extinction coefficient of *ssn*BVMO-bound FAD.

### Melting point determinations

As *ssn*BVMO is a flavoprotein, the melting temperature (*T*_m_) at which half of *ssn*BVMO or PTDH-L1-*ssn*BVMO is unfolded can be determined using the differential fluorescence of the bound and unbound FAD.^43^ 5 µL worth of BVMO stocks were diluted with 15 µL of 100 mM Tris buffer at pH 7.5 (final concentration of both BVMOs was 0.019 mM), dispensed into a 96-well plate and placed into a BioRad CFX-Connect RT-PCR System with fluorescence measurements set to the Förster Resonance Energy Transfer (FRET) setting. The typical excitation wavelengths for RT-PCRs (450-530nm) corresponds well with the flavin excitation maxima at 373-375 and 445-450 nm.^43^ The instrument was set to ramp from 20.0 °C to 95.0 °C at 0.5 °C increments, with each temperature being held for 10 s before the fluorescence was recorded. All measurements were performed in triplicate. For *ssn*BVMO, identical experiments were conducted in the presence of varying concentrations of sodium chloride (NaCl) to determine the influence of ionic strength on its thermostability.

### Steady-state kinetic assays

Typical kinetic assays were performed using 0.5 – 1 µM *ssn*BVMO or PTDH-L1-*ssn*BVMO, 0.1 mM NADPH, 100 mM Tris at pH 7.5, and variable concentrations of the potential substrate. All substrate stocks were prepared in *p*-dioxane, with the final assay concentration of *p*-dioxane at 2% (v/v). Initial rates of NADPH oxidation were recorded for at least 60 seconds at 340 nm (ε_340_ = 6.22 mM^-1^cm^-1^) using an Agilent Cary 3500. As 3-hydroxyacetophenone and 4-hydroxyacetophenone absorb strongly at 340 nm, kinetic measurements with these substrates were carried out using the absorbance of NADPH at 370 nm (ε_370_ = 2.7 mM^-1^cm^-1^).^56^ Background rates of spontaneous NADPH oxidation was measured by monitoring the absorbance of a mixture of the BVMO and NADPH prior to the initiation of the reaction by addition of the substrate. Substrate stocks were incubated in a water bath set to the same temperature as the enzyme-catalyzed reaction was run to minimize potential temperature fluctuations upon substrate addition. All reactions were performed in triplicate.

### Evaluating the influence of pH and temperature on BVMO catalysis

To determine the optimal temperature for both *ssn*BVMO and PTDH-L1-*ssn*BVMO catalysis, reactions were run between 35 °C and 65 °C. Each of these experiments was conducted using 5 mM 2-octanone and 0.1 mM NADPH at pH 7.5 in 100 mM Tris.

pH-rate profiles were constructed for the BVMO-catalyzed oxidations of 5 mM 2-octanone at 50 °C in 100 mM Tris, pH 7.5. Initial rates of reaction were measured between pH 5.5 and pH 10.0 using a series of Good’s buffers (pH 5.5 – 6.5, 100 mM MES; pH 7.0 – 8.5, 100 mM Tris; pH 9.0 – 10.0, 100 mM CHES). The stability of *ssn*BVMO in different acidities was also assayed by measuring the residual activity of aliquots that were stored for 24 h at 4 °C.

The kinetic stability of the BVMOs were assayed by incubating several 2 μM enzyme aliquots from a given enzyme preparation at 50 °C. At various time points, aliquots were removed from the water bath and placed on ice for 5 – 10 minutes. Samples were then centrifuged briefly to remove any precipitate, and an equal volume of supernatant was added to a cuvette held at 50 °C that contained 0.1 mM NADPH in 100 mM Tris, pH 7.5. The background rate of NADPH oxidation was monitored for approximately 5 minutes before the reaction was initiated by the addition of 5 mM 2-octanone. The apparent first-order rate constants for the thermal inactivation of *ssn*BVMO and PTDH-L1-*ssn*BVMO (*k*_i_) were determined by fitting the residual activity data to Eqn. 1. The half-life of thermal inactivation at 50 °C (*t*_1/2_ (50 °C)) was then obtained using the relationship given in Eqn. 2.

### Product characterization using GC-MS

The products of the BVMO-catalyzed oxidations were analyzed by gas chromatography-mass spectroscopy (GC-MS). For *ssn*BVMO-catalyzed oxidations, reactions were carried out in 1 mL of 100 mM Tris-Base buffer at pH 7.5 with 5 µM of *ssn*BVMO, 5 µM of the thermostable phosphite dehydrogenase (17X-PTDH), 250 µM NADPH, 100 mM sodium phosphite, and 25 mM substrate.^57,58^ For PTDH-L1-*ssn*BVMO-catalyzed oxidations, 17X-PTDH was not used, but all other conditions and reagents remained the same. Due to the limited solubility of 4-hydroxyacetophenone and progesterone, substrate concentrations for the oxidation of these compounds were 5 mM and 0.8mM, respectively. The reactions were left for 24 h at 35 °C and 180 RPM before the mixture was loaded onto a HyperSep solid phase extraction (SPE) cartridge (500 mg/2.8 mL; Thermo Scientific). Following a water wash and subsequent gentle drying under vacuum, 1 mL of ethyl acetate was used to elute the organic compounds. Extracted samples were then analyzed using a Thermo Scientific iSQ7000 GC-MS with a Phenomenex ZS-5ms capillary column (30 m, 0.25 mm I.D.). Oven settings and related parameters are available in Table S5. Negative controls were run and processed identically, with the exception that an equal volume of buffer was added to the sample in place of BVMO. Identical conditions were employed to characterize the products of PTDH-L1-*ssn*BVMO-catalyzed oxidations.

### Progesterone conversion assay using UPLC-MS

Due to the low volatility of progesterone, our typical GC separation was replaced by ultra-high pressure liquid chromatography (UPLC) with a diode array detector (DAD) and mass spectrometer (MS). Reaction conditions for both PTDH-L1-*ssn*BVMO and *ssn*BVMO remained the same. Reactions were prepared for UPLC analysis by adding 1:1 (v/v) acetonitrile and subsequent centrifugation for 5 min at 13 000 x *g.* Following centrifugation, reactions were passed through a 0.22 µm filter. UPLC-DAD-MS analysis was completed using a C18 column (CORTECS T3 column 1.6 μm particle size, 2.1×100 mm) with a CORTECS VanGuard pre-column. Elution conditions are available in Table S6. Negative controls were run and processed identically, with the exception that an equal volume of buffer was added to the sample in place of BVMO.

### Large scale progesterone reaction for NMR analysis

The limited solubility of progesterone necessitated larger-scale reactions to determine the product of the BVMO-catalyzed oxidations. To this end, reactions were run in 50 mL of 100 mM Tris buffer at pH 7.5 with 2 µM of BVMO, 2.6 µM 17X-PTDH, 100 µM NADPH, 60 mM sodium phosphite, and 1 mM progesterone. The reactions were incubated at 35 °C and 180 RPM for 7 days with 24-hour supplementations of BVMO (final concentration after 7 days, ∼7 µM) and 17X-PTDH (final concentration for *ssn*BVMO reaction after 7 days, ∼18 µM). After one week, *p*-dioxane was removed by rotary evaporation. The samples were then lyophilized to dryness before being re-dissolved in CDCl_3_ and analyzed using a Bruker 400 MHz UltraShield^TM^ NMR Instrument. This large-scale experiment was only done with *ssn*BVMO, with the assumption the fusion of 17X-PTDH onto *ssn*BVMO would not alter regiospecificity of the parent enzyme.

### Quantitative substrate conversion experiments

For quantitative conversion experiments, BVMO-catalyzed oxidations of propiophenone were carried out in 1 mL of 100 mM Tris buffer at pH 7.5. These reactions included 8 µM BVMO, 9 µM 17X-PTDH (only for *ssn*BVMO-catalyzed reactions), 250 µM NADPH, 100 mM sodium phosphite, and 3 mM propiophenone. Negative controls were run with an equal volume of buffer added in place of the BVMO. Reactions were incubated for 48 h at 35 °C and 180 RPM. After 24 h, additional portions of the BVMO (to a final concentration of ∼16 µM) and 17X-PTDH (to a final concentration of ∼18 µM, only for *ssn*BVMO reaction) were added to the reaction mixture. Products were extracted using SPE as described above, and toluene was then added as an internal standard at a final concentration of 20 mM.

Separate calibration solutions of propiophenone, phenol, and phenyl propionate were made at 0.05 mM, 0.1 mM, 0.2 mM, 0.3 mM, 0.4 mM, 0.5 mM, 1 mM, 2 mM, and 3 mM concentrations. In each of these solutions, toluene was held fixed at 20 mM as an internal standard. Calibration curves were constructed for propiophenone, phenol, and phenyl propionate, and the ratio of the known concentrations of each analyte to that of toluene was plotted against the ratio of the peak area for each analyte to that of toluene. The slopes of these plots serve as correction factors that permit conversion of peak areas in the GC-MS chromatograph (measured as described above) into concentrations.

### Computational analysis

Protein and ligands were setup according to AutoDock4 docking protocol outlined by Forli *et. al* and were docked using AutoDockFR for easier flexible residue docking.^59,60^ NADP^+^ and FAD were docked first, utilizing the crystal structure of PAMO (PDB ID: 4D03) as a reference for selecting structures where FAD and NADP^+^ are correctly oriented in relation to the active site residues.^35^ After FAD and NADP^+^ were successfully docked, substrate docking was performed with ASP58 and ARG330 set as flexible residues to allow for correct binding conformation.

## ASSOCIATED CONTENT

### Supporting Information

A listing of the contents of each file supplied as Supporting Information should be included.

## AUTHOR INFORMATION

### Author Contributions

The manuscript was written through contributions of all authors. All authors have given approval to the final version of the manuscript. †These authors contributed equally.

### Funding Sources

This work was supported in part by a Discovery Grant (RGPIN-2020-04455) from the Natural Sciences and Engineering Council (NSERC).

### Notes

The authors declare no competing financial interests.

## ACKNOWLEDGMENT

We thank Dr. Igor Kozin for assistance with GC-MS measurements. SED and EH thank NSERC for financial support through undergraduate student research awards and postgraduate scholarships.

## Notes

### Competing Interest Statement

The authors have declared no competing interest.

