## Supplementary Information for "Genome mining leads to the identification of a stable and promiscuous Baeyer-Villiger monooxygenase from a thermophilic microorganism"

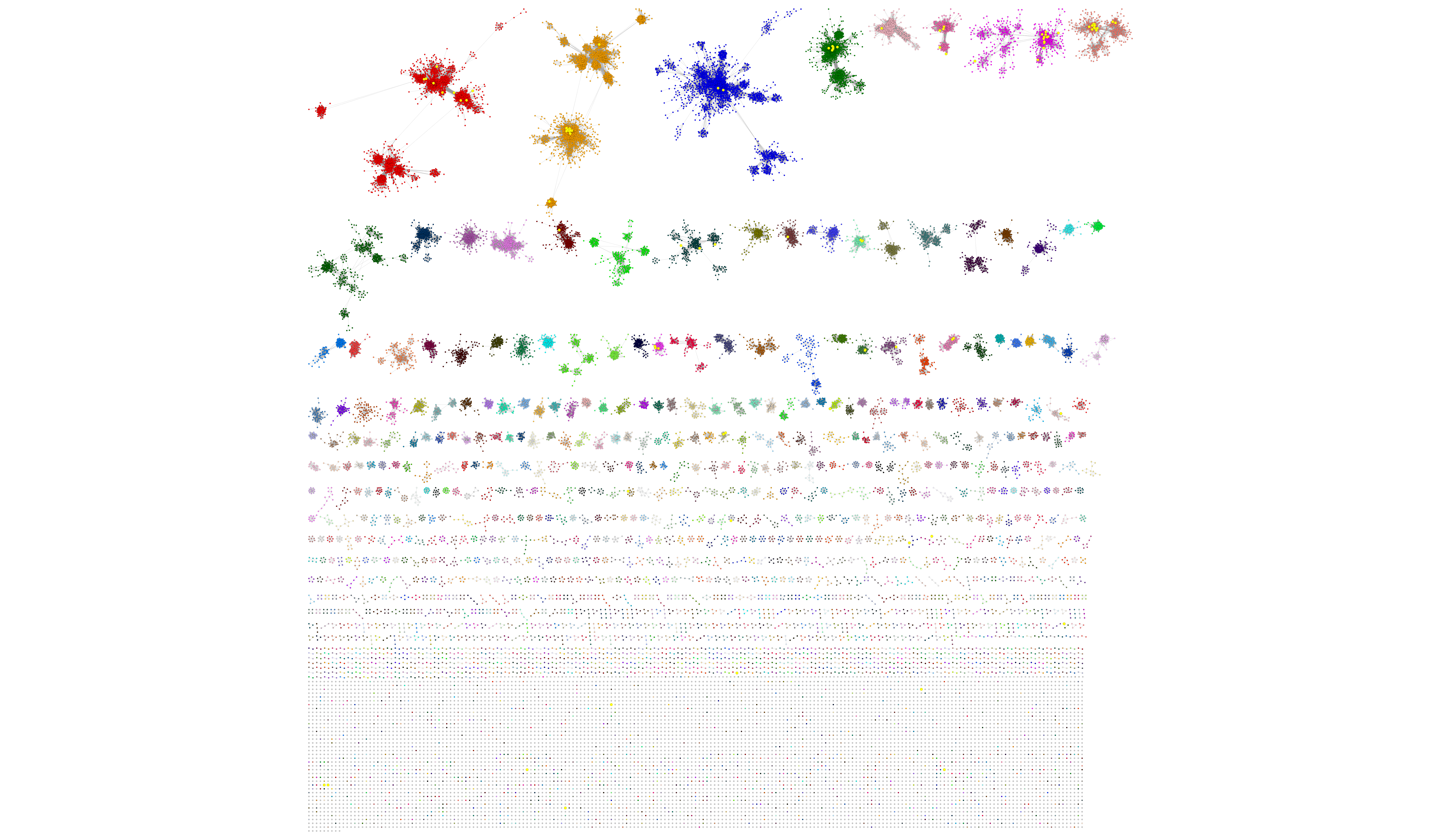

**Figure S1.** The RepNode 60 SSN for the combined PFams 00743 and 13738 (UniRef90) at an AS of 140. Sequences with experimentally verified activities are represented with yellow nodes. Sequences encoding PAMO and *Ac*CHMO are both found within the largest cluster (shown in red).

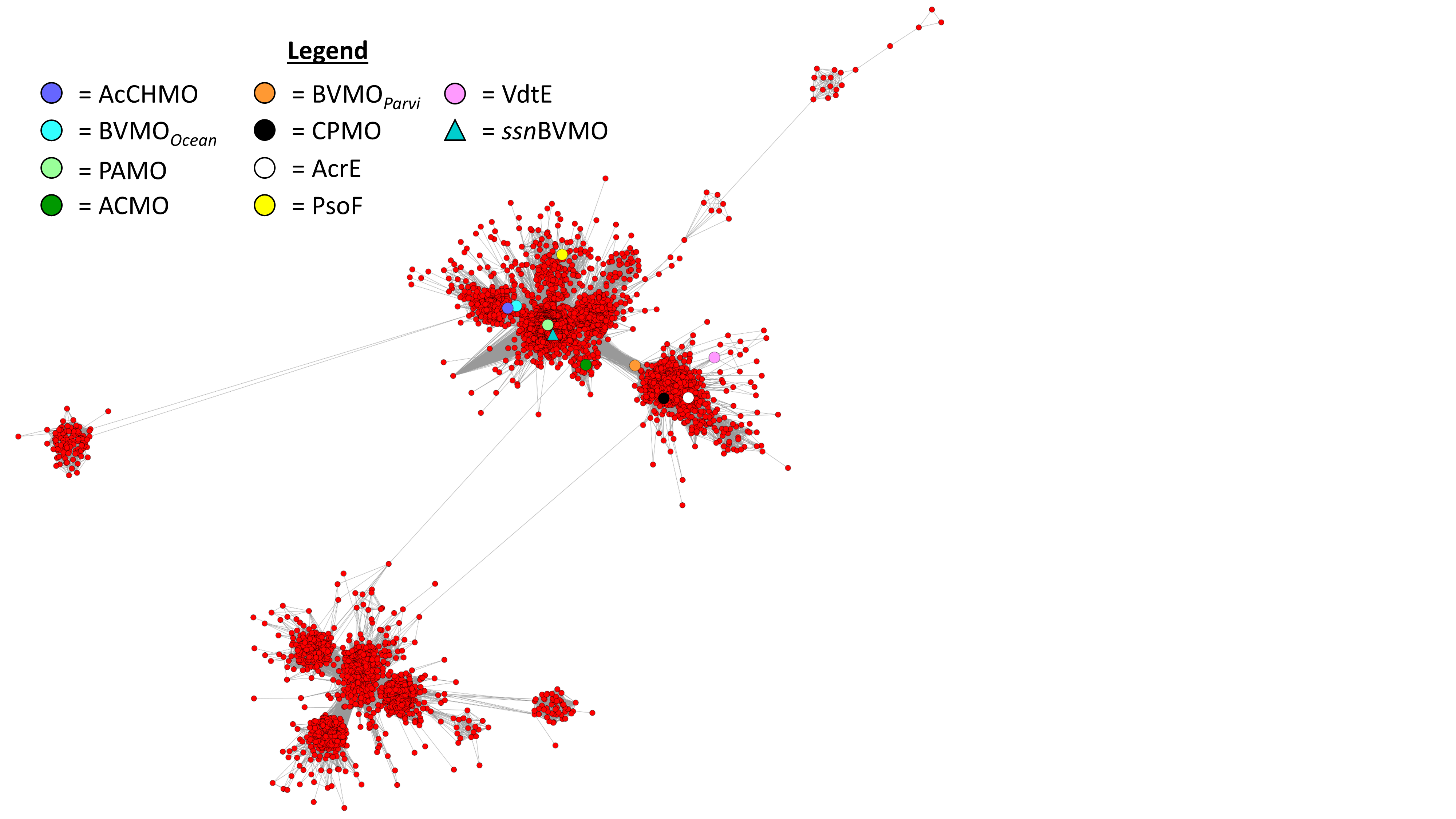

**Figure S2.** A daughter network constructed from the largest cluster in the SSN shown in Figure S2. Sequences encoding enzymes with known functions are represented with differently coloured shapes: Cyclohexanone monooxygenase (*Ac*CHMO; ); Baeyer-Villiger monooxygenase (BVMO*_Ocean_*; ); phenylacetone monooxygenase (PAMO; ); acetone monooxygenase (ACMO; ••); Baeyer-Villiger monooxygenase (BVMO*_Parvi_*; ); cyclopentanone monooxygenase (CPMO; ••); FAD-binding monooxygenase (AcrE; ); dual-functional monooxygenase/methyltransferase (PsoF; ); FAD-binding monooxygenase (VdtE; ). The sequence encoding *ssn*BVMO was identified within this network (*ssn*BVMO; ).

**Table S1.** Sequence information for entries in Figure S2 with known function.

| Entry | UniProt code | Source organism | Reference |
| --- | --- | --- | --- |
| *Ac*CHMO | P12015 | *Acinetobacter* sp. | (1) |
| BVMO*_Ocean_* | A3U3H1 | *Oceanicola batsensis* | (2) |
| PAMO | Q47PU3 | *Thermobifida fusca* (strain YX) | (3) |
| ACMO | A1IHE6 | *Gordonia* sp. (strain TY-5) | (4) |
| BVMO*_Parvi_* | A7HU16 | *Parvibaculum lavamentivorans* (strain DS-1) | (2) |
| CPMO | Q8GAW0 | *Comamonas* sp. (strain NCIMB 9872) | (5) |
| AcrE | A0A1L9WQQ1 | *Aspergillus aculeatus* (strain ATCC 16872) | (6) |
| PsoF | Q4WAZ0 | *Aspergillus fumigatus* (strain ATCC MYA-4609) | (7) |
| VdtE | A0A443HK11 | *Byssochlamys spectabilis* | (8) |

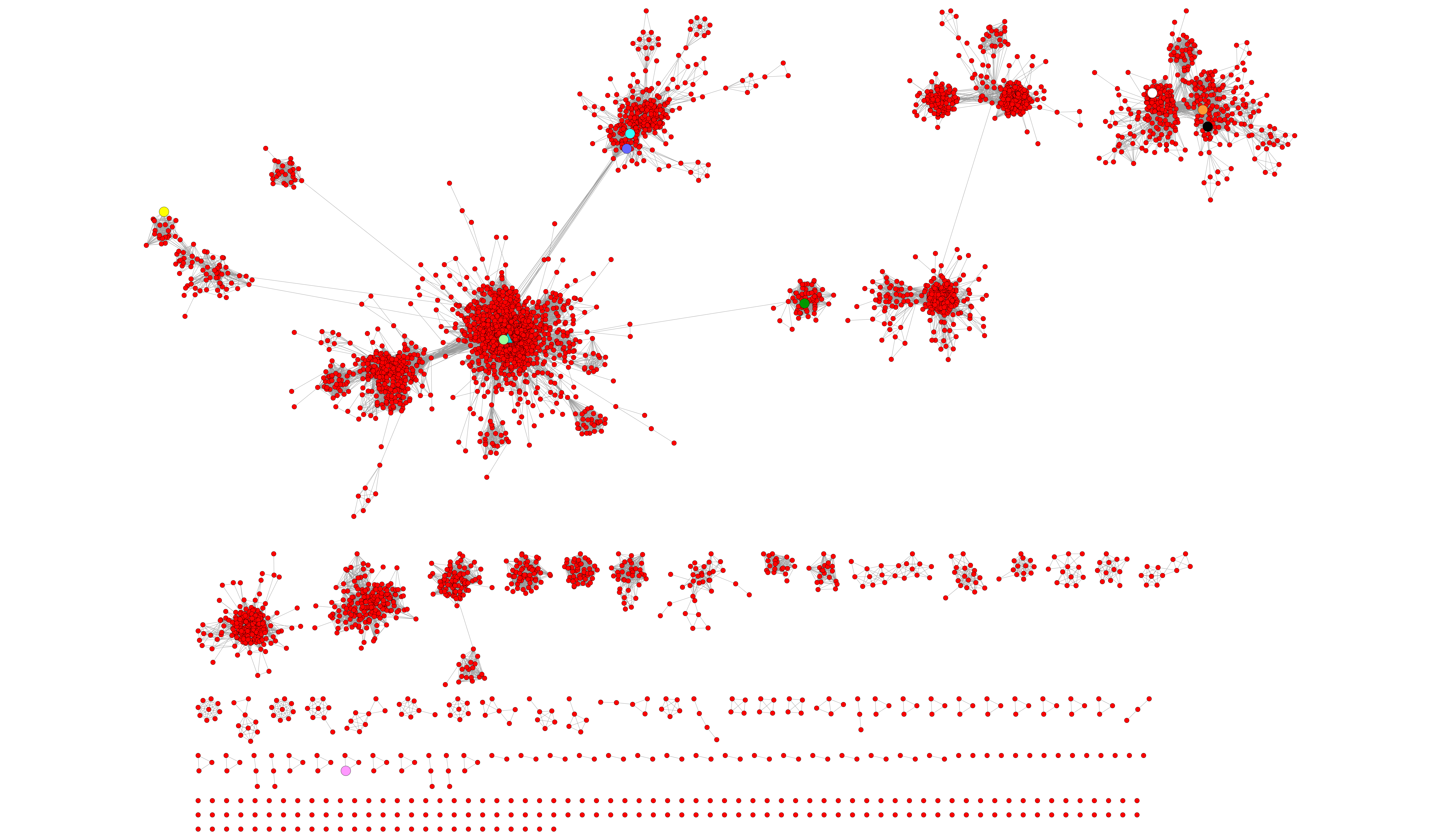

**Figure S3.** Analysis of the daughter network shown in Figure S2 at an AS of 170. With this increased level of stringency, sequences encoding *ssn*BVMO (	), PAMO (	), and *Ac*CHMO ( ) are all found within the largest cluster. Node colours represent the same enzymatic activities as denoted in the legend of Figure S2.

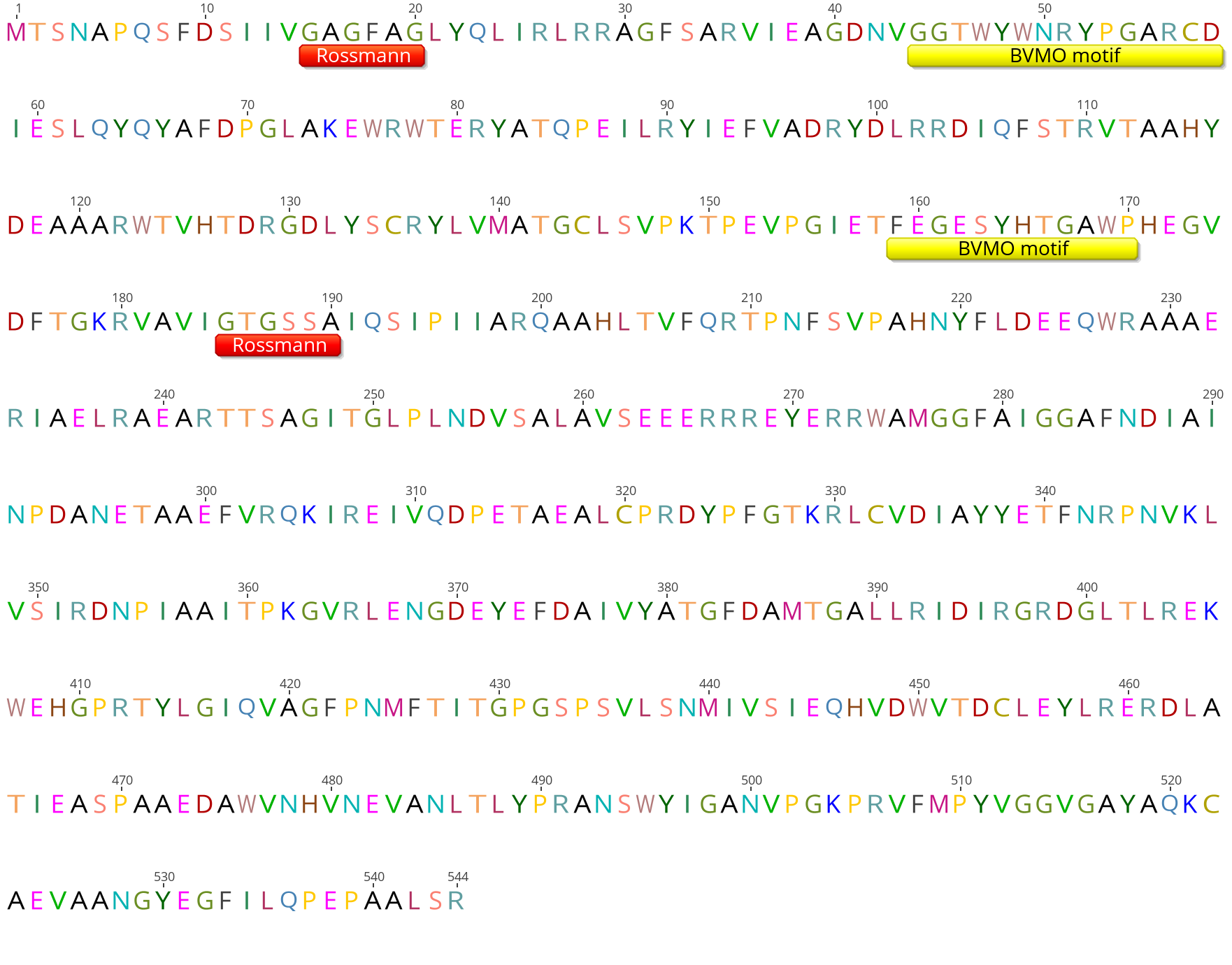

**Figure S4.** The amino acid sequence of *ssn*BVMO with both Rossmann folds (red) and BVMO fingerprint motifs (yellow) annotated.

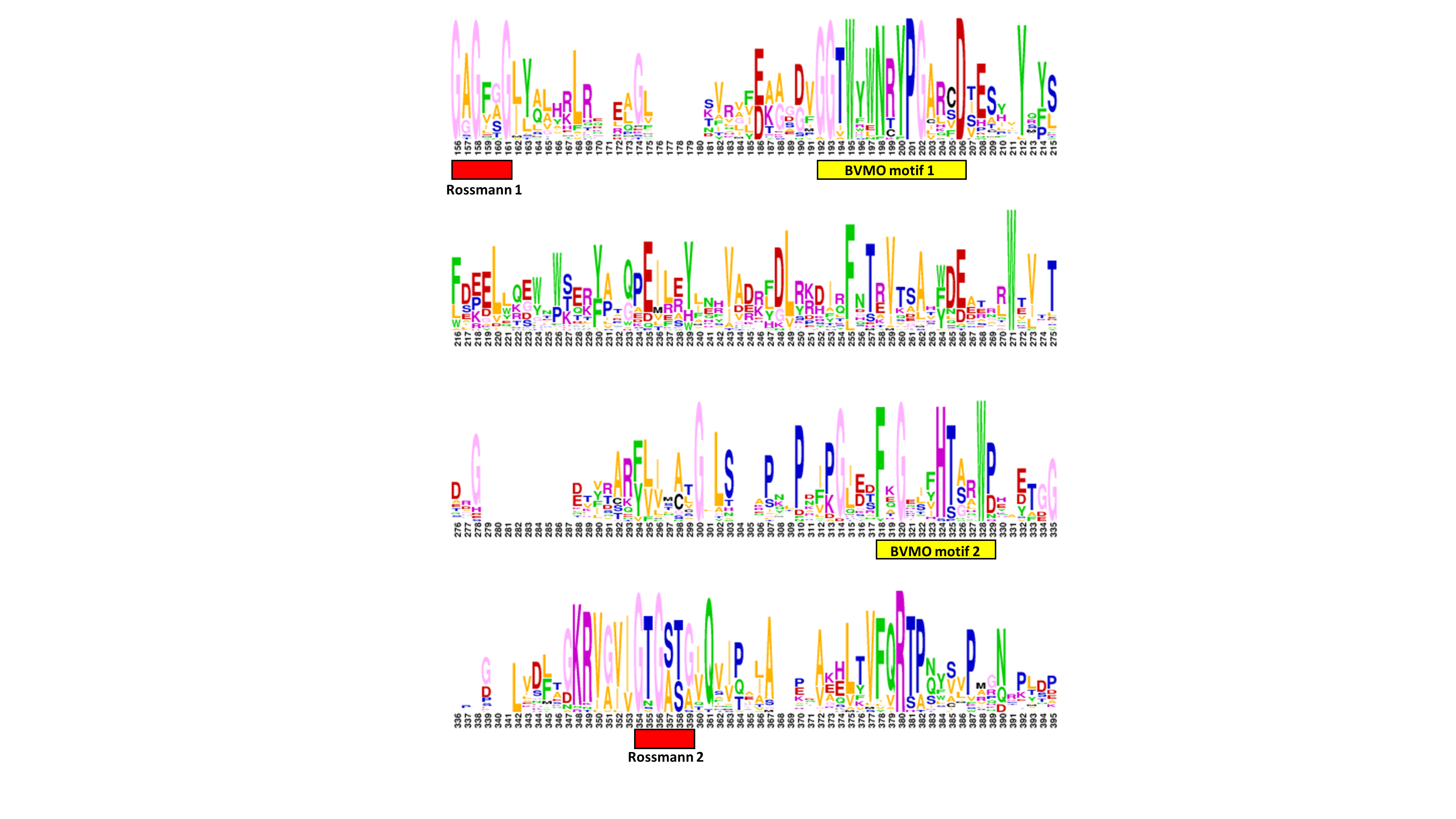

**Figure S5.** A WebLogo illustration of the partial sequence conservation amongst type I BVMOs. Consensus motifs are indicated with coloured bars underneath the relevant sequence. Proteins used in the multiple sequence alignment and the construction of this graphic are given in Table S2.

**Table S2.** Type I BVMOs used in the multiple sequence alignment and construction of the WebLogo shown in Figure S5.

| **BVMO** | **Name** | **Source organism** | **UniProt ID** |
| --- | --- | --- | --- |
| *Rj*BVMO4 | Baeyer-Villiger MO | *Rhodoccous jostii* | Q0SC70 |
| *Co*CPMO | Cyclopentanone 1,2-MO | *Comamonas* sp. | Q8GAW0 |
| *Pl*BVMO | Baeyer-Villiger MO | *Parvibaculum lavamentivorans* | A7HU16 |
| BVMOAfl838 | Baeyer-Villiger MO | *Aspergillus flavus* | B8N653 |
| *Cm*BVMO | Baeyer-Villiger MO | *Cyanidioschyzon merolae* | M1VDM5 |
| *Go*ACMO | Acetone MO | *Gordonia* sp. TY-5 | A1IHE6 |
| *Pv*MEKMO | Methyl ethyl ketone MO | *Pseudomonas veronii* | Q0MRG6 |
| BVMOAfl706 | Baeyer-Villiger MO | *Aspergillus flavus* | B8NCF3 |
| *Rp*CHMO | Cyclohexanone MO | *Rhodococcus* sp. Phi1 | Q84H73 |
| *Rh*CHMO | Cyclohexanone MO | *Rhodococcus* sp. HI-31 | C0STX7 |
| *Ob*BVMO | Baeyer-Villiger MO | *Pseudooceanicola batsensis* | A3U3H1 |
| *Ac*CHMO | Cyclohexanone 1,2-MO | *Acinetobacter calcoaceticus* | P12015 |
| *Tm*CHMO | Cyclohexanone MO | *Thermocrispum municipale* | A0A1L1QK39 |
| *Cr*CAMO | Cycloalkanone MO | *Cylindrocarpon radicicola* | G8H1L8 |
| *Rj*BVMO24 | Baeyer-Villiger MO | *Rhodoccous jostii* | Q0S5T2 |
| *Rr*STMO | Steroid MO | *Rhococcus rhodochrous* | O50641 |
| *Ct*SAPMO | 4-Sulfoacetophenone MO | *Comamonas testosteroni* | B7X4D9 |
| *Tf*PAMO | Phenylacetone MO | *Thermobifida fusca* | Q47PU3 |
| PockeMO | Polycyclic ketone MO | *Thermothelomyces thermophila* | G2QA95 |
| *Ps*CPDMO | Cyclopentadecanone MO | *Pseudomonas* sp. HI-70 | T2HVF7 |
| *Di*BVMO4 | Baeyer-Villiger MO | *Dietzia* sp. D5 | U5S003 |
| *Sa*PtlE | Neopentalenolactone D synthase | *Streptomyces avermitilis* | Q82IY8 |
| *Sl*BVMO | Baeyer-Villiger MO | *Streptomyces leeuwenhoekii* | A0A0F7W6X7 |
| *Pp*OTEMO | 2-oxo-∆^3^-4,5,5-trimethylcyclopentenylacetyl-CoA MO | *Pseudomonas putida* | H3JQW0 |
| *Lb*BVMO | Baeyer-Villiger MO | *Leptospira biflexa* | B0SRK0 |
| *Pf*HAPMO | 4-Hydroxyacetophenone MO | *Pseudomonas fluorescens* | Q93TJ5 |
| *Mt*EthA | Flavin-containing MO | *Mycobacterium tuberculosis* | P9WNF9 |
| BVMO_Halo_ | Cyclohexanone monooxygenase | *Halopolyspora algeriensis* | A0A368VHL5 |
| *ssn*BVMO | Cyclohexanone monooxygenase | *Chloroflexota* bacterium strain G233 | A0A2A9HFE7 |

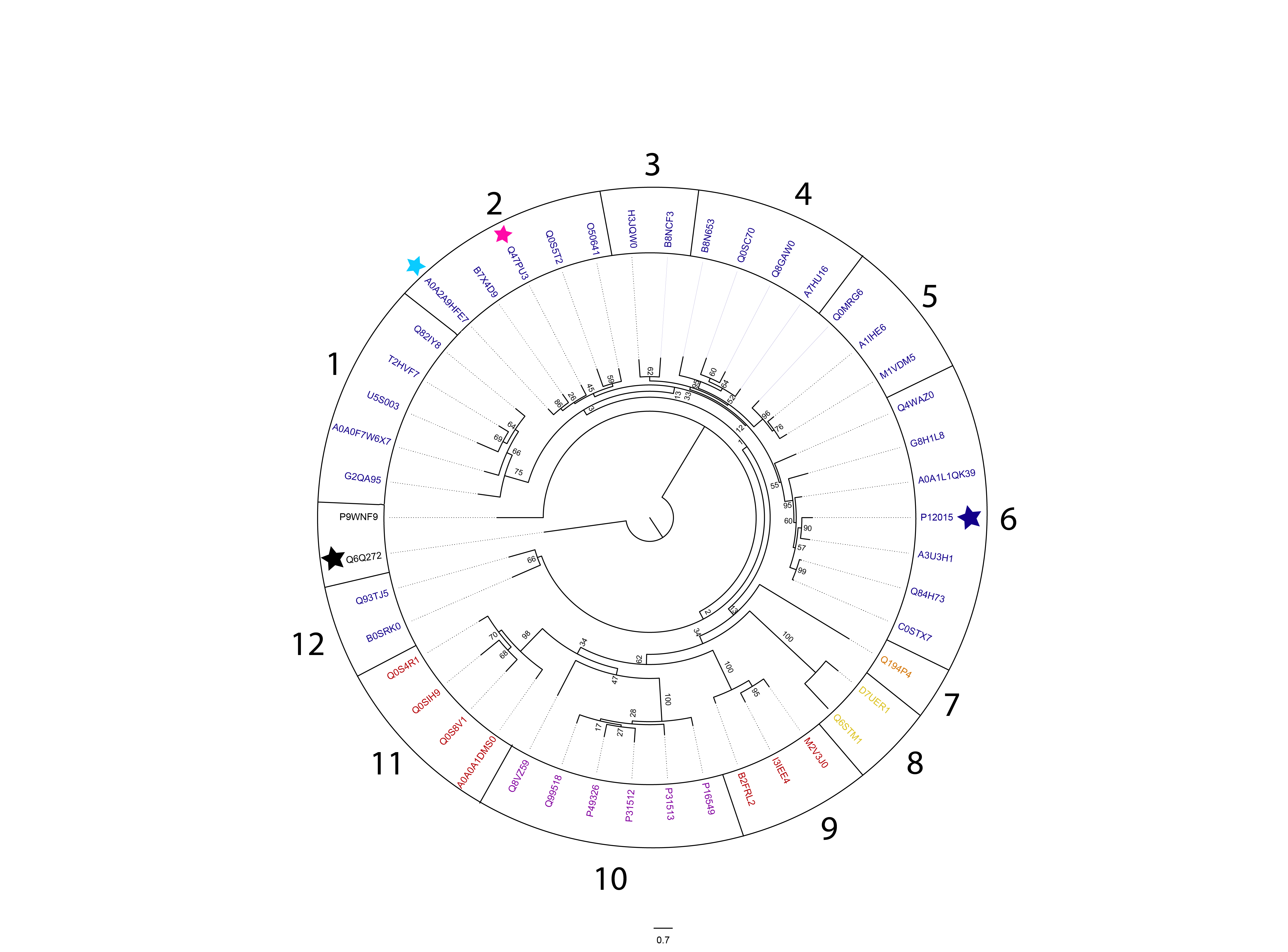

**Figure S6.** Phylogenetic tree using 46 BVMO sequences with bootstrap values supporting the 12 subgroups are indicated at each node/branch. The color of the sequences in the subgroups represents the flavoprotein group to which the respective BVMOs belong (blue for type I BVMOs, yellow for type II BVMOs, orange for type O BVMOs, purple for type I FMOs, and red for type II FMOs). *Ac*CHMO is denotated with a blue star, PAMO is denoted with a magenta start, and *ssn*BVMO is denoted with a cyan star. A Class D flavoprotein (Q6Q272) was included as an outgroup to root the tree (black star). Each sequence is denoted with its Uniprot Accession number.

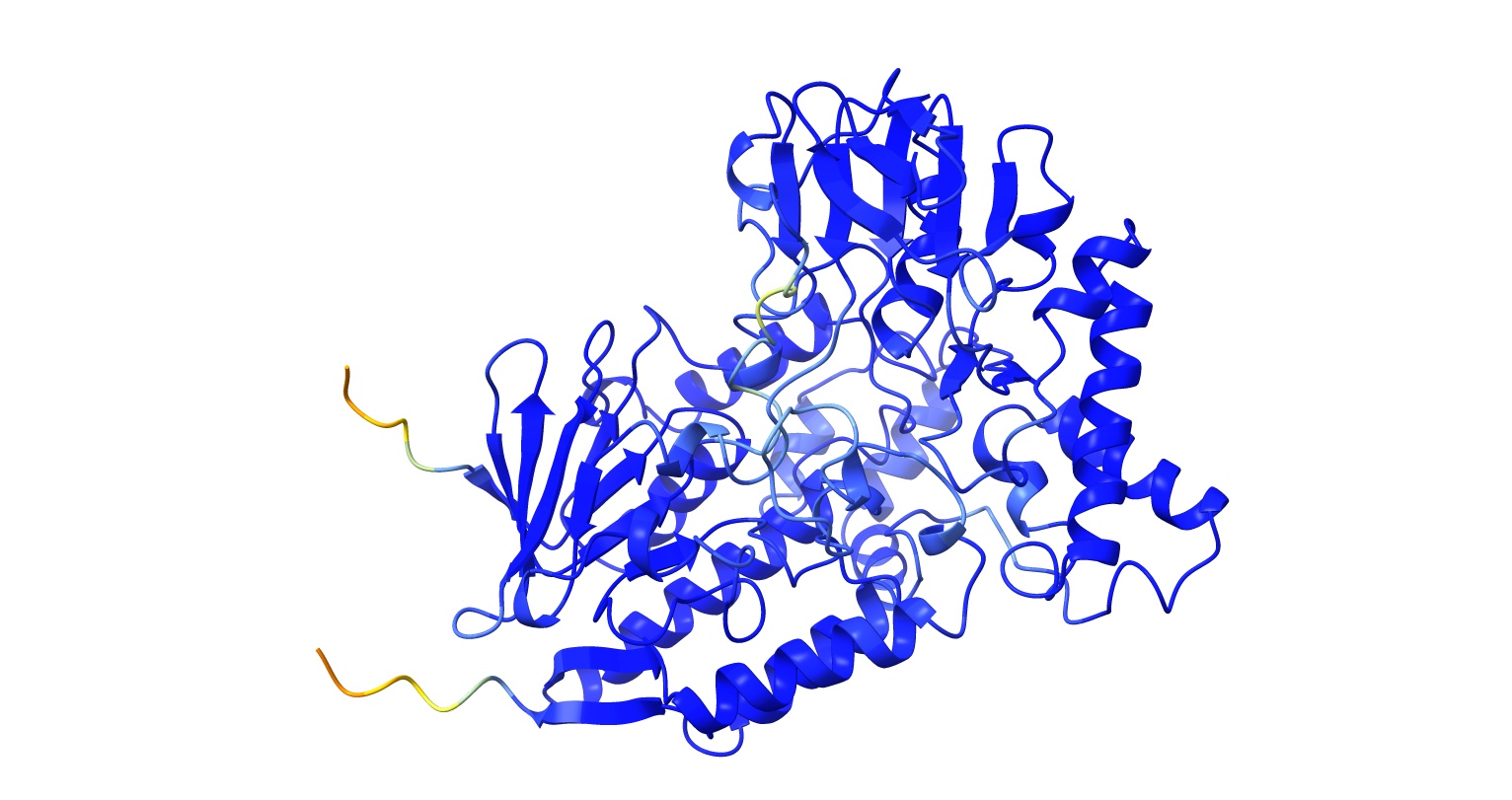

**Figure S7.** AlphaFold prediction of *ssn*BVMO with pLDDT plot colour-overlayed over the three-dimensional predicted structure.

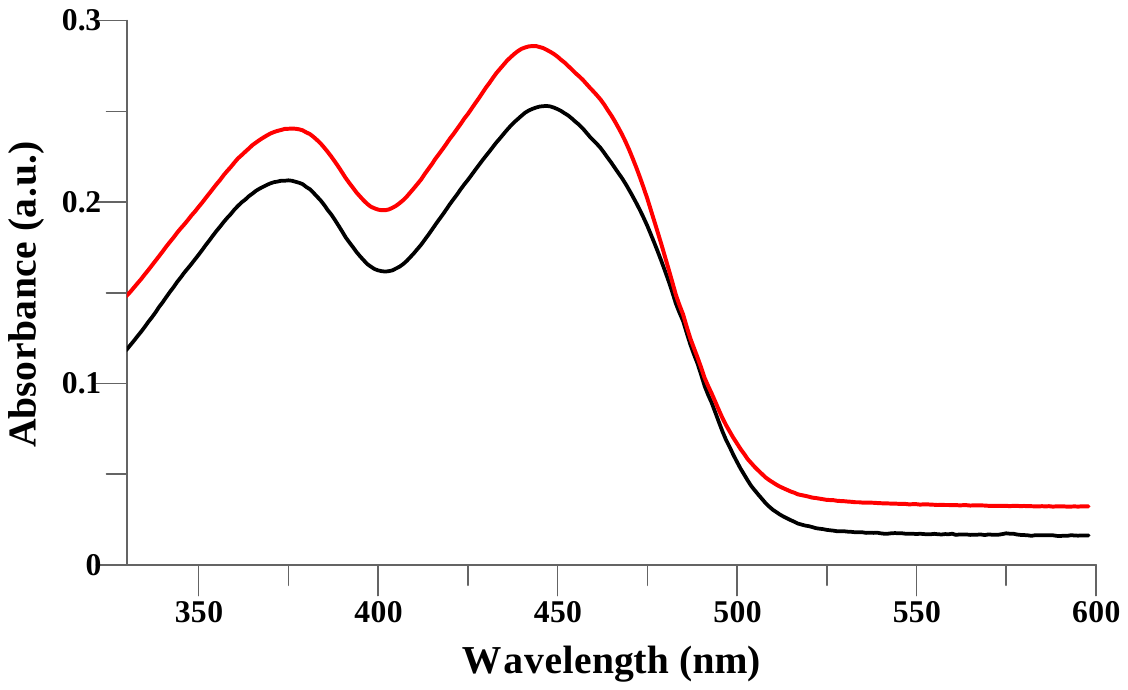

**Figure S8**. Absorbance spectra of FAD supplemented *ssn*BVMO stock at t = 0.5 min (**−**) and absorbance spectra of FAD supplemented *ssn*BVMO stock at t = 10 min (**−**). The blueshift shift in λ_max_ 450 nm is consistent with uptake of added FAD by *ssn*BVMO.

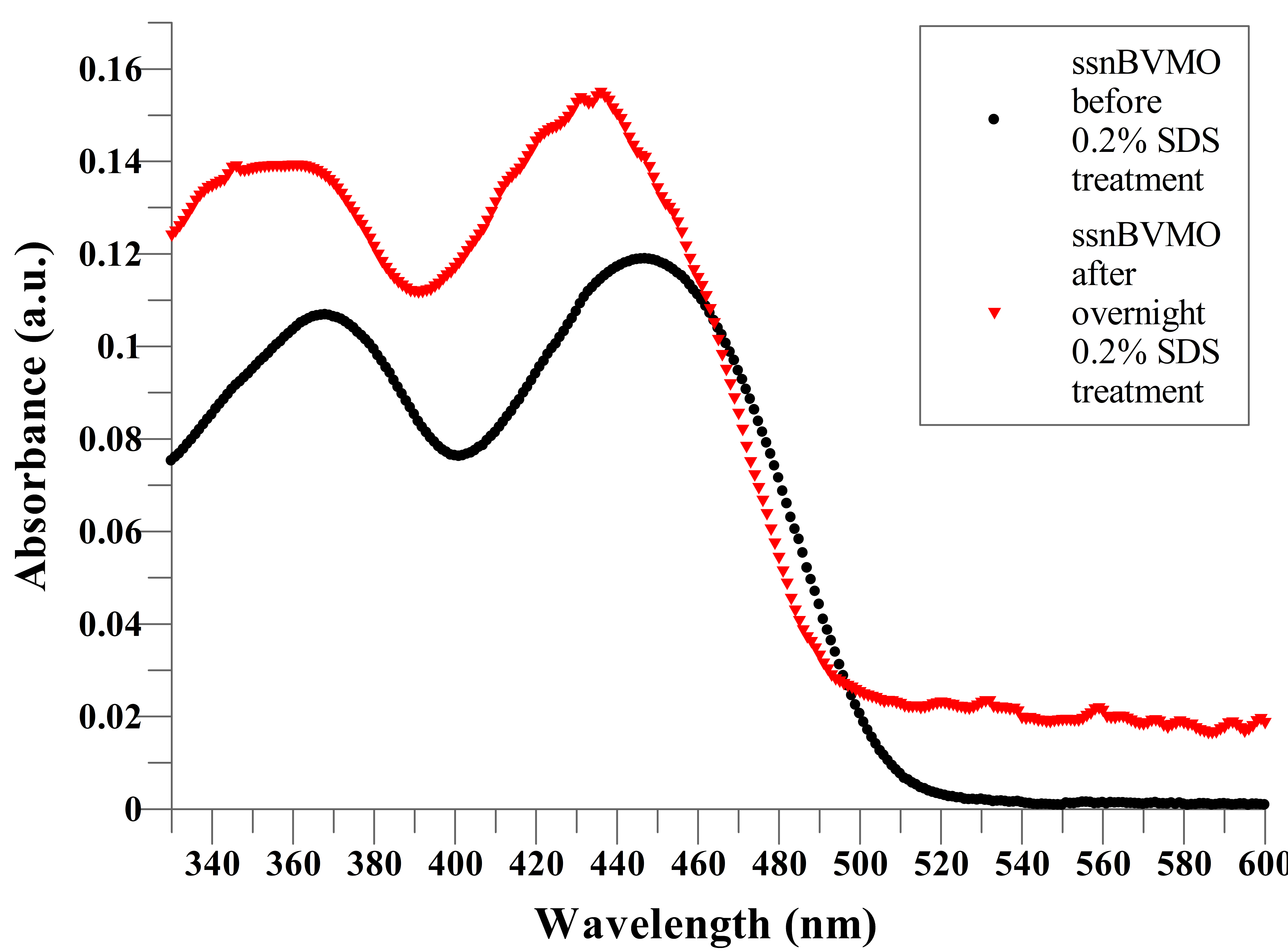

**Figure S9.** Absorbance change for *ssn*BVMO treated with 0.2% SDS overnight. Note the shift in λ_max_ from 430 nm to characteristic 450 nm for unbound FAD, suggesting that all protein bound flavin has been released after SDS treatment allowing for determination of molar extinction coefficient for the enzyme and justifying the flavin supplementation of *ssn*BVMO purifications.

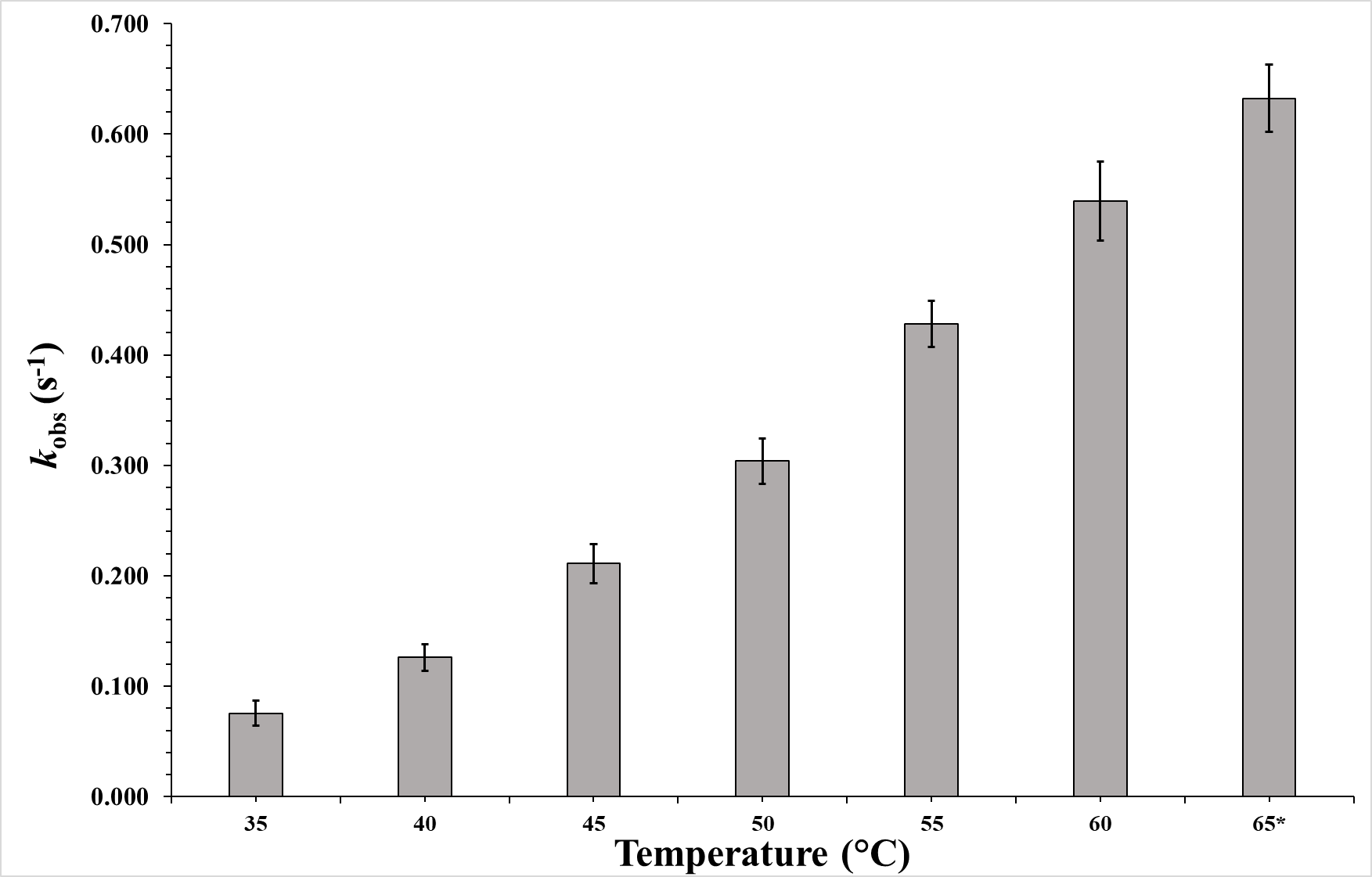

**Figure S10.** Changes in *k*_obs_ with changing temperature. *- Indicates that the absorbance started increasing after some time. The errors are represented by standard deviation derived from running measurements in triplicates. Kinetic measurements were carried out using the following concentrations, [*ssn*BVMO]= 1 µM, [NADPH]= 0.1 mM, and [2-octanone]= 5 mM.

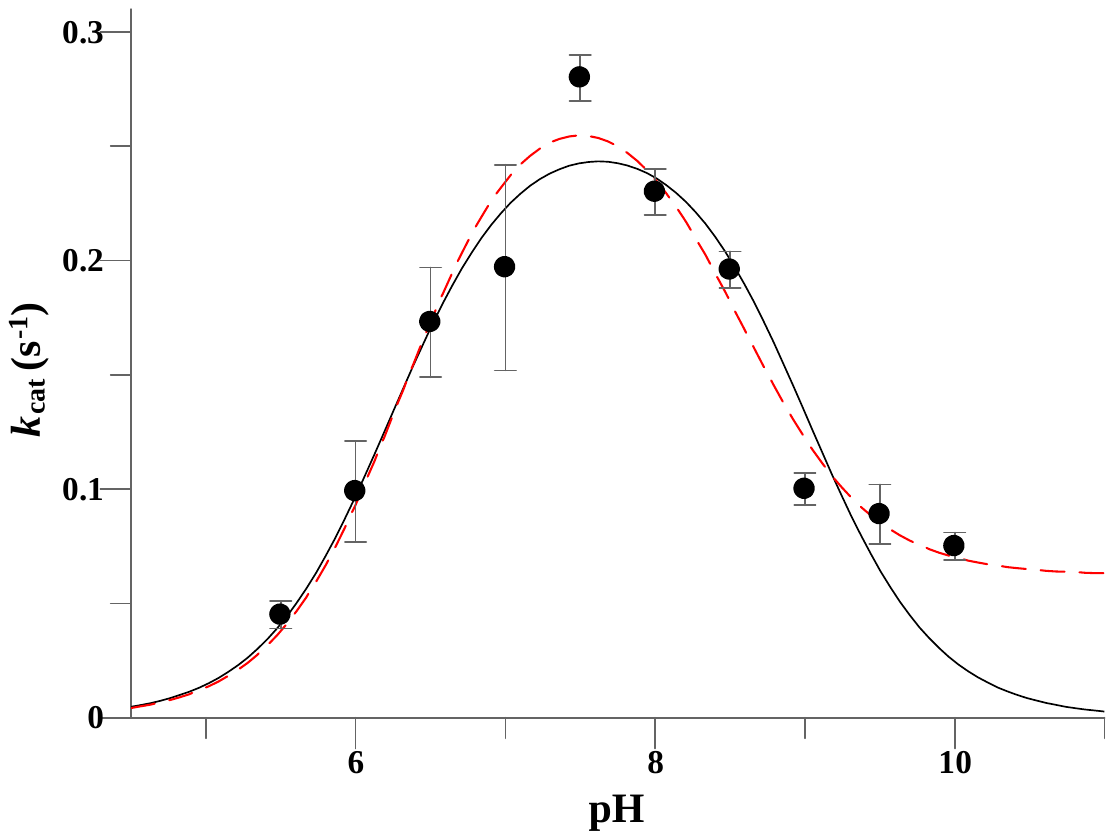

**Figure S11.** The pH-rate profile for the *ssn*BVMO -catalyzed oxidation of 2-octanone suggests that two ionizable residues influence catalysis by *ssn*BVMO. Data is fit to Eqn. 1 with *k*ʹʹ_cat_ either held fixed at zero (**−**) or allowed to vary during the non-linear regression analysis (− −).

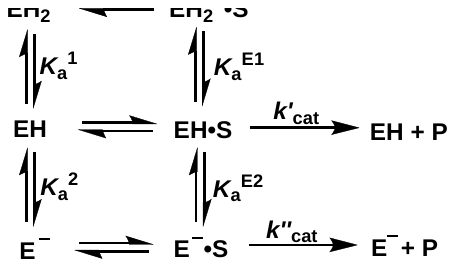

**Scheme S1.** The simplified kinetic scheme used to interpret the dependence of the apparent *k*_cat_ values on pH.

$k_{cat}=\frac{k_{cat}^{'}K_{a}^{1}\left[ H^{+} \right]+k_{cat}^{''}K_{a}^{1}K_{a}^{2}}{({\left[ H^{+} \right]^{2} + K}_{a}^{1}\left[ H^{+} \right] + K_{a}^{1}K_{a}^{2})}$	(**Eqn. S1**)

**Table S3.** Kinetic parameters derived from fitting the pH-rate profile data to Eqn. 1 with *k*ʹʹ_cat_ = 0 (Fit 1) and with *k*ʹʹ_cat_ ≠ 0 (Fit 2).

| **Parameter** | **Fit 1** | **Fit 2** |
| --- | --- | --- |
| *k*ʹ_cat_ (s^-1^) | 0.26 ± 0.02 | 0.29 ± 0.02 |
| *k*ʹʹ_cat_ (s^-1^) | Fixed at 0 | 0.06 ± 0.02 |
| p*K*_a_^1^ | 6.2 ± 0.2 | 6.3 ± 0.1 |
| p*K*_a_^2^ | 9.0 ± 0.2 | 8.6 ± 0.2 |

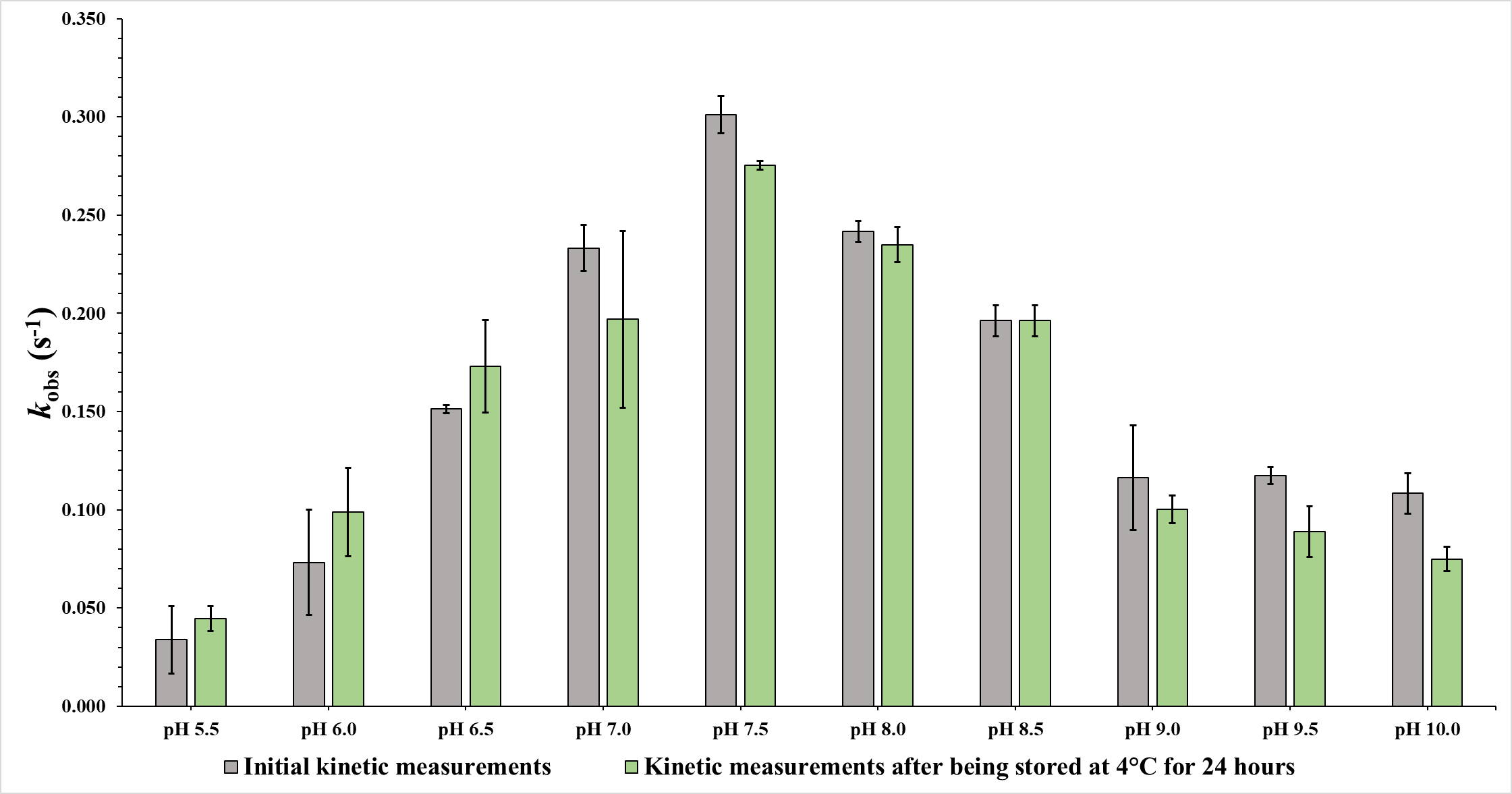

**Figure S12.** Changes in *k*_obs_ with changing pH. The errors are represented by standard deviation derived from running measurements in triplicates. Kinetic measurements were carried out using the following concentrations, [*ssn*BVMO] = 1 µM, [NADPH] = 0.1 mM, and [2-octanone] = 5 mM.

**Table S4.** Amino acid linker sequences used to fuse *ssn*BVMO and 17X-PTDH to prepare the self-sufficient fusion enzyme.

| **Linker Variant Code** | **Linker Sequence** |
| --- | --- |
| L1 | SRSAAG |
| L2 | SSATGSATGSAG |
| L3 | W |
| L4 | ALIPG |
| L5 | PMHFSTHNYY |

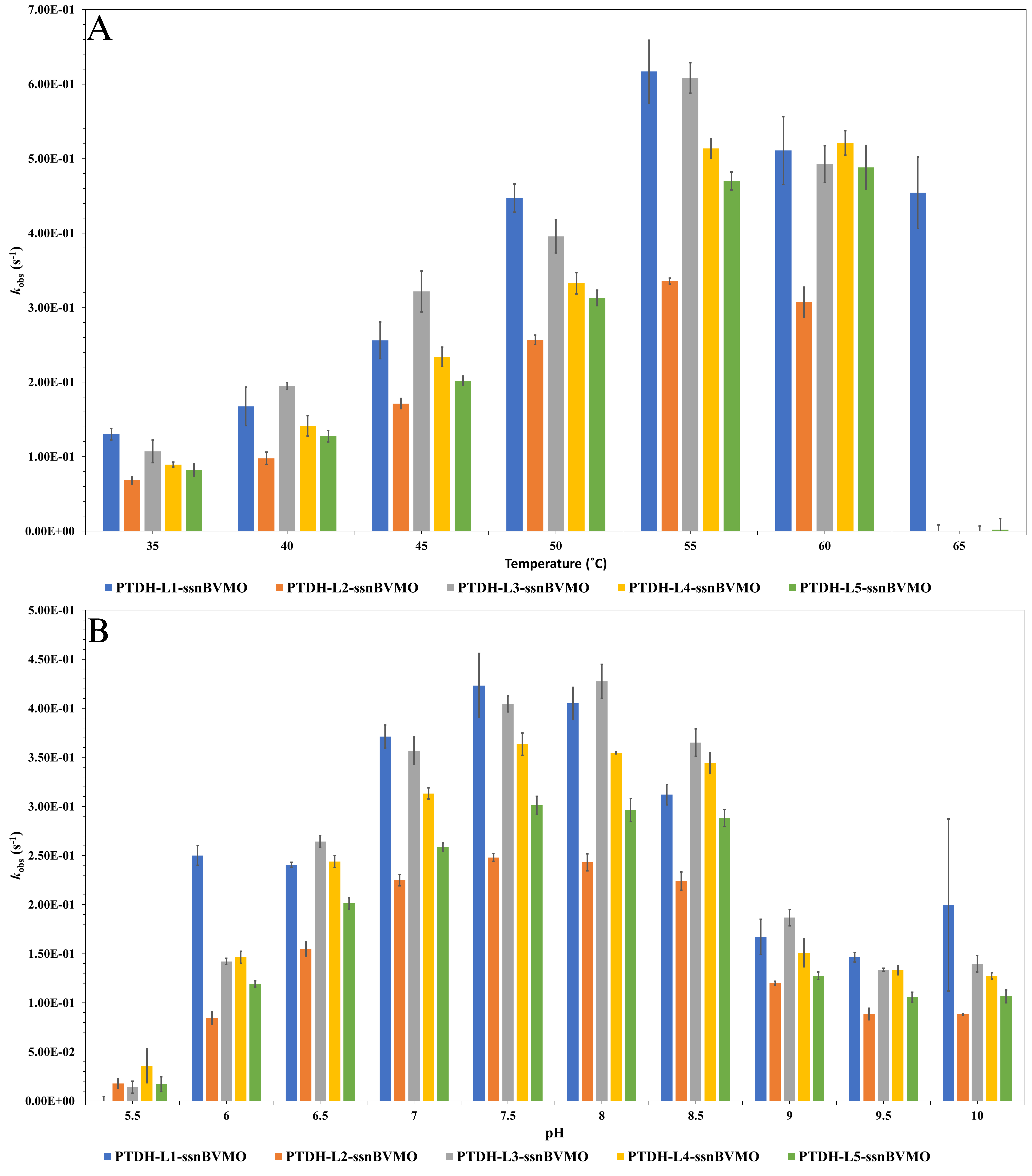

**Figure S13.** Observed rate constants for initial rates of oxidation by PTDH-L-*ssn*BVMO variants at varying temperature (A) and pH conditions (B), [PTDH-L-*ssn*BVMO] = 1 µM, [NADPH] = 0.1 mM, and [2-octanone] = 5 mM.

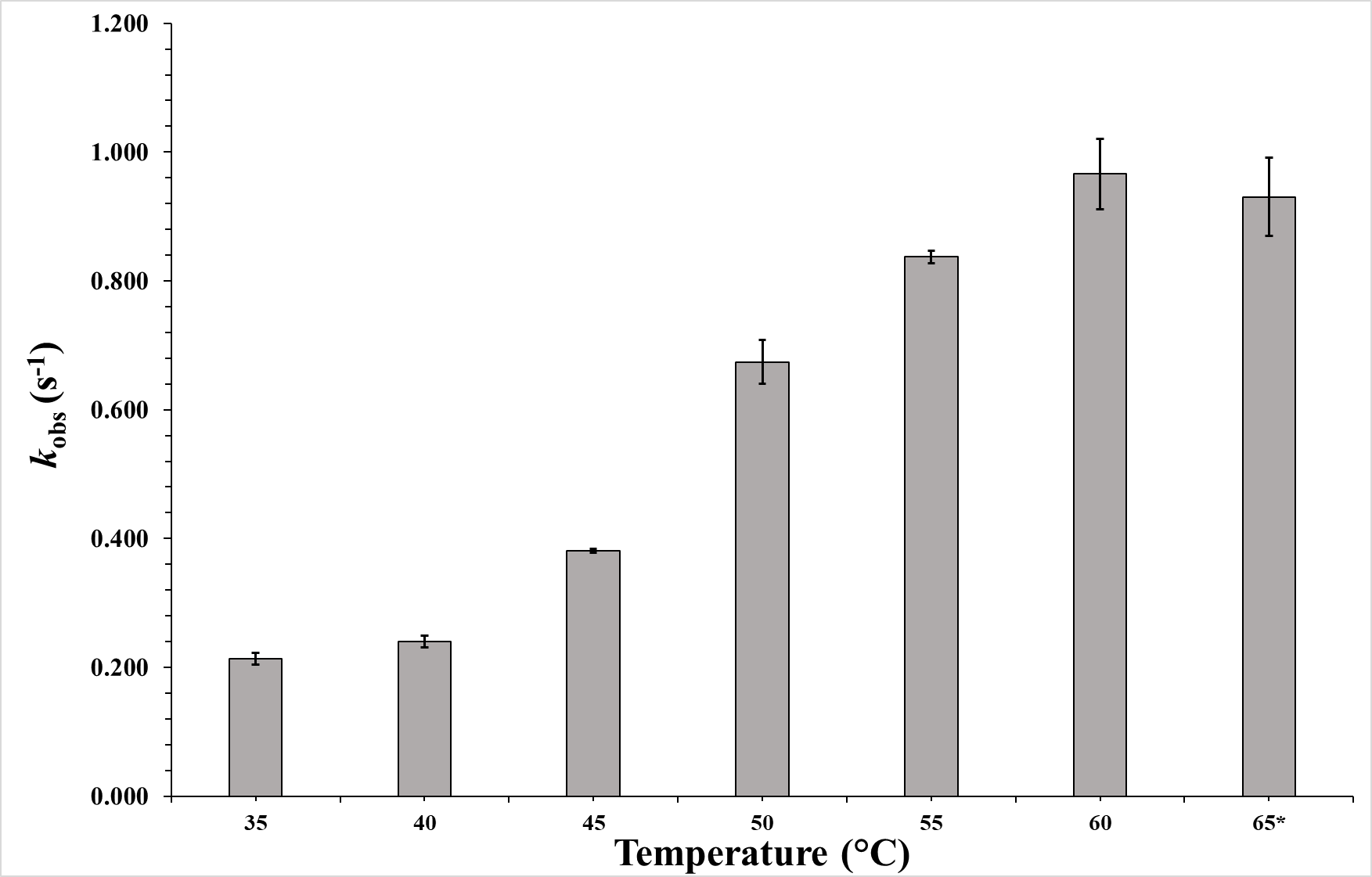

**Figure S14.** Observed rate constants for initial rates of oxidation by FAD-supplemented PTDH-L1-*ssn*BVMO at varying temperatures. Error bars represent standard deviation obtained from running measurements in triplicate. Kinetic measurements were carried out using the following concentrations, [PTDH-L1-*ssn*BVMO] = 1 µM, [NADPH] = 0.1 mM, and [2-octanone] = 5 mM.

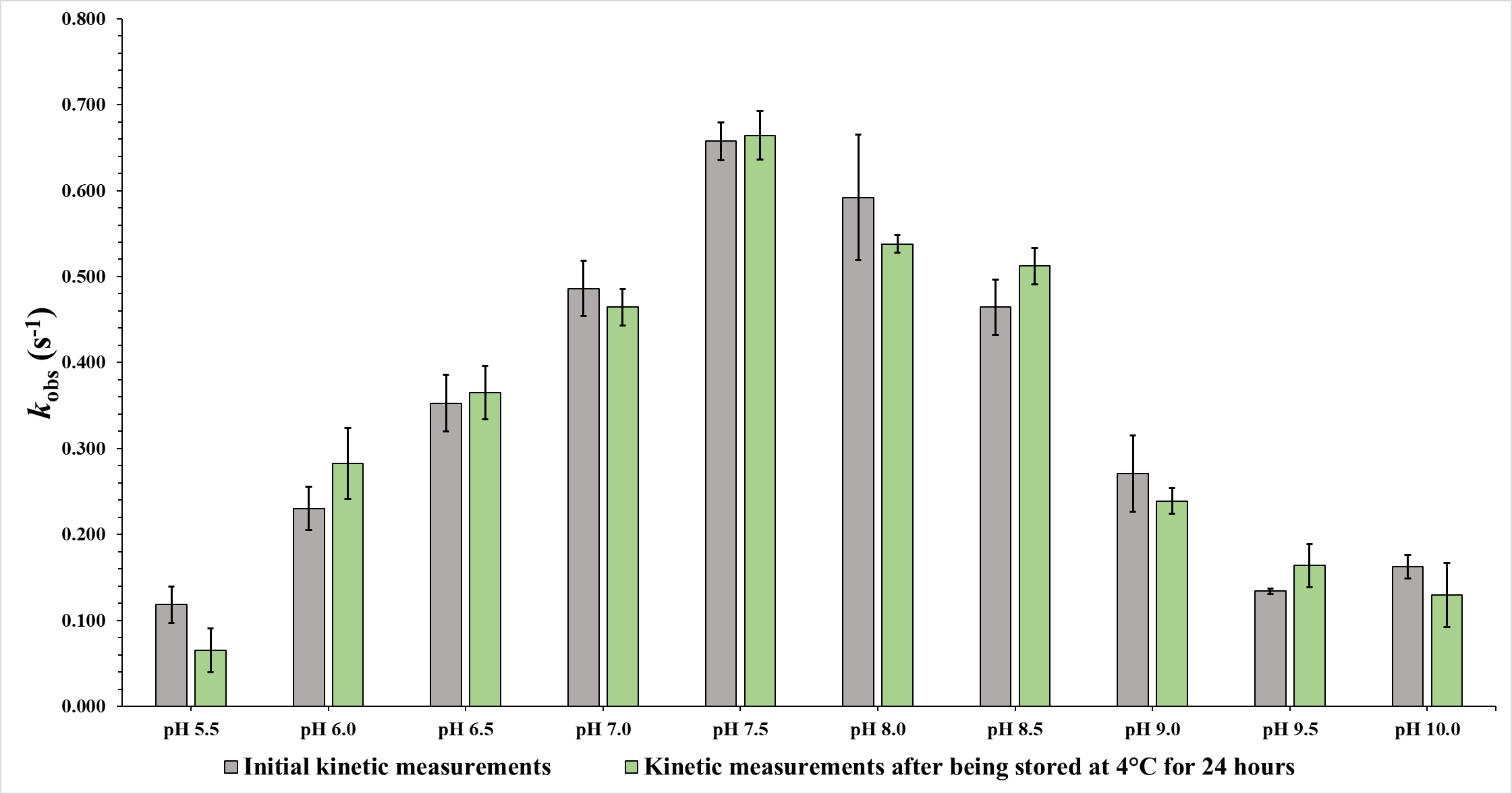

**Figure S15.** Observed rate constants for initial rates of oxidation by FAD-supplemented PTDH-L1-*ssn*BVMO at varying temperatures. Error bars represent standard deviation obtained from running measurements in triplicate. Kinetic measurements were carried out using the following concentrations, [PTDH-L1-*ssn*BVMO] = 1 µM, [NADPH] = 0.1 mM, and [2-octanone] = 5 mM.

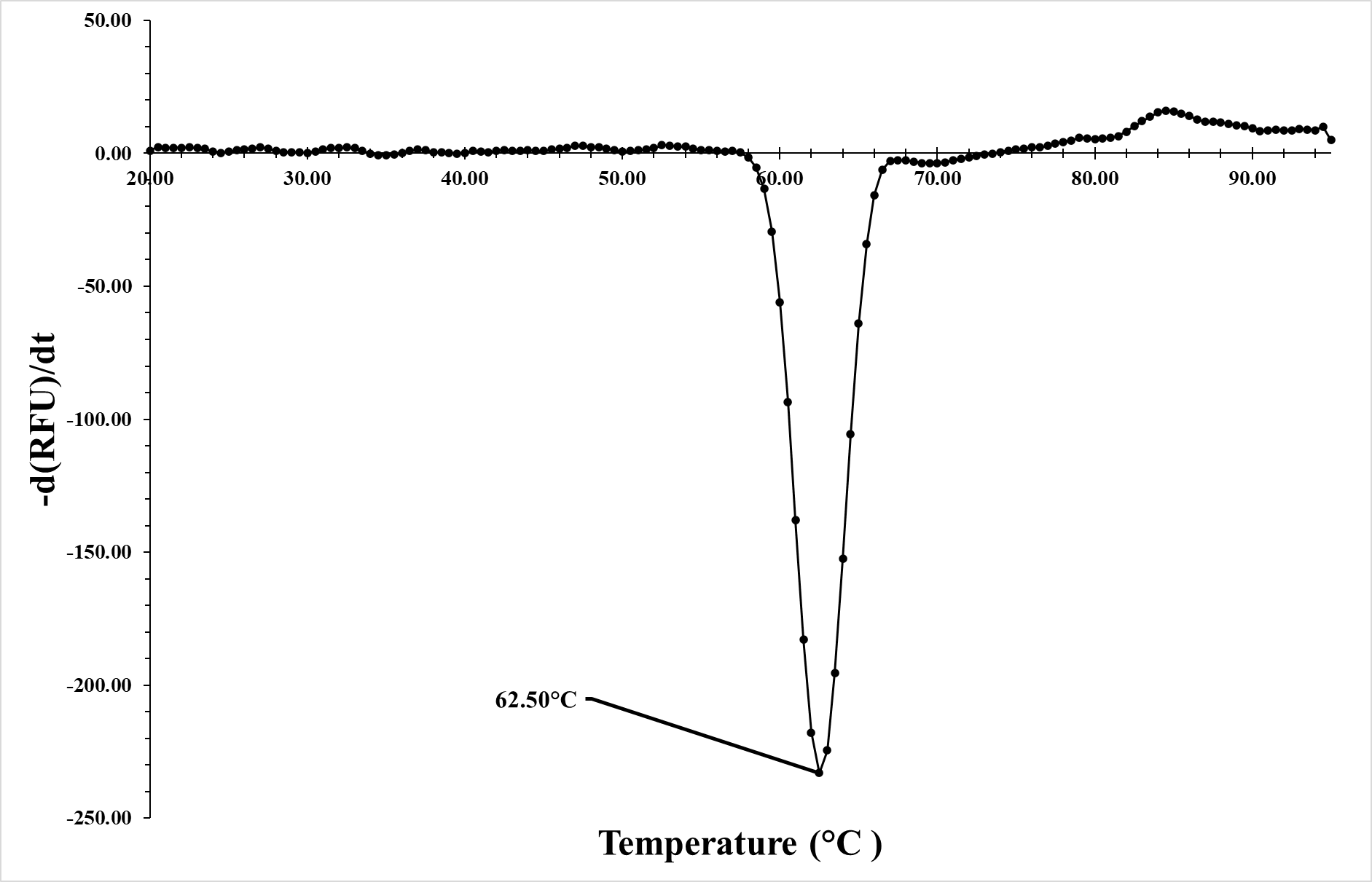

**Figure S16.** 1^st^ derivative plot for fluorescence of *ssn*BVMO vs. changing temperature.

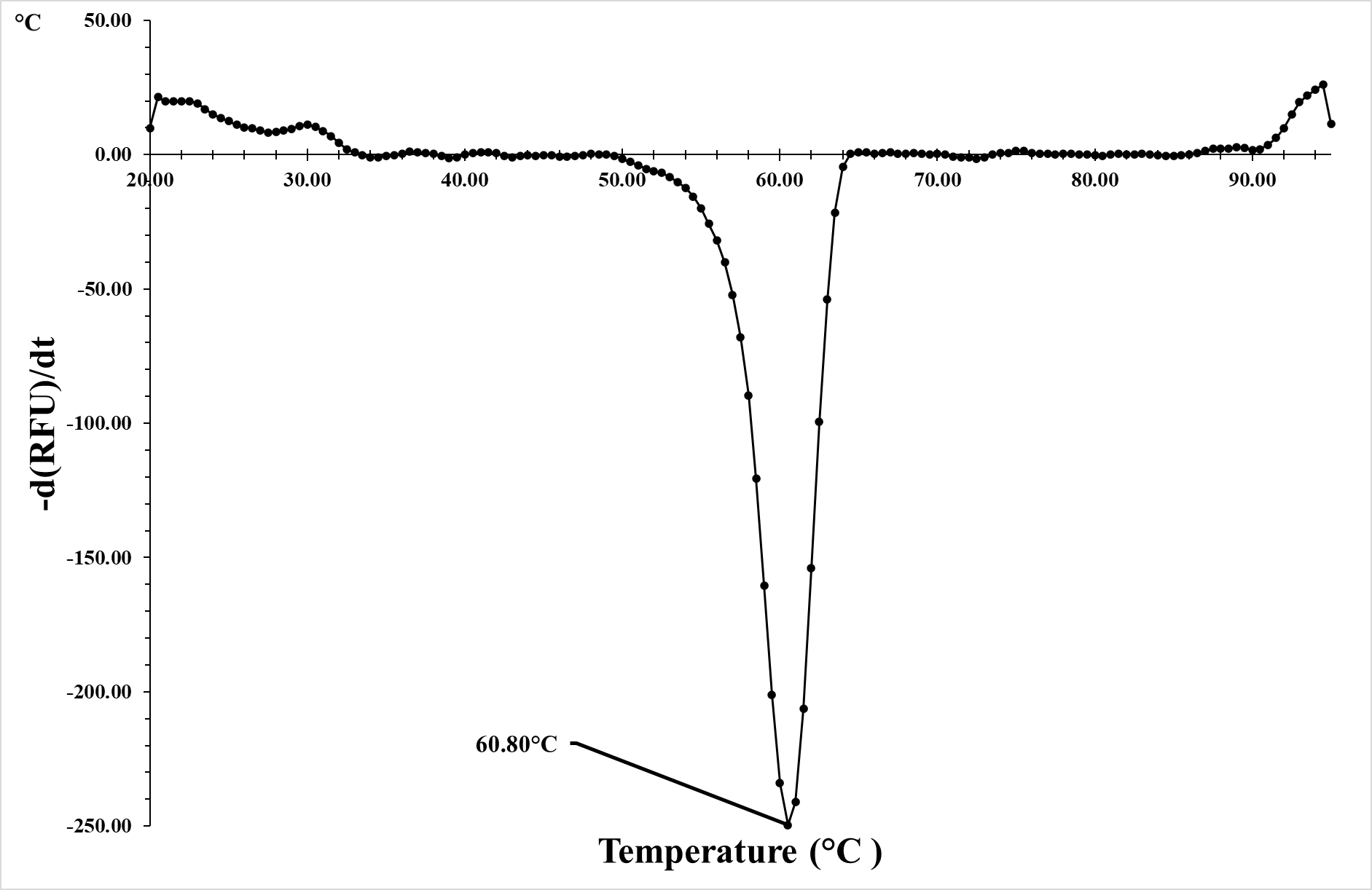

**Figure S17.** 1^st^ derivative plot for fluorescence of PTDH-L1-*ssn*BVMO vs. changing temperature.

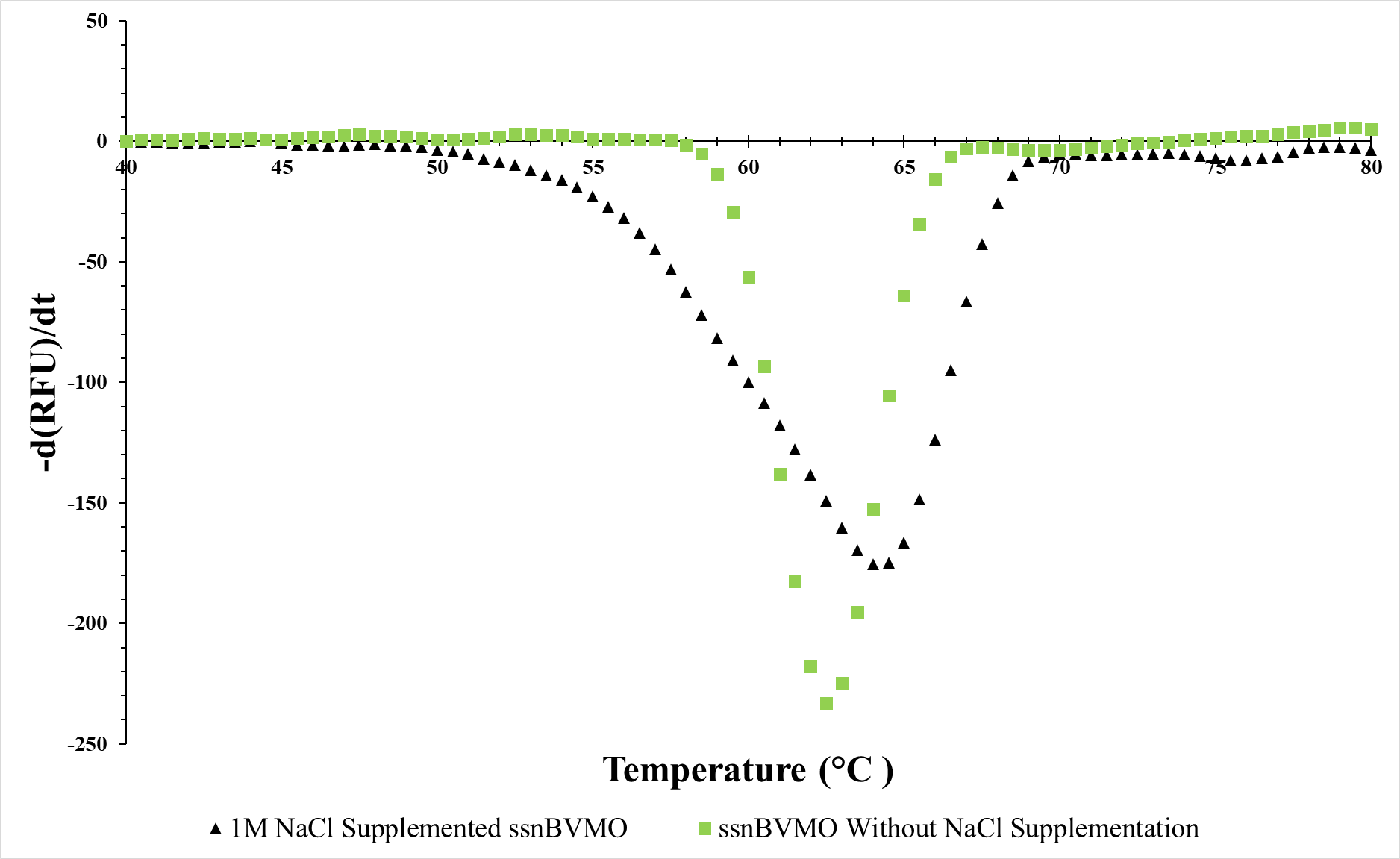

**Figure S18.** Comparison of 1^st^ derivative plots for fluorescence of *ssn*BVMO samples with/without NaCl supplementation vs. changing temperature.

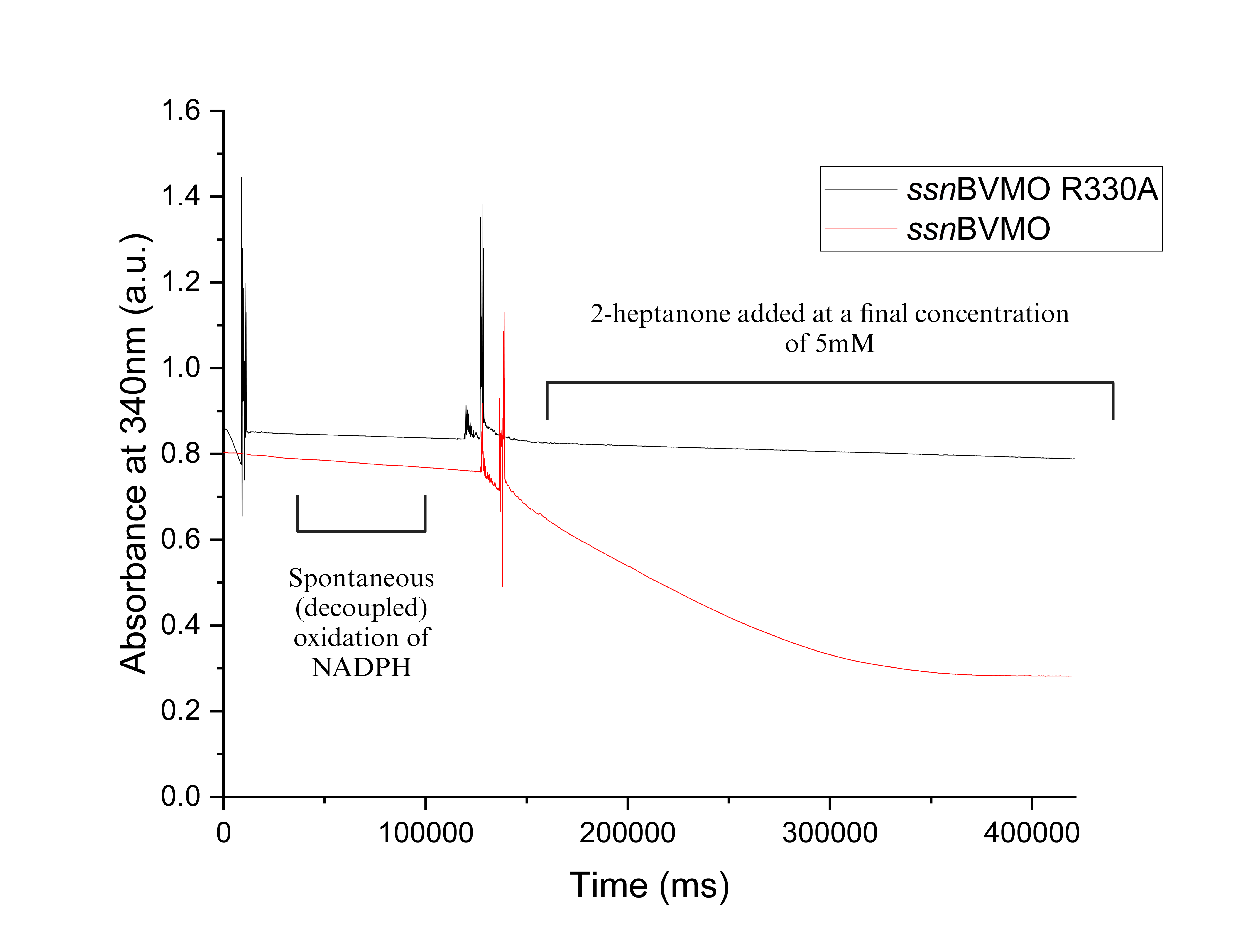

**Figure S19.** Kinetic assay measured at 340 nm for the oxidation of 2-heptanone (5 mM) using 1.0 µM of native *ssn*BVMO or *ssn*BVMO R330A and 0.1 mM NADPH. Initial measurements are completed in absence of 2-heptanone to record the rate of spontaneous NADPH oxidation. BVMO-catalysed oxidation begins at approximately 150 seconds.

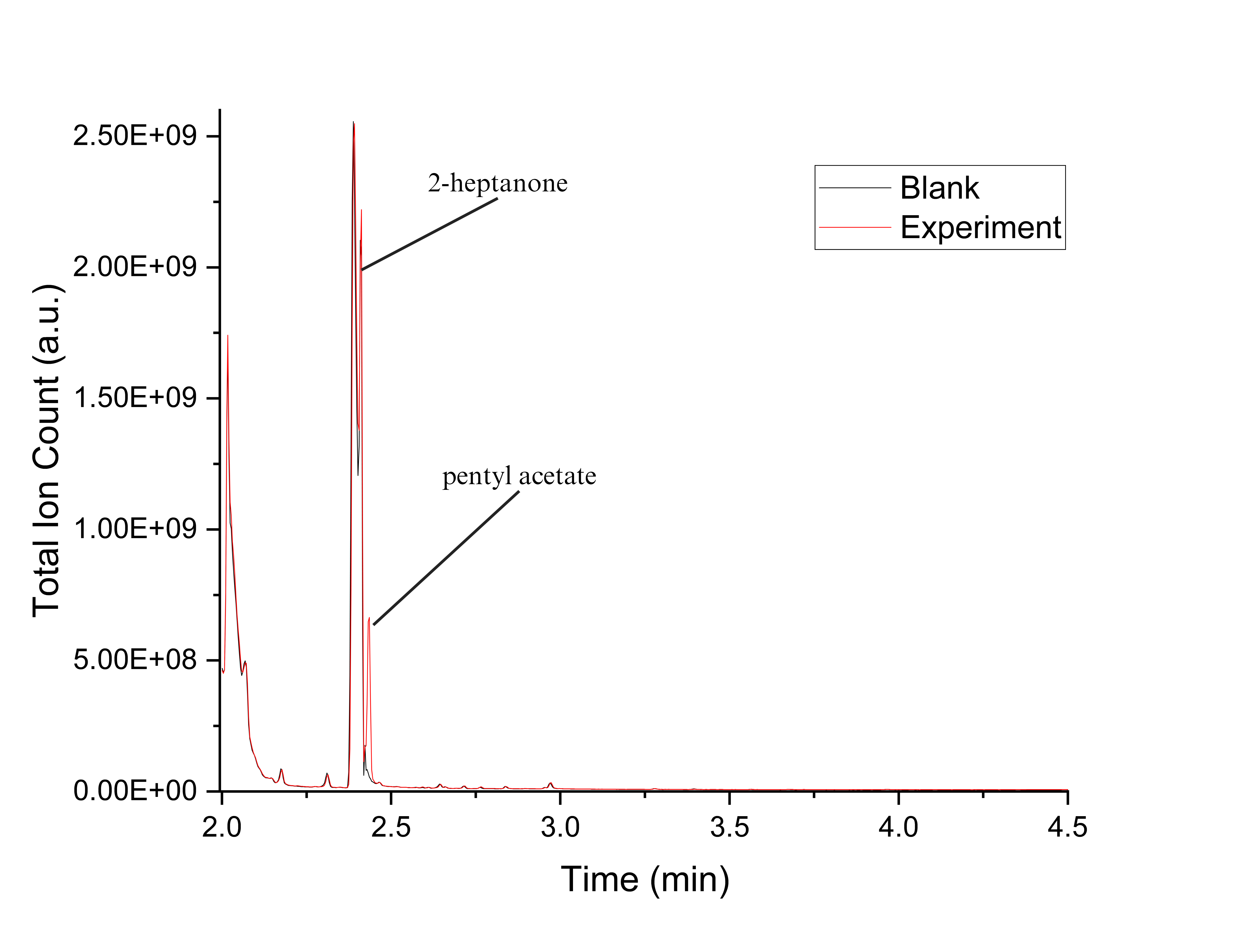

**Figure S20.** GCMS chromatogram for *ssn*BVMO reaction with 2-heptanone.

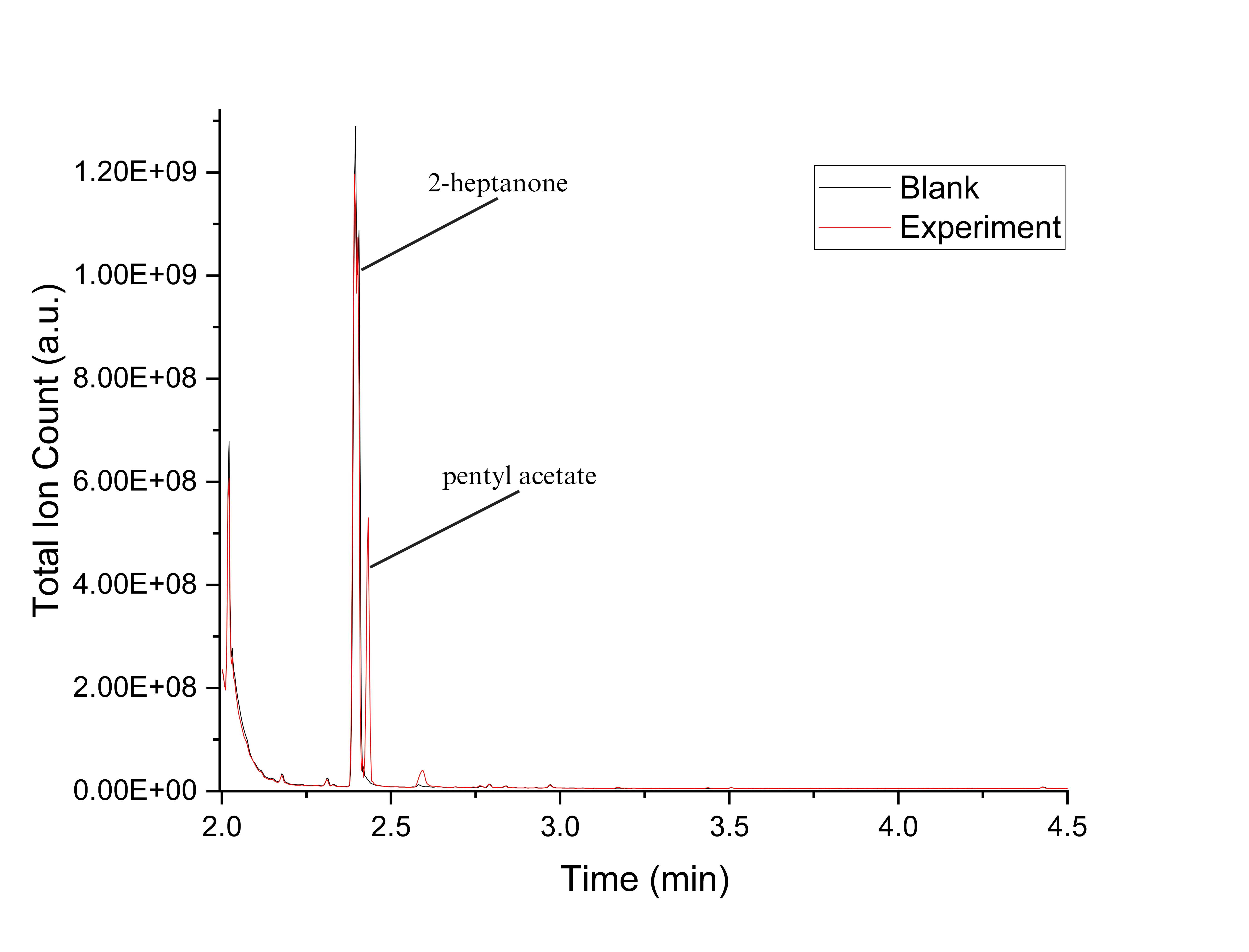

**Figure S21.** GCMS chromatogram for PTDH-L1-*ssn*BVMO reaction with 2-heptanone

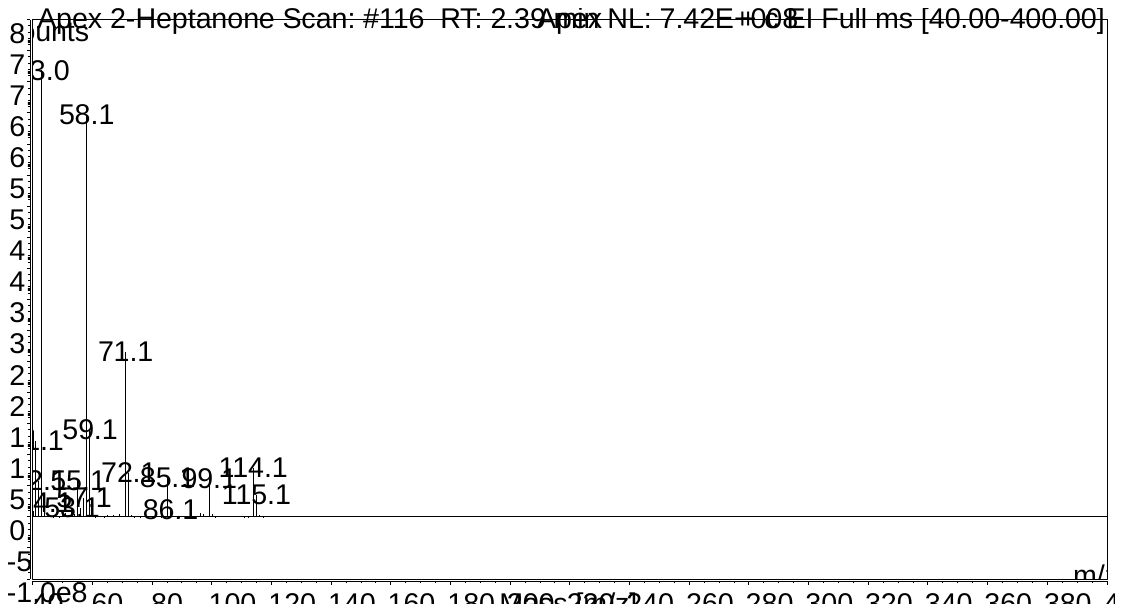

**Figure S22.** Fragmentation pattern for 2-heptanone.

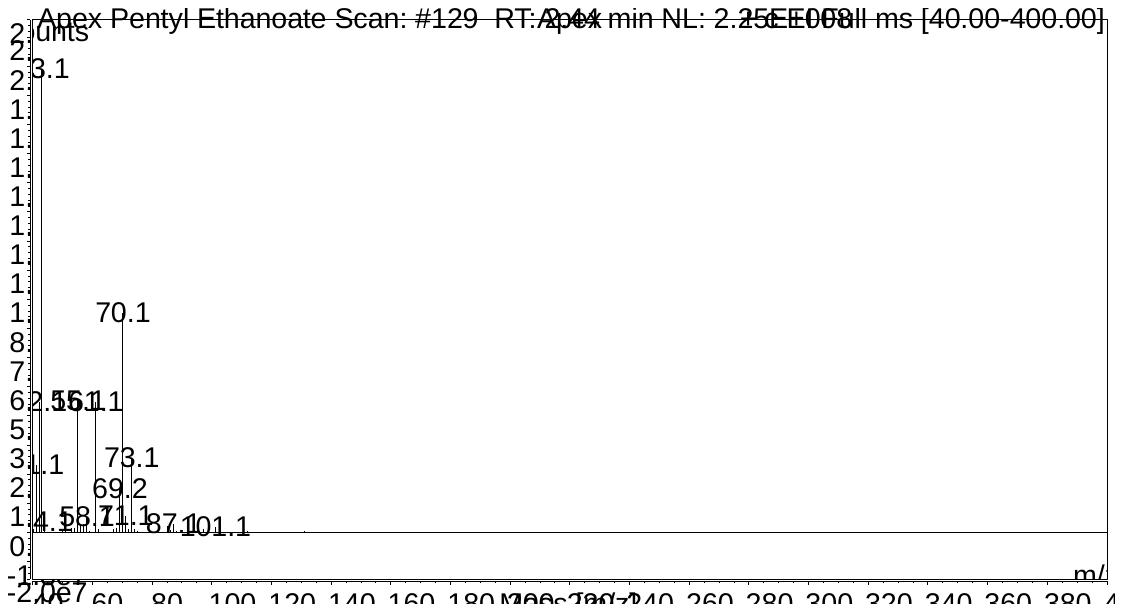

**Figure S23.** Fragmentation pattern for pentyl acetate.

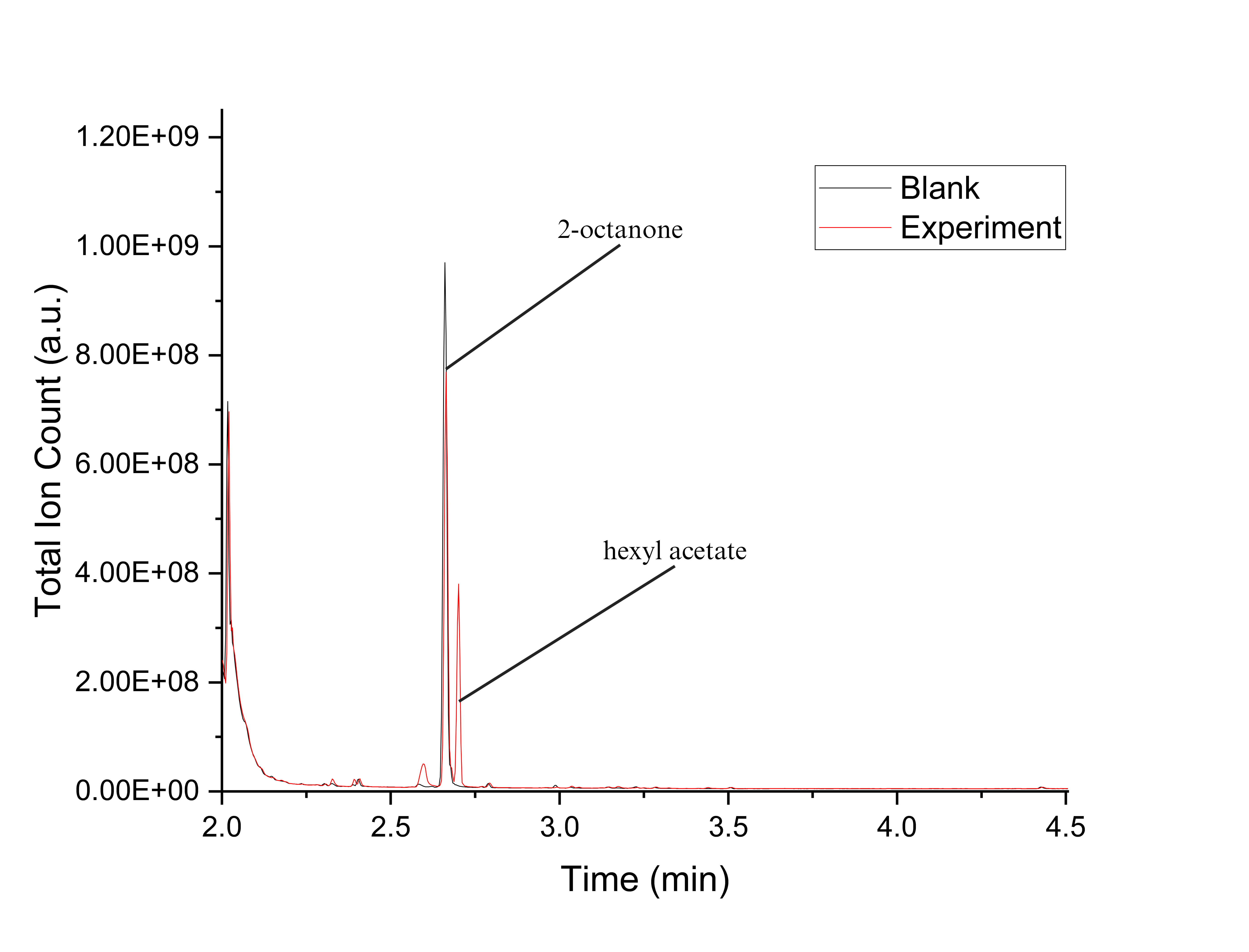

**Figure S24.** GCMS chromatogram for *ssn*BVMO reaction with 2-octanone.

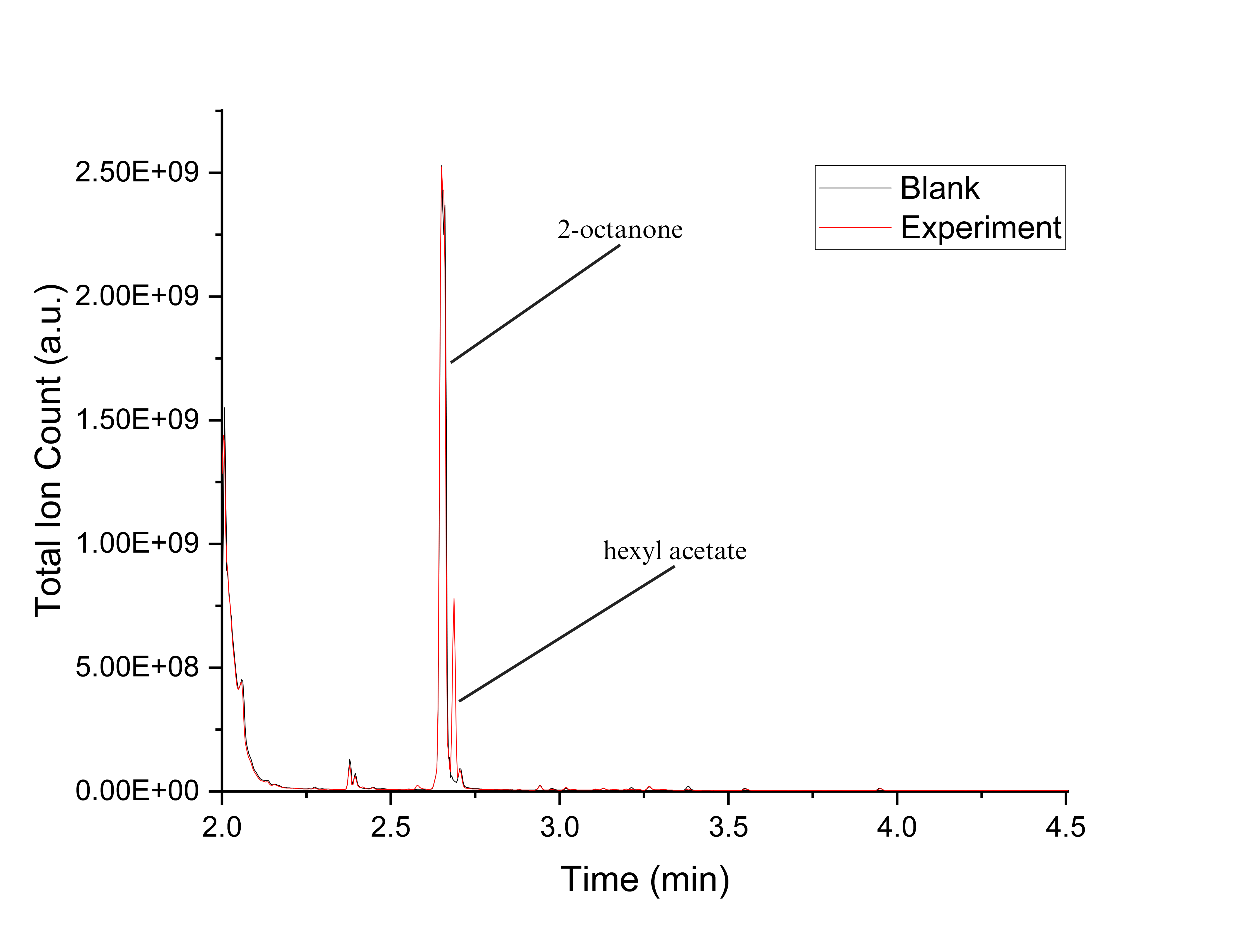

**Figure S25.** GCMS chromatogram for PTDH-L1-*ssn*BVMO reaction with 2-octanone.

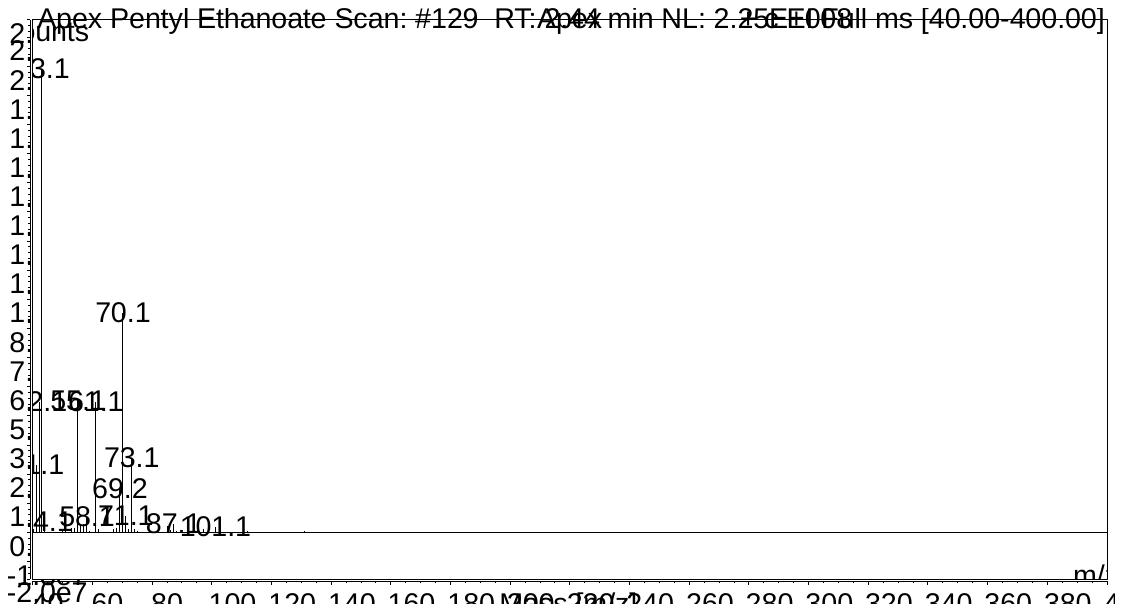

**Figure S26.** Fragmentation pattern for 2-octanone.

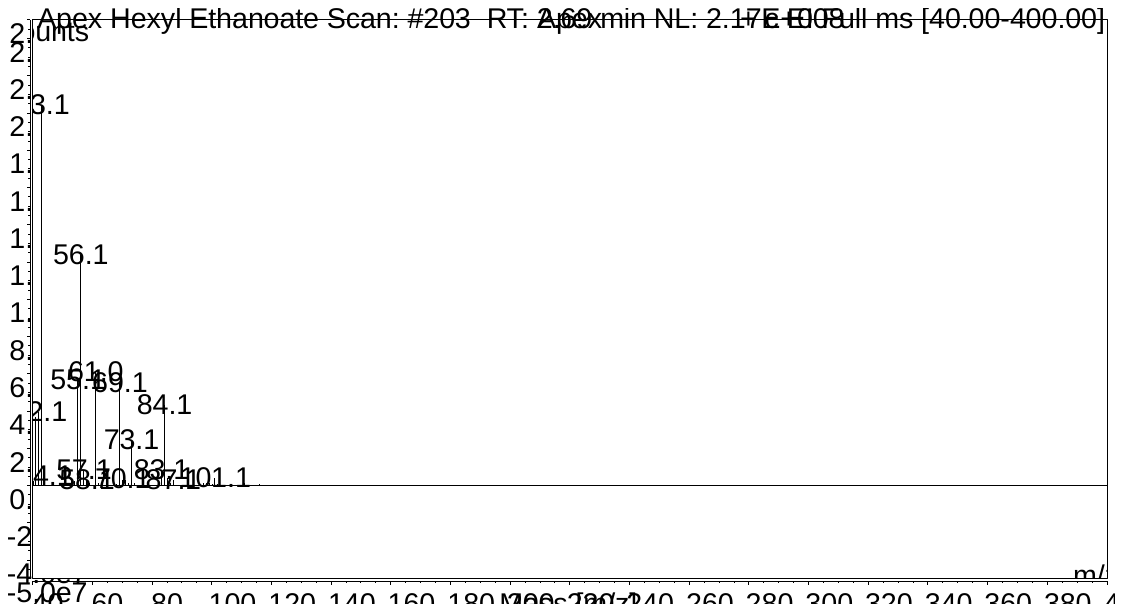

**Figure S27.** Fragmentation pattern for hexyl acetate.

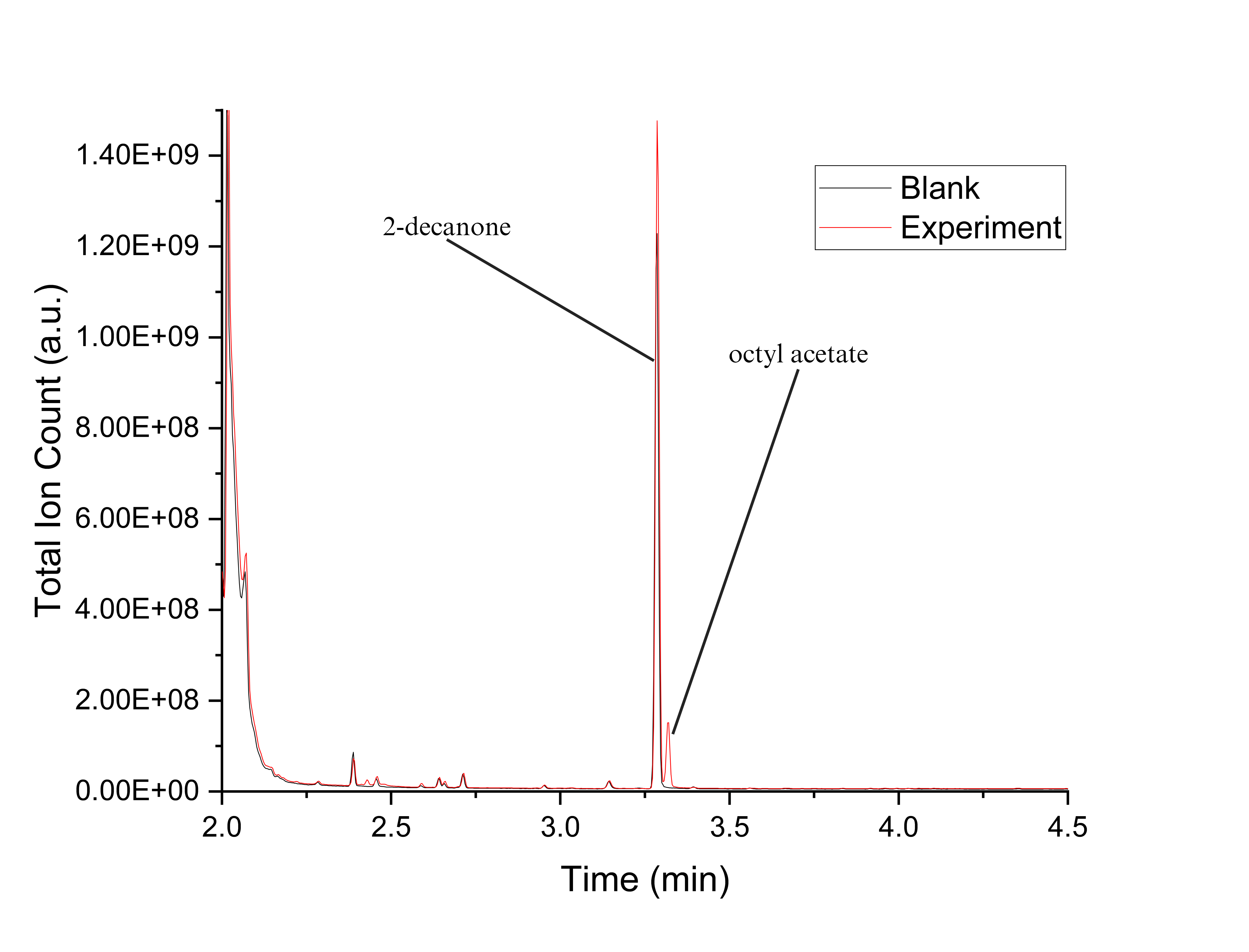

**Figure S28.** GCMS chromatogram for *ssn*BVMO reaction with 2-decanone.

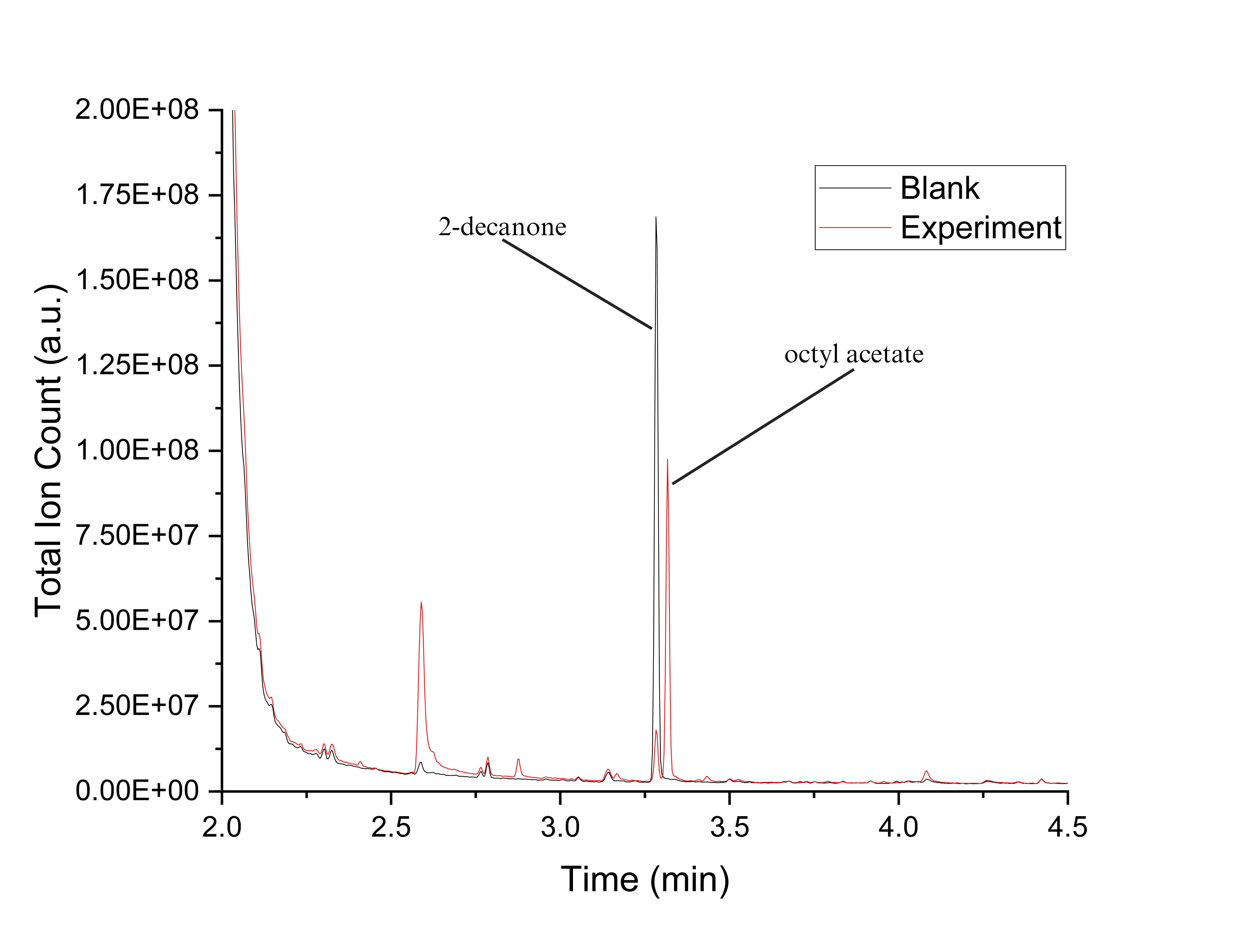

**Figure S29.** GCMS chromatogram for PTDH-L1-*ssn*BVMO reaction with 2-decanone.

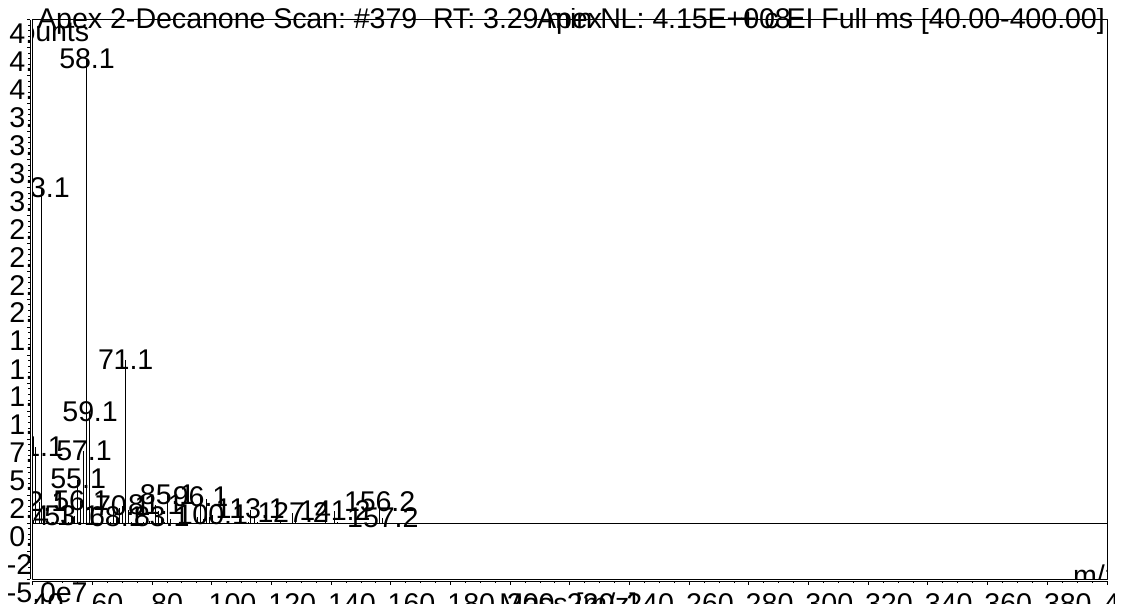

**Figure S30.** Fragmentation pattern for 2-decanone.

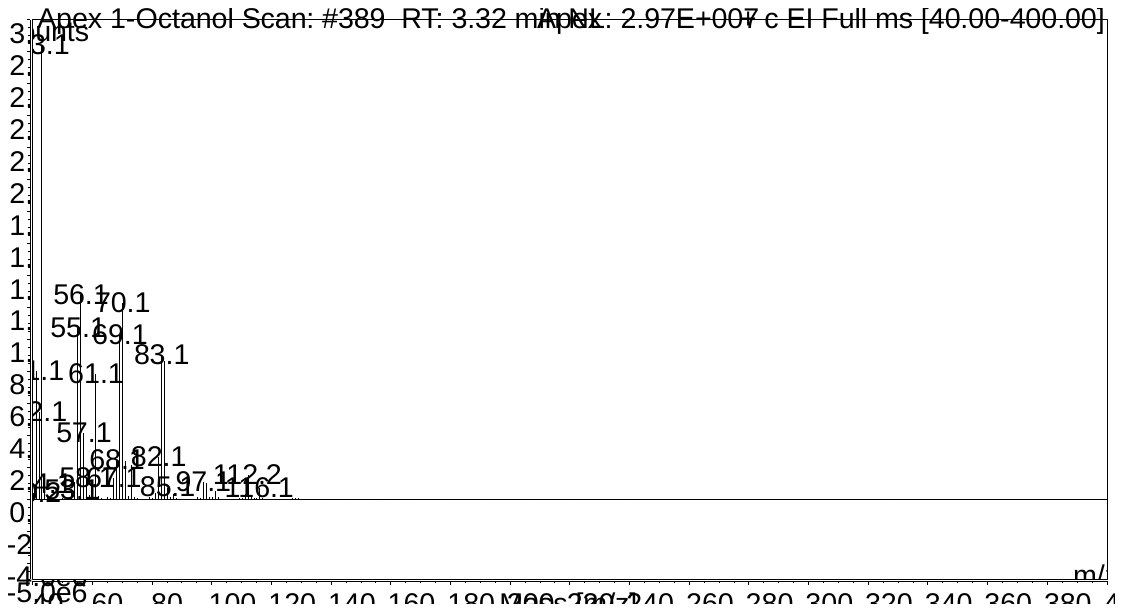

**Figure S31.** Fragmentation pattern for octyl acetate.

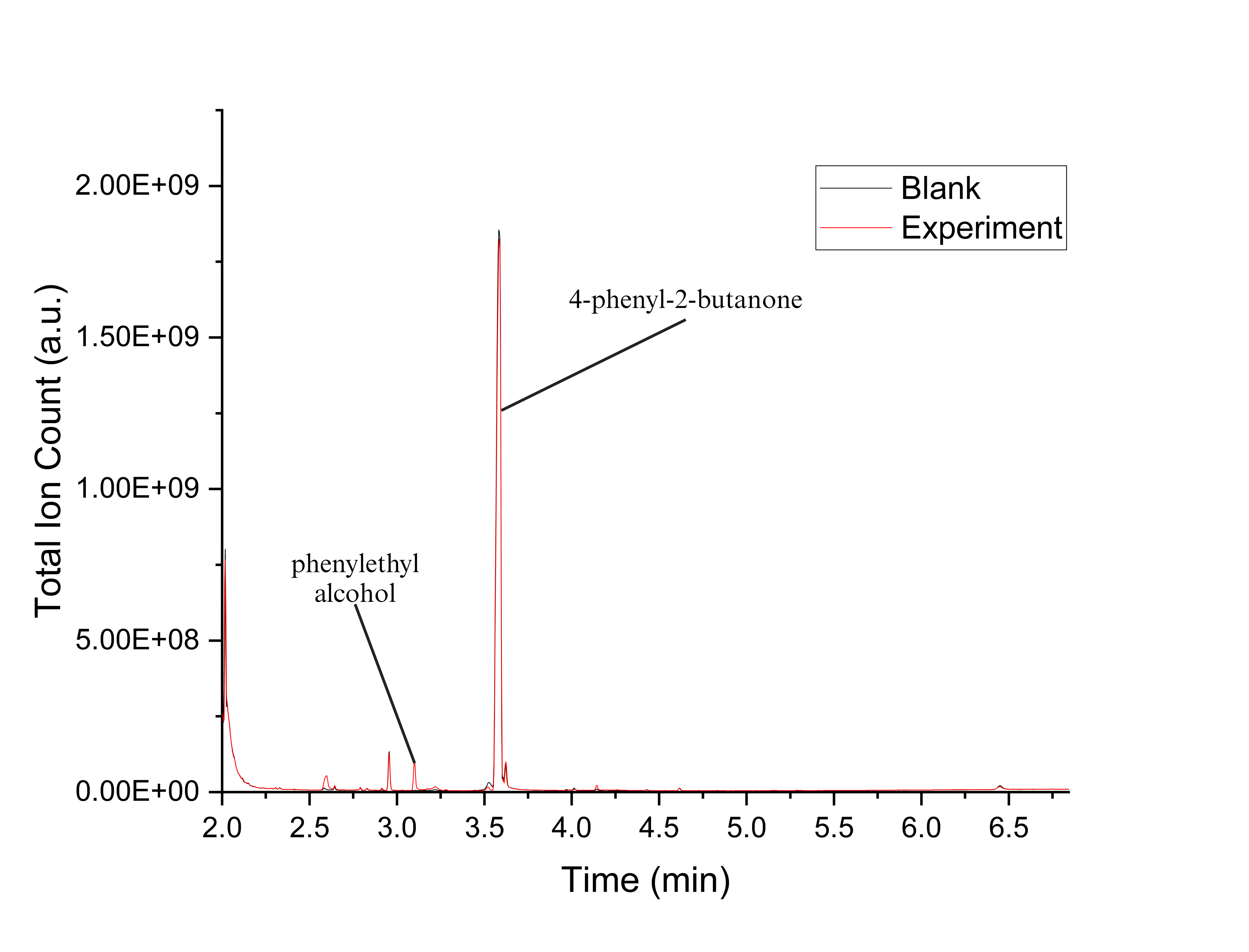

**Figure S32.** GCMS chromatogram for *ssn*BVMO reaction with 4-phenyl-2-butanone.

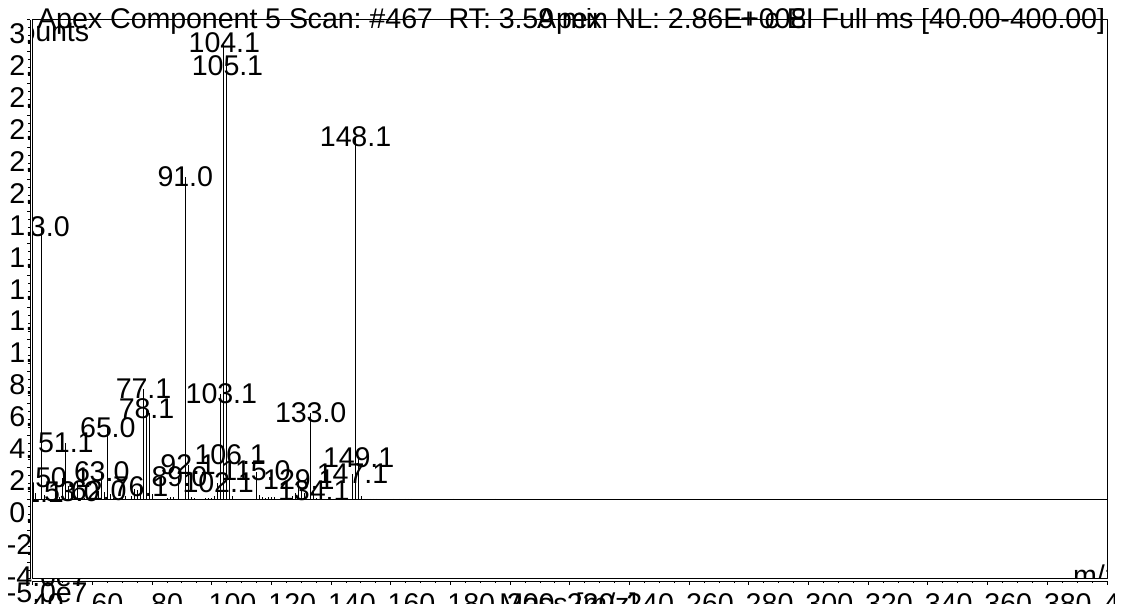

**Figure S33.** Fragmentation pattern for 4-phenyl-2-butanone.

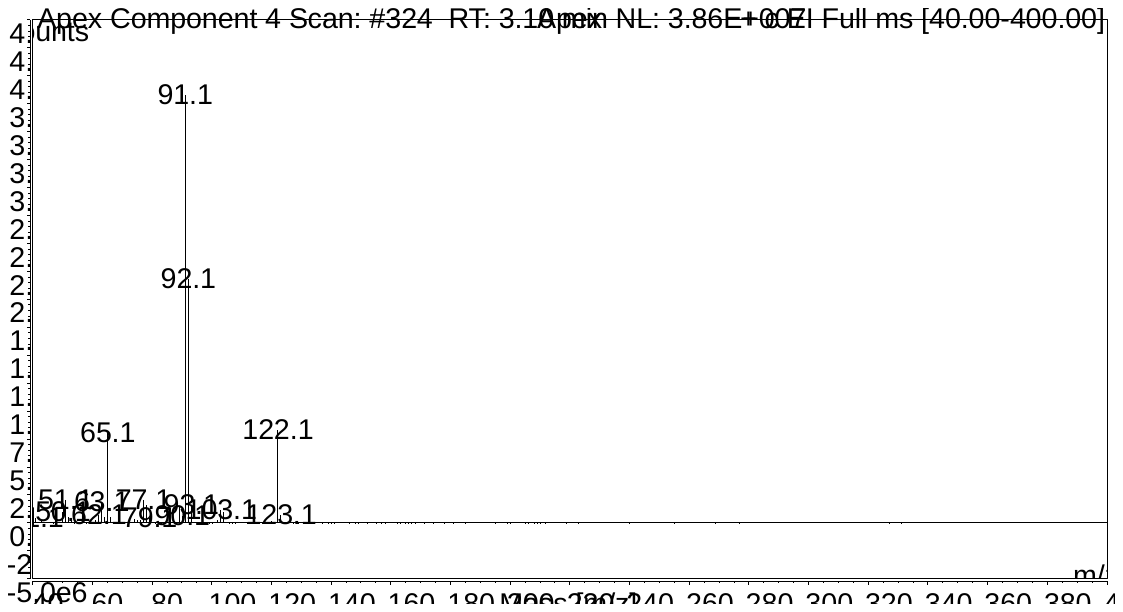

**Figure S34.** Fragmentation pattern for phenyl ethyl alcohol.

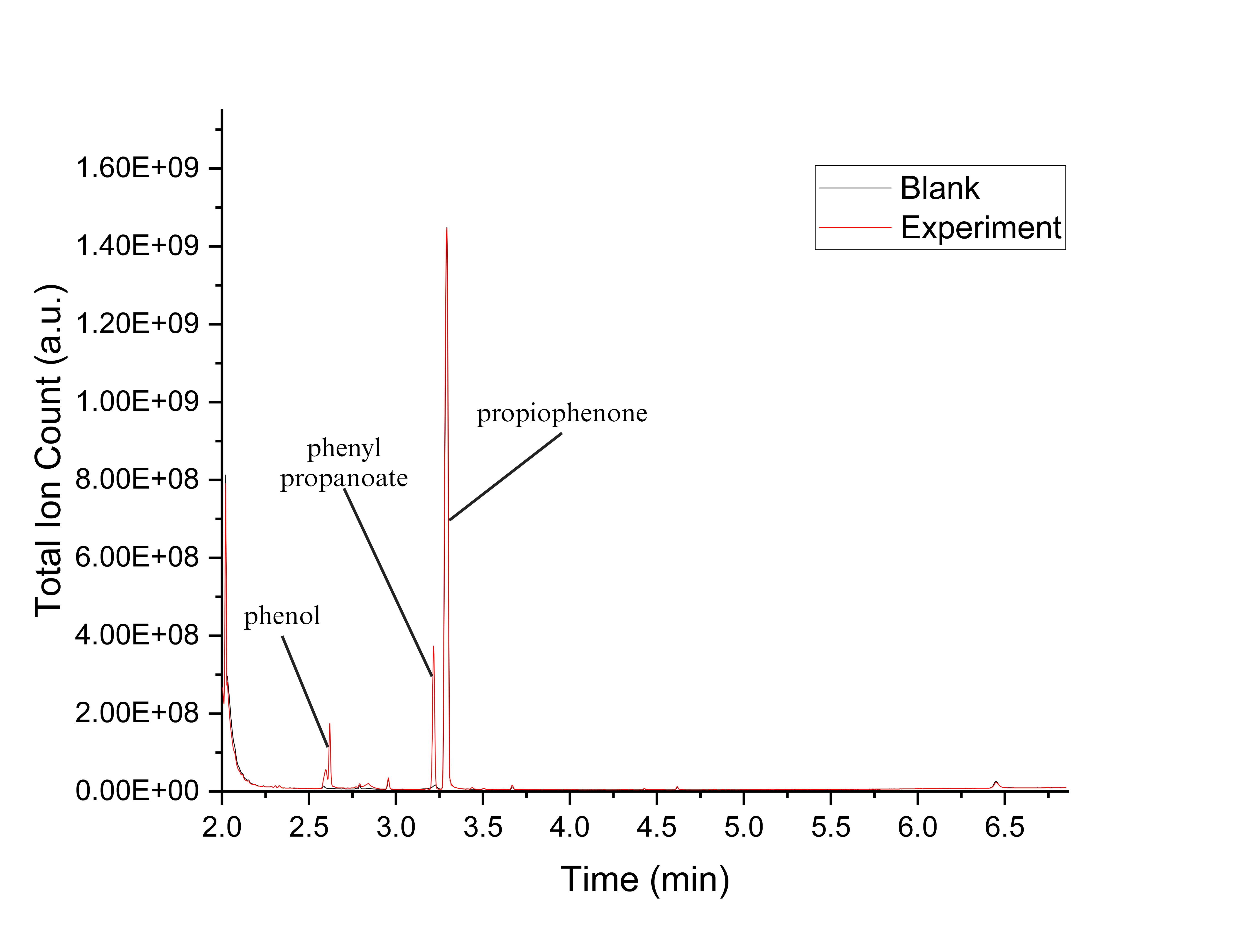

**Figure S35.** GCMS chromatogram for *ssn*BVMO reaction with propiophenone.

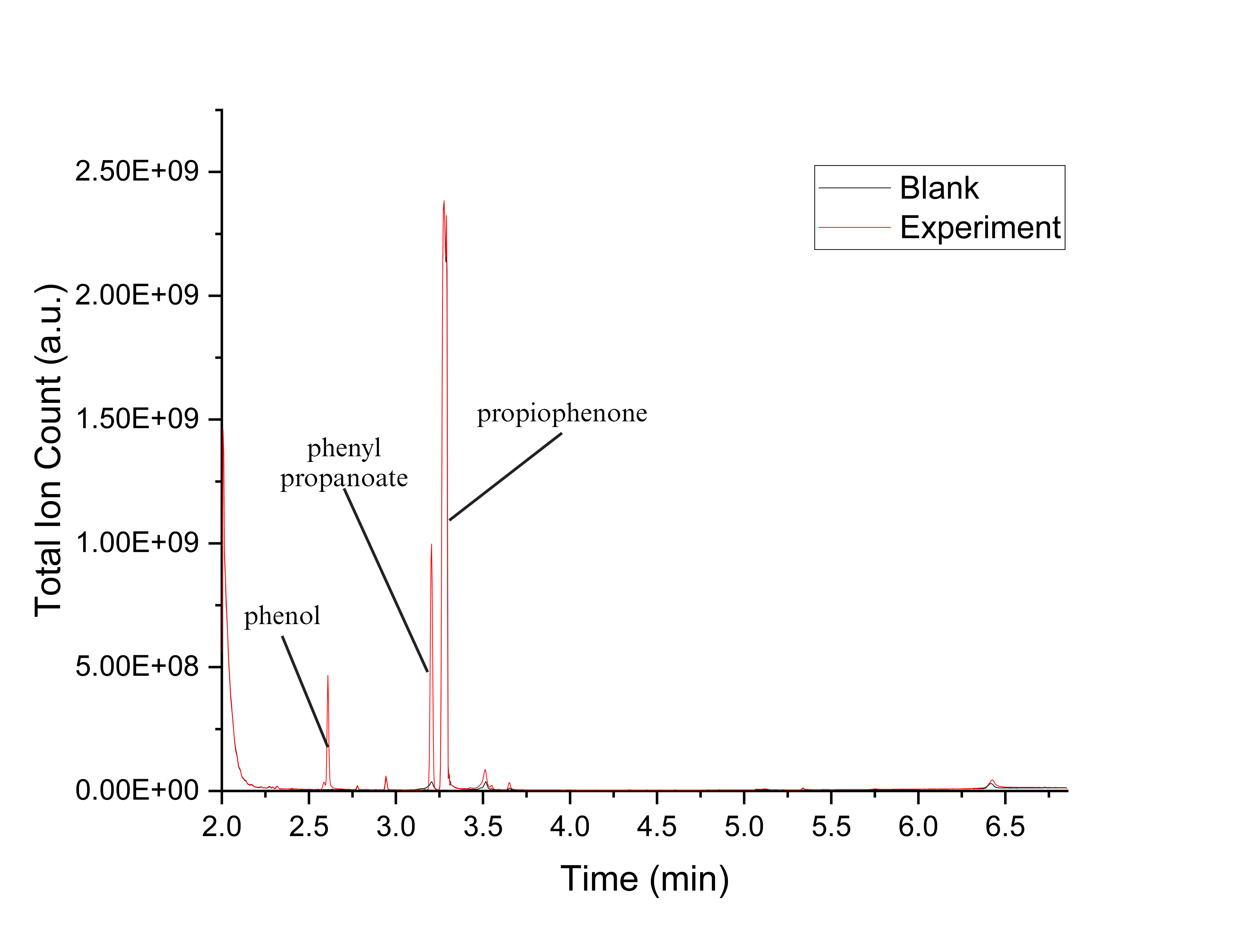

**Figure S36.** GCMS chromatogram for PTDH-L1-*ssn*BVMO reaction with propiophenone.

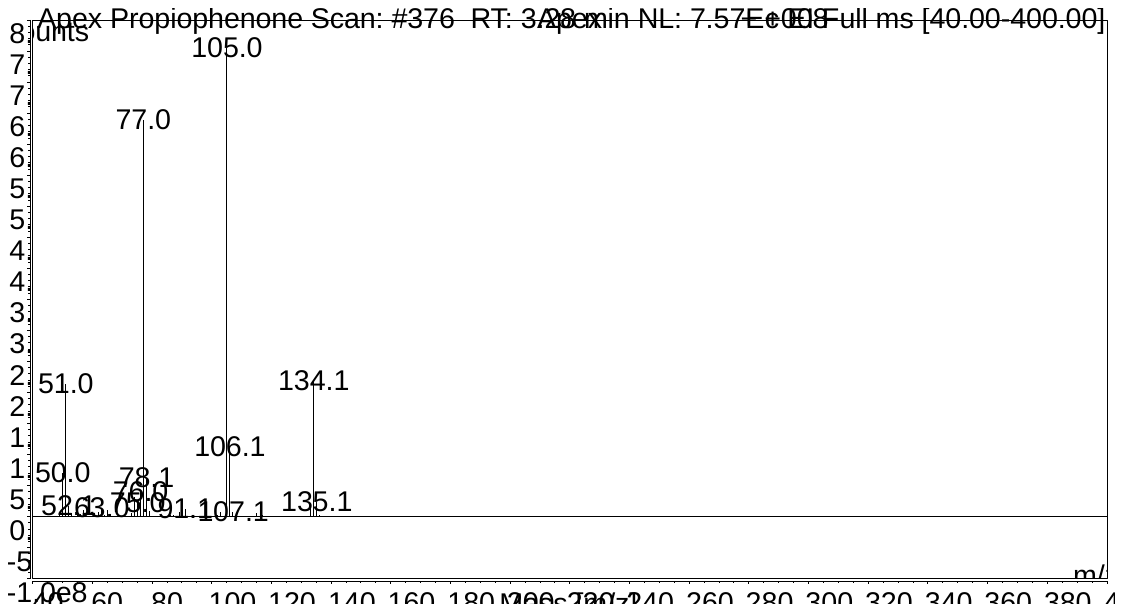

**Figure S37.** Fragmentation pattern for propiophenone.

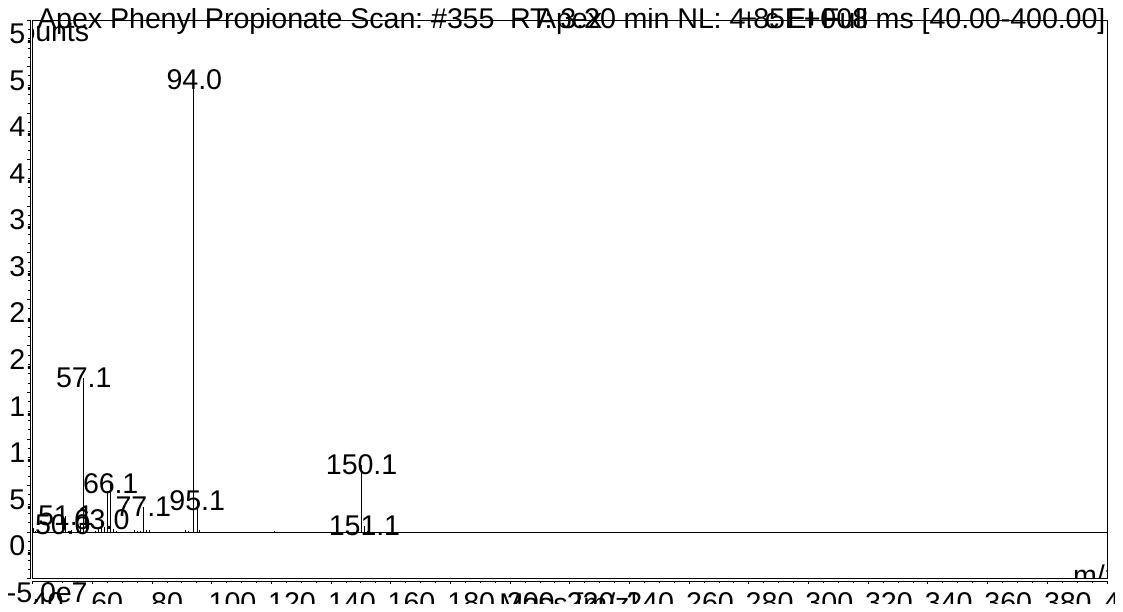

**Figure S38.** Fragmentation for phenyl propanoate

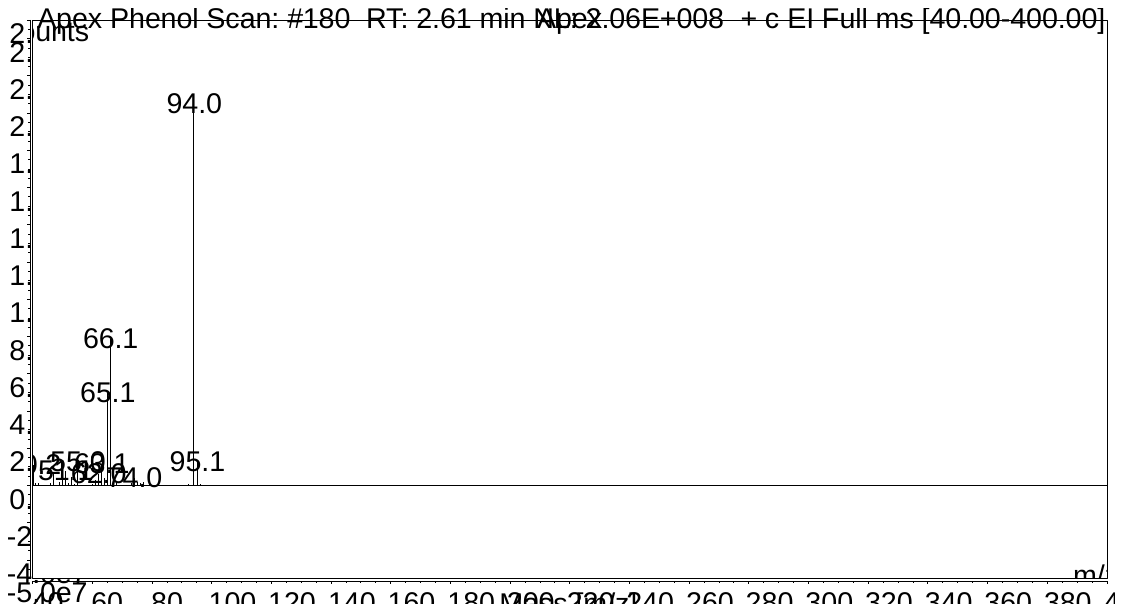

**Figure S39.** Fragmentation pattern for phenol.

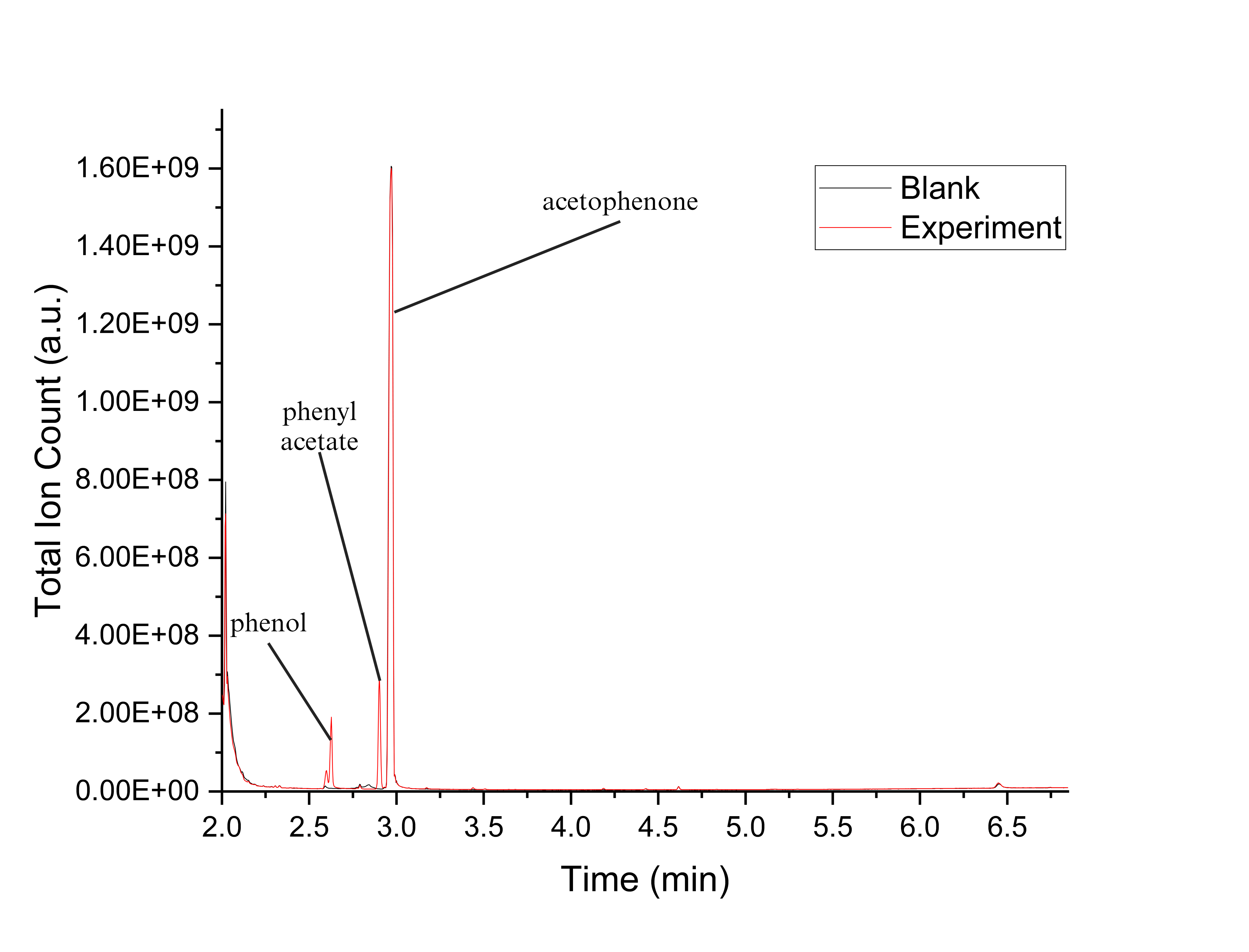

**Figure S40.** GCMS chromatogram for *ssn*BVMO reaction with acetophenone.

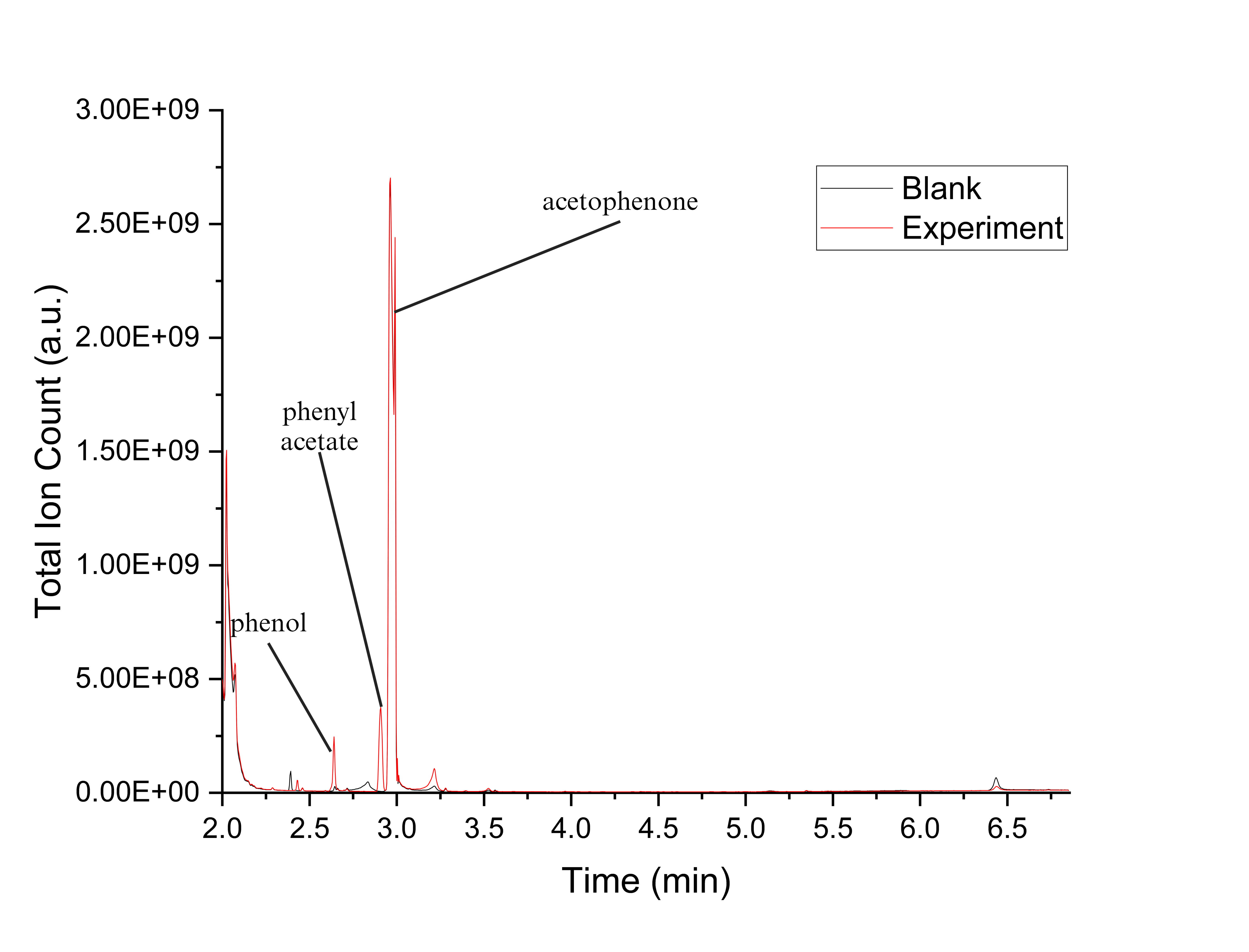

**Figure S41.** GCMS chromatogram for PTDH-L1-*ssn*BVMO reaction with acetophenone.

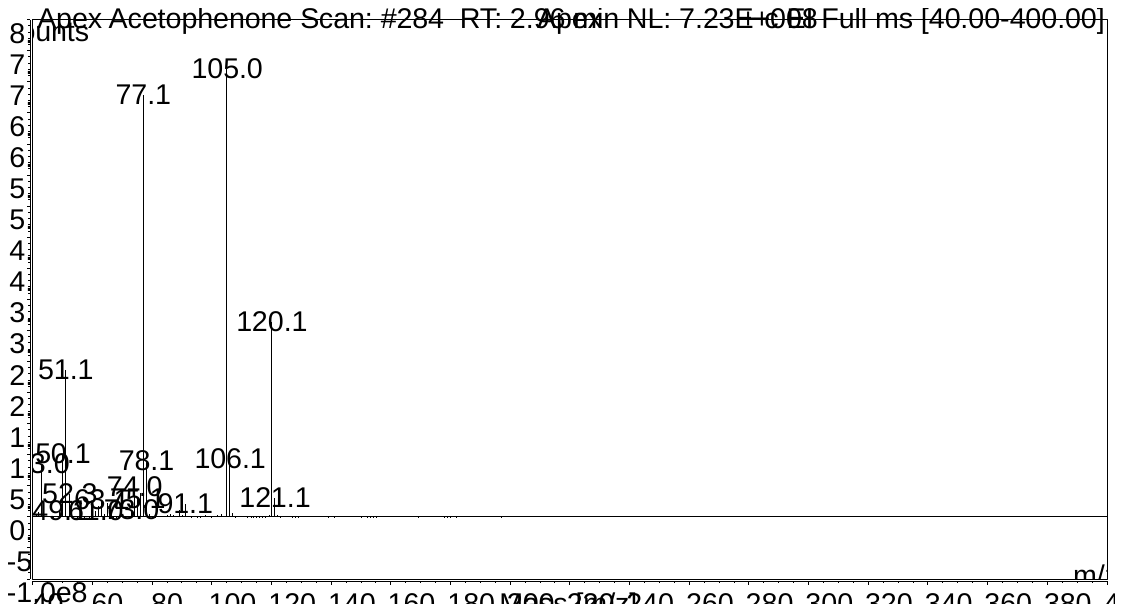

**Figure S42.** Fragmentation pattern for acetophenone.

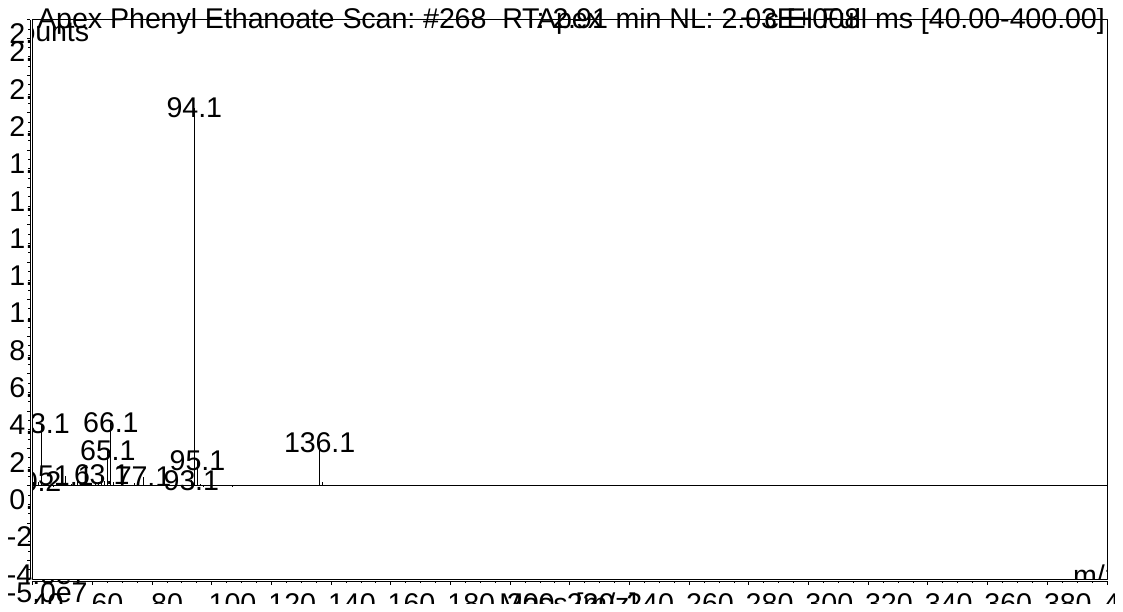

**Figure S43.** Fragmentation pattern phenyl acetate.

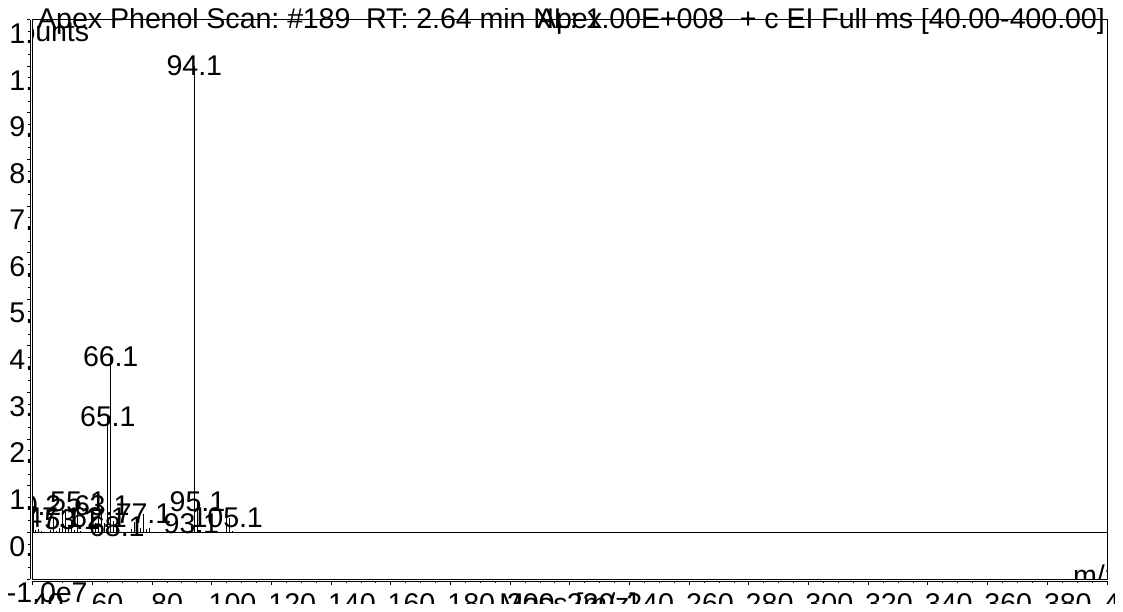

**Figure S44.** Fragmentation pattern for phenol.

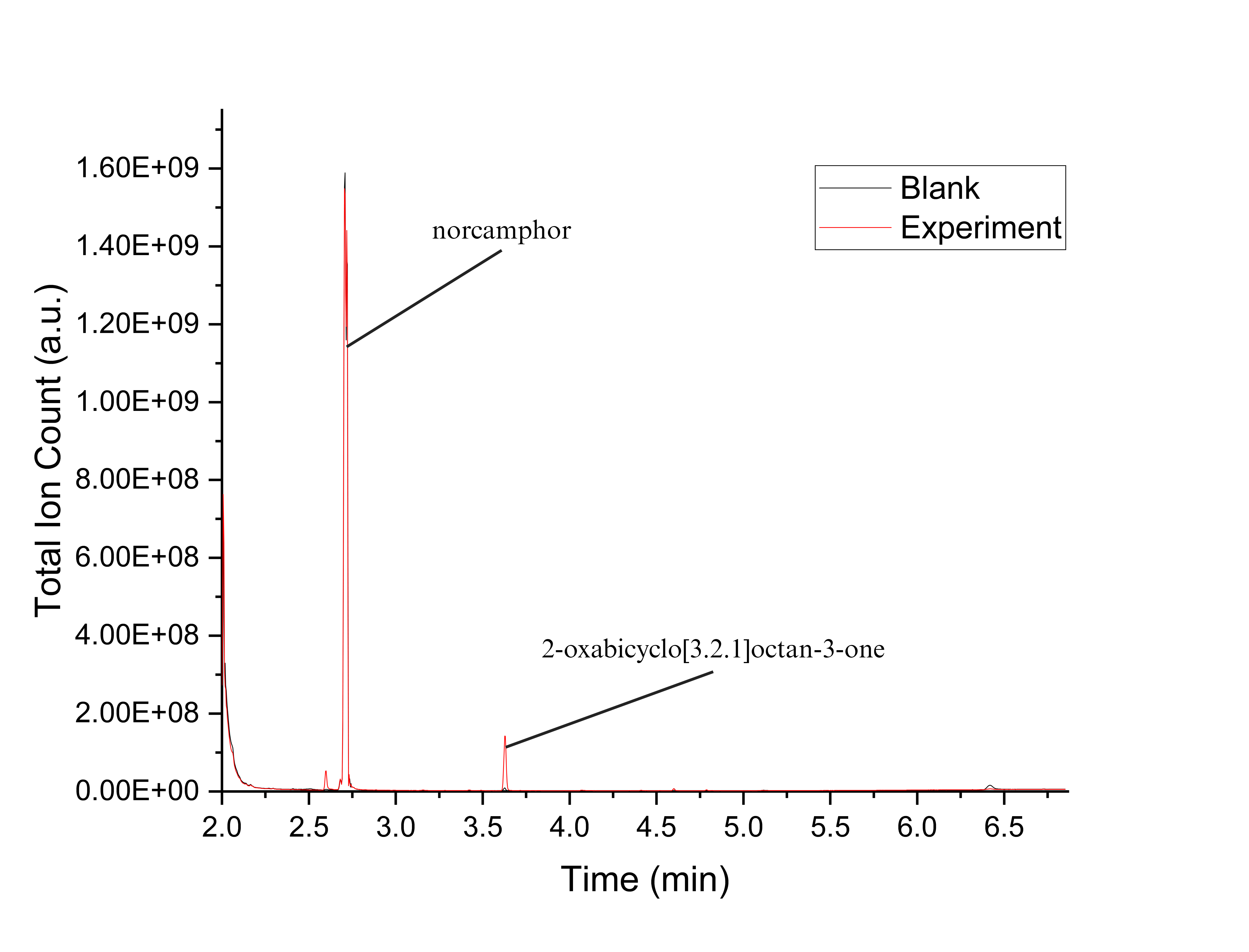

**Figure S45.** GCMS chromatogram for *ssn*BVMO reaction with norcamphor.

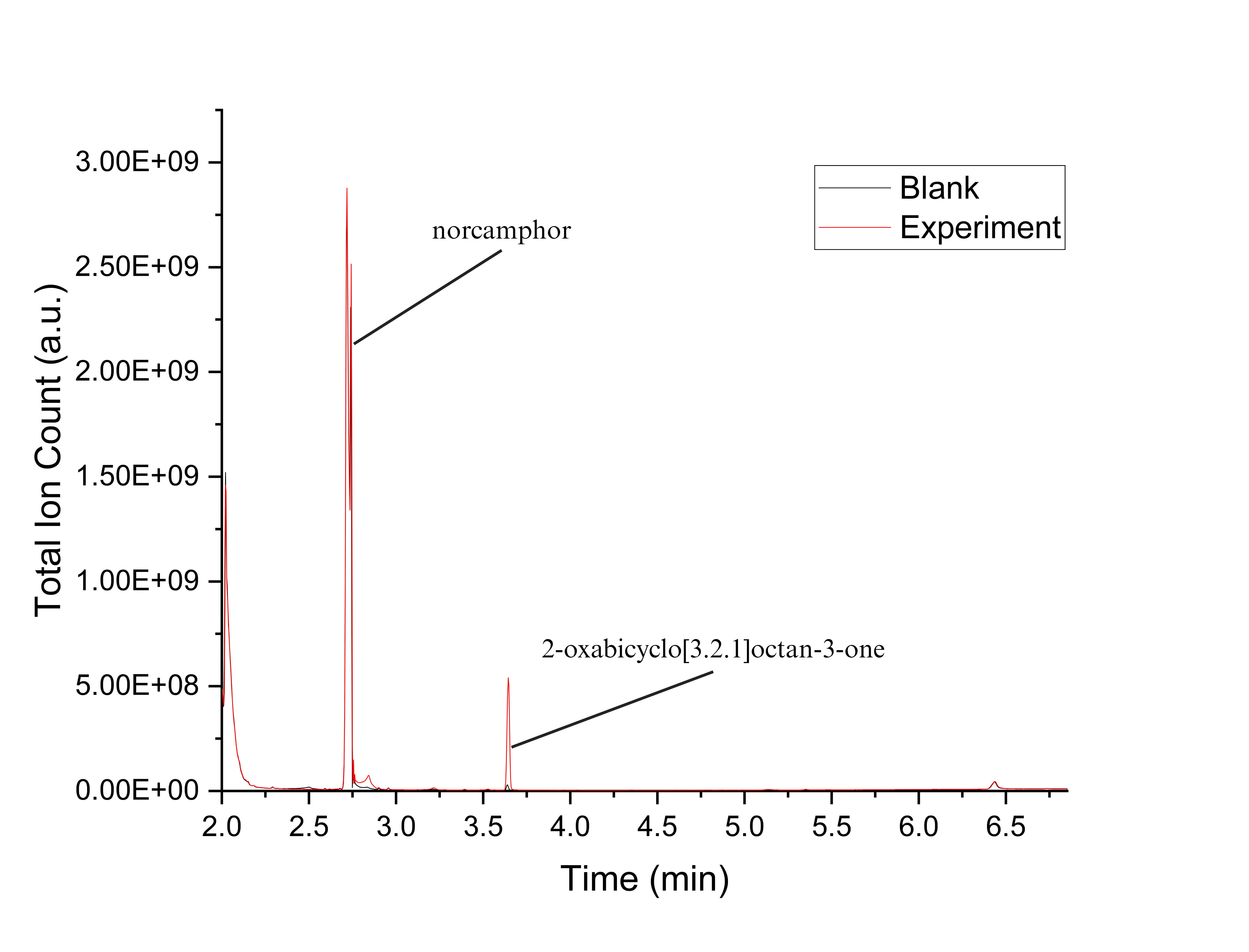

**Figure S46.** GCMS chromatogram for PTDH-L1-*ssn*BVMO reaction with norcamphor.

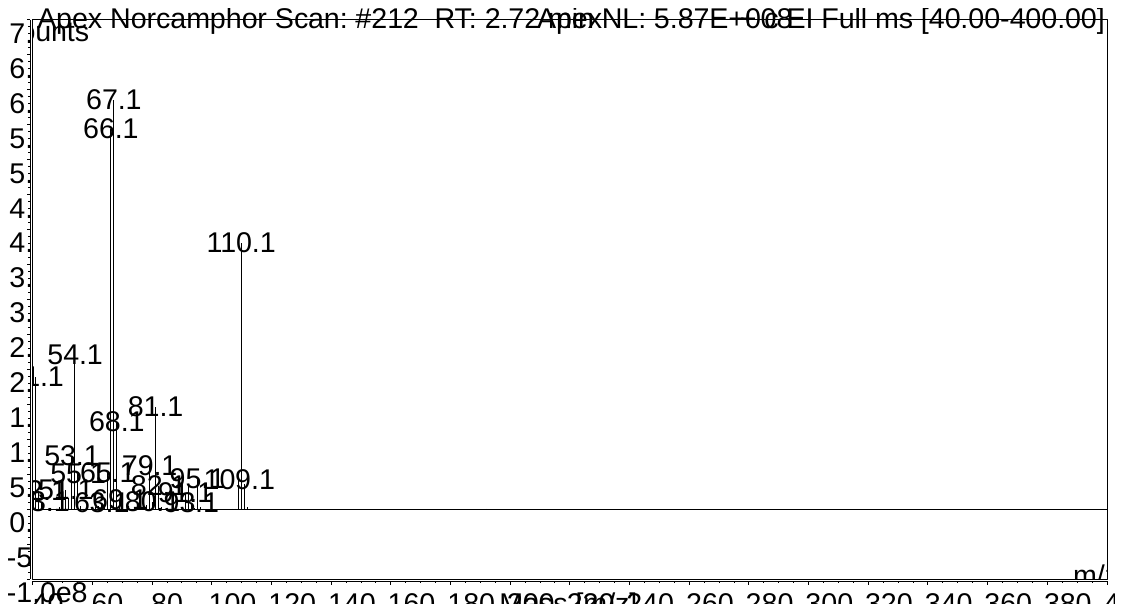

**Figure S47.** Fragmentation pattern for norcamphor.

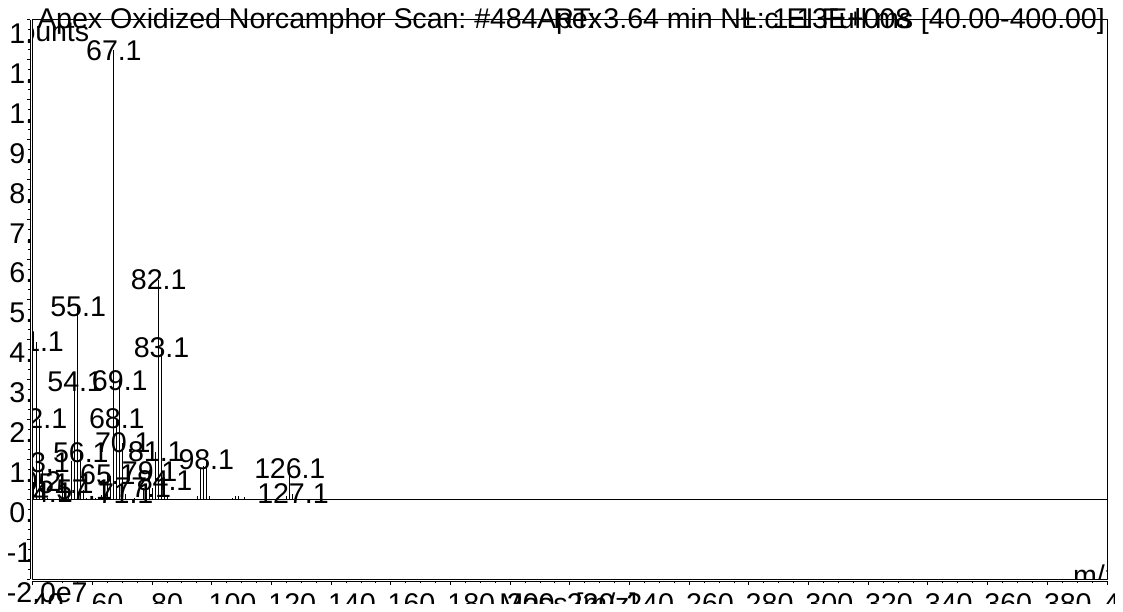

**Figure S48.** Fragmentation pattern for 2-oxabicyclo[3.2.1]octan-3-one.

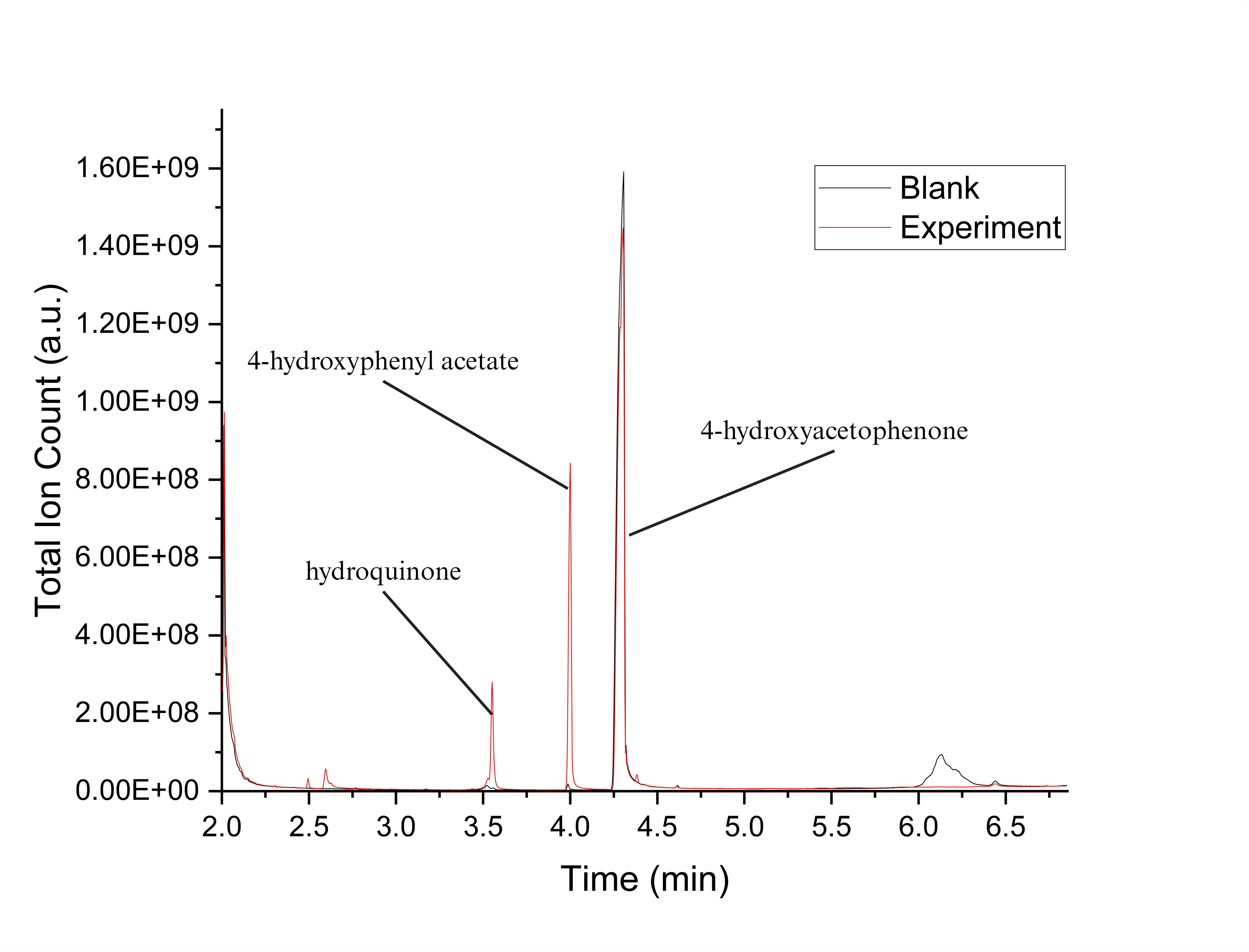

**Figure S49.** GCMS chromatogram for *ssn*BVMO reaction with 4-hydroxyacetophenone.

.

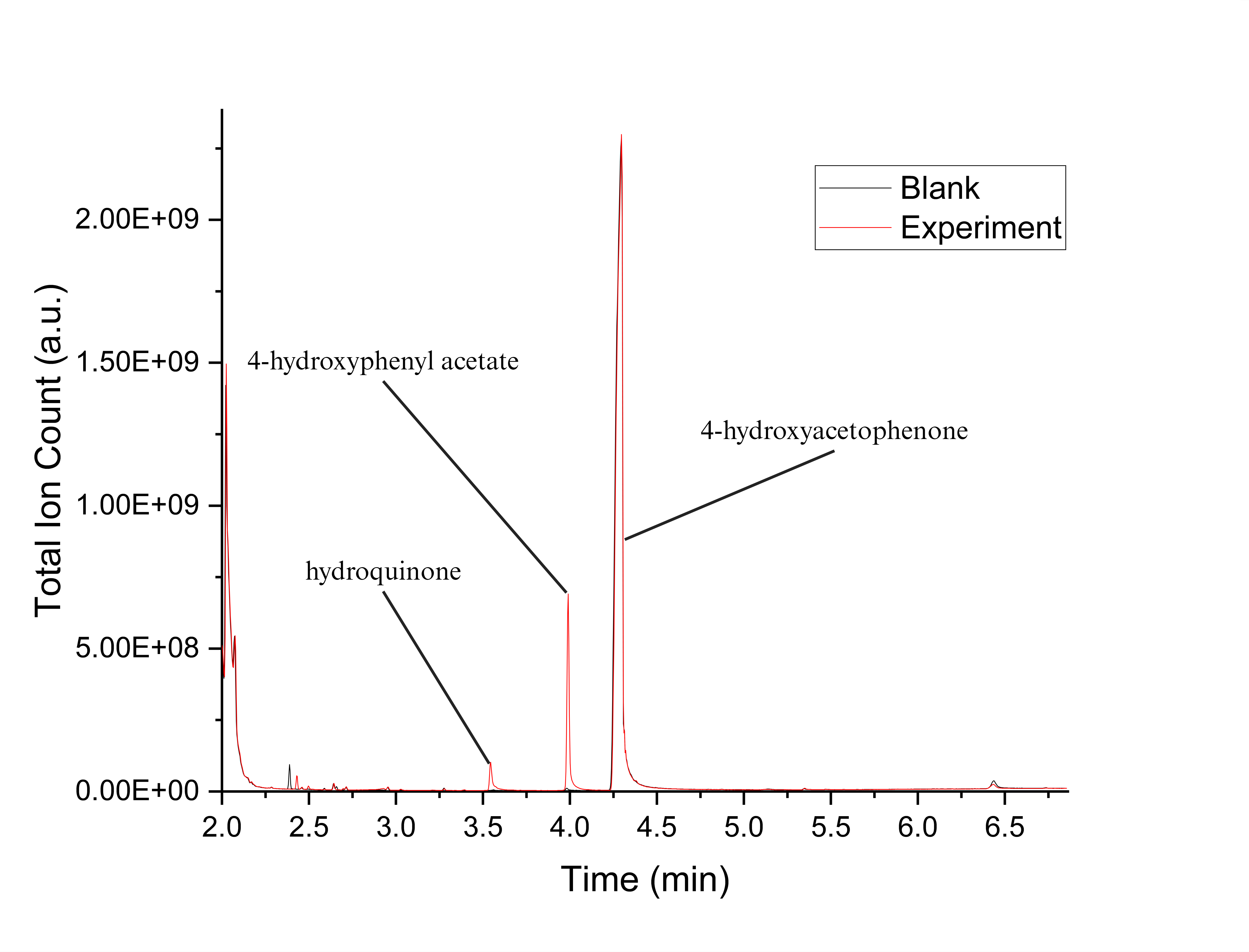

**Figure S50.** GCMS chromatogram for PTDH-L1-*ssn*BVMO reaction with 4-hydroxyacetophenone.

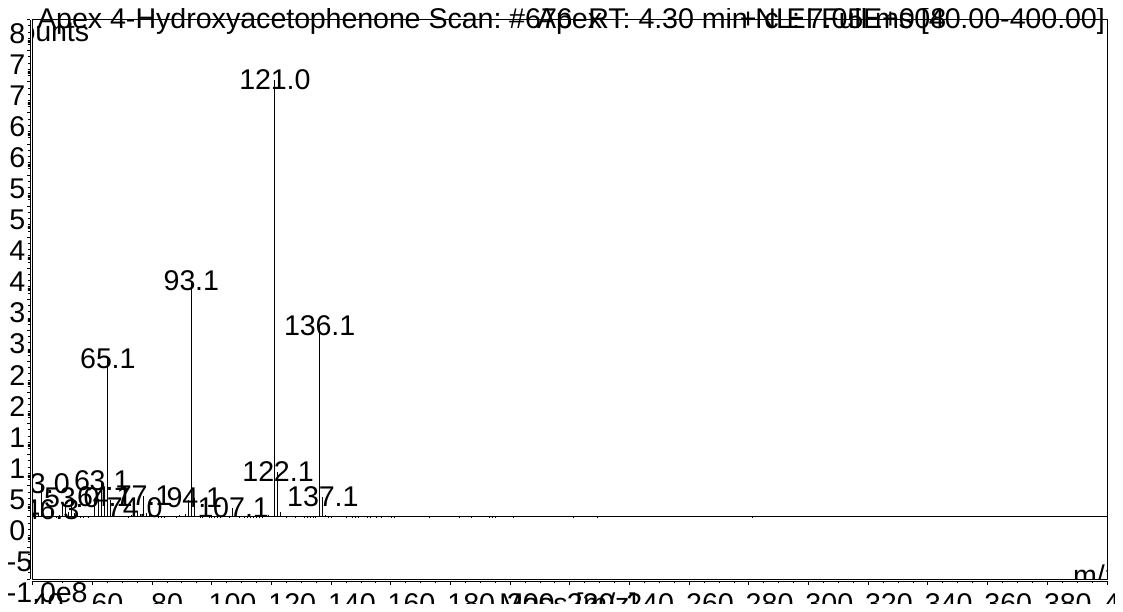

**Figure S51.** Fragmentation pattern for 4-hydroxyacetophenone.

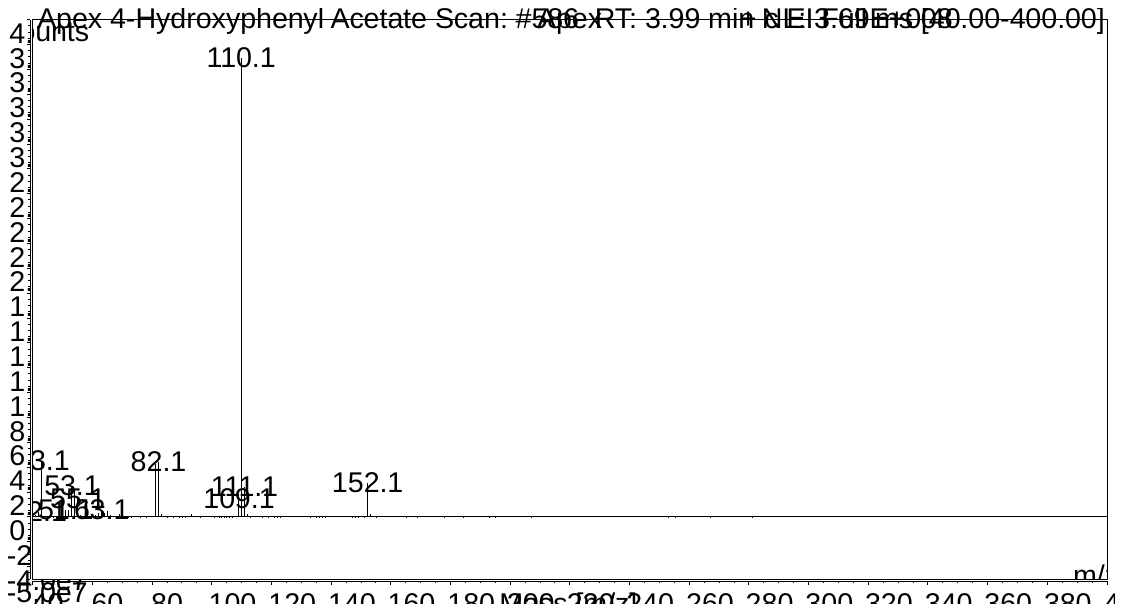

**Figure S52.** Fragmentation pattern for 4-hydroxyphenyl acetate.

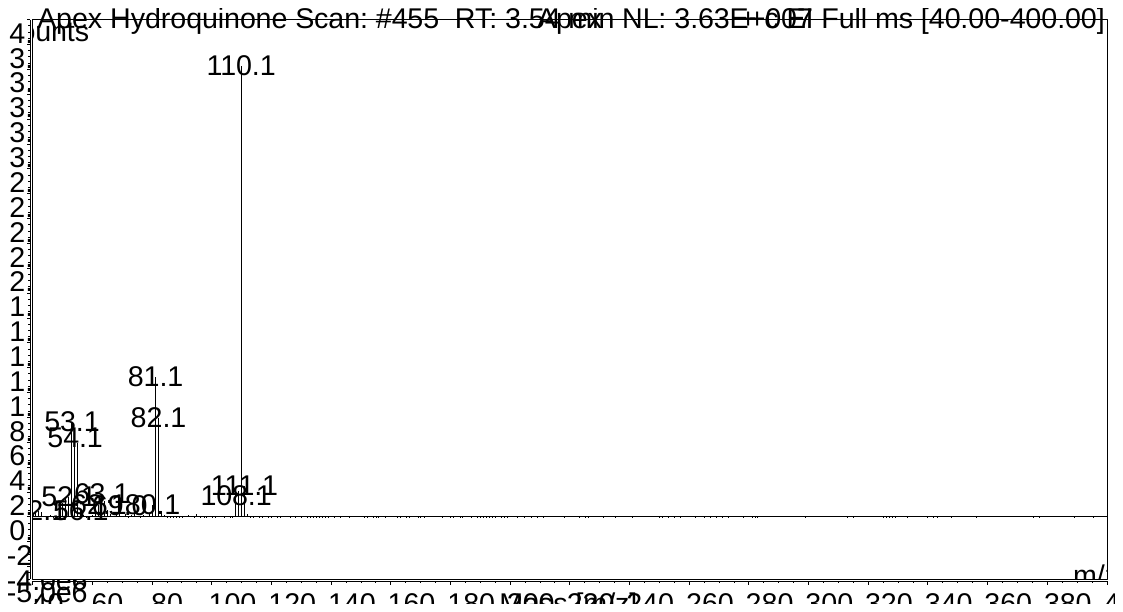

**Figure S53.** Fragmentation pattern for hydroquinone.

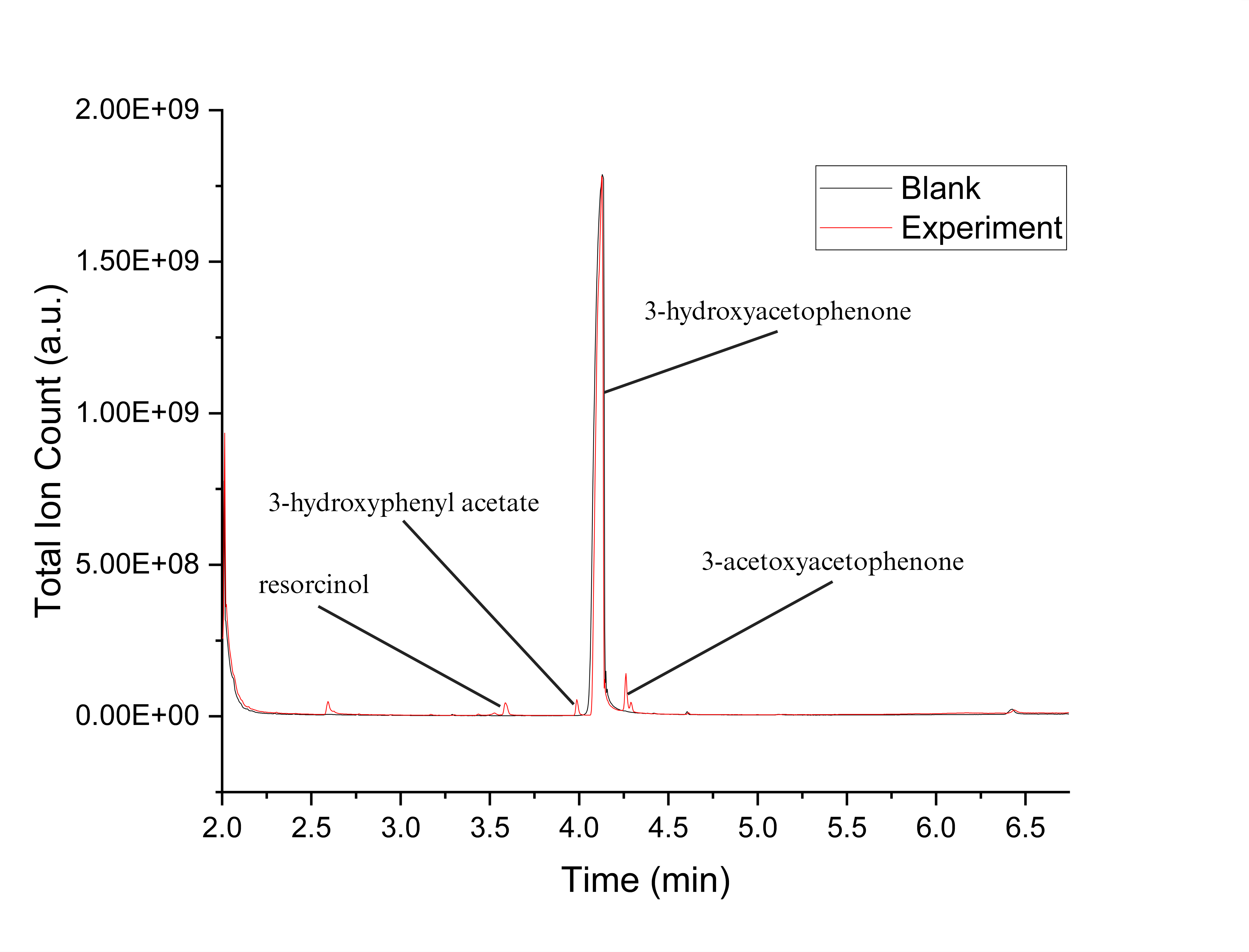

**Figure S54.** GCMS chromatogram for *ssn*BVMO reaction with 3-hydroxyacetophenone.

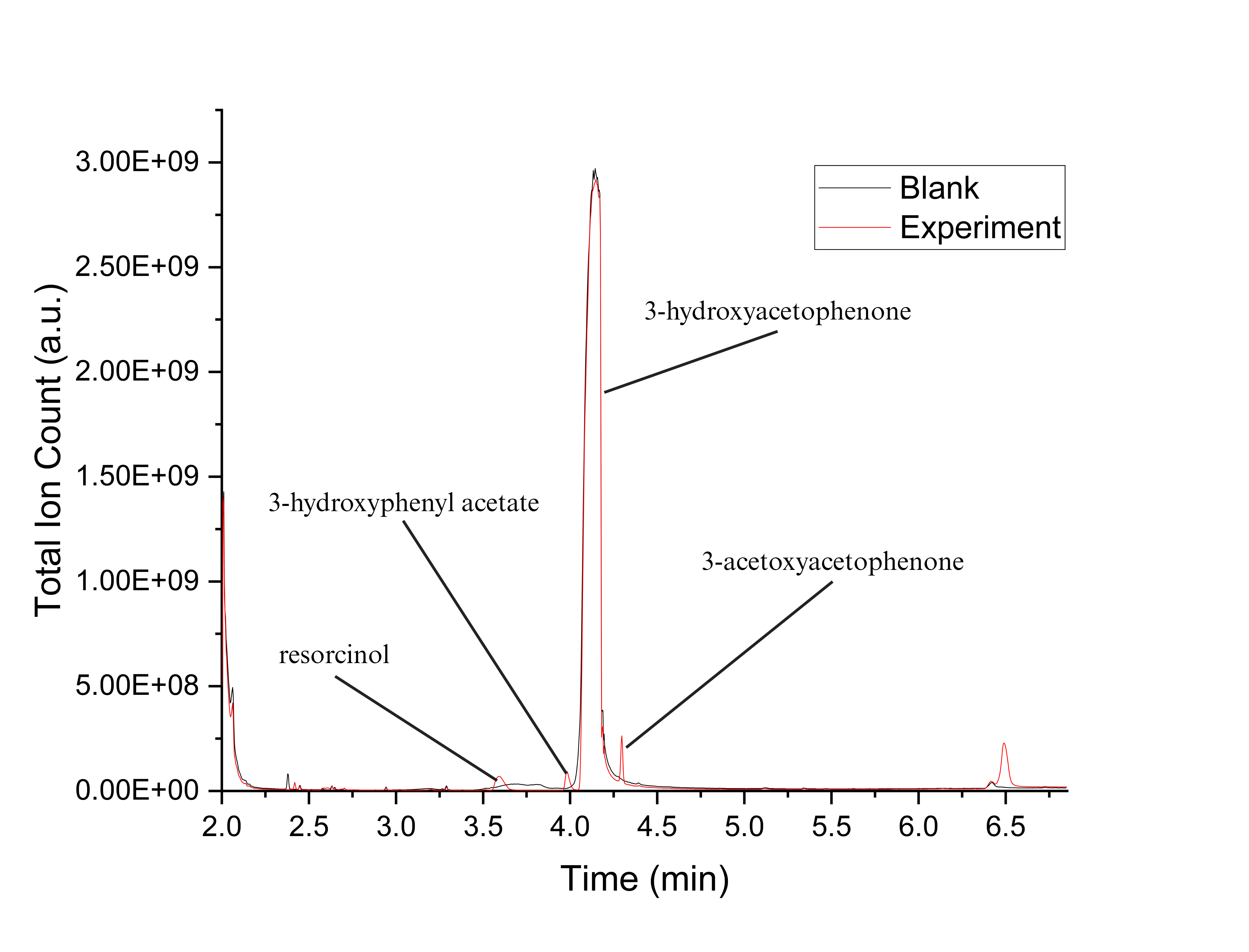

**Figure S55.** GCMS chromatogram for PTDH-L1-*ssn*BVMO reaction with 3-hydroxyacetophenone.

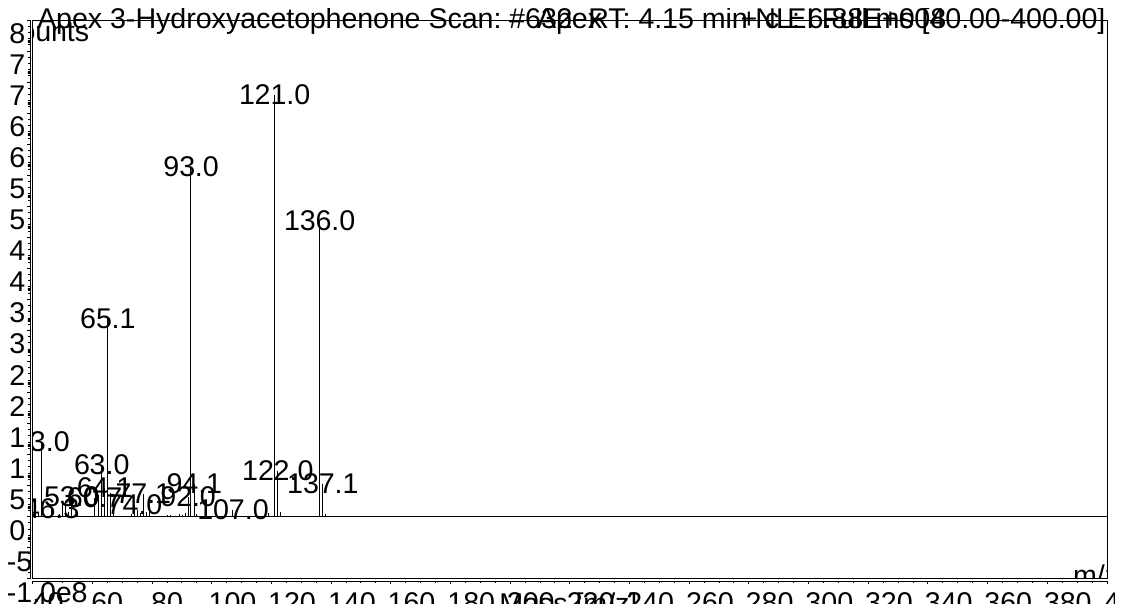

**Figure S56.** Fragmentation pattern for 3-hydroxyacetophenone.

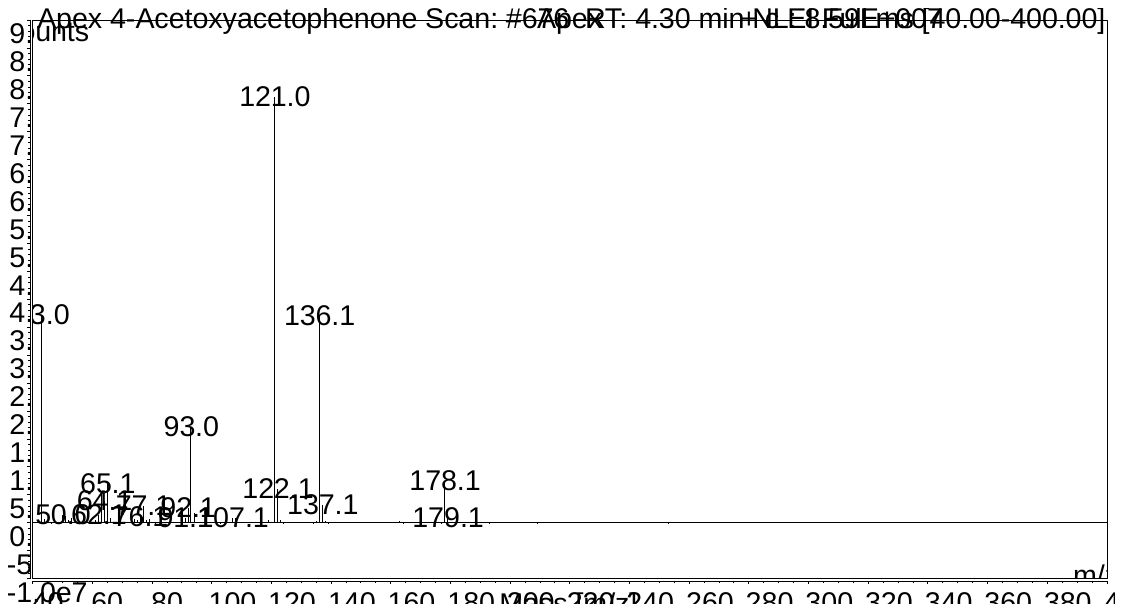

**Figure S57.** Fragmentation pattern for 3-acetoxyacetophenone, most likely formed through nucleophilic attack of acetyl group by the alcohol group.

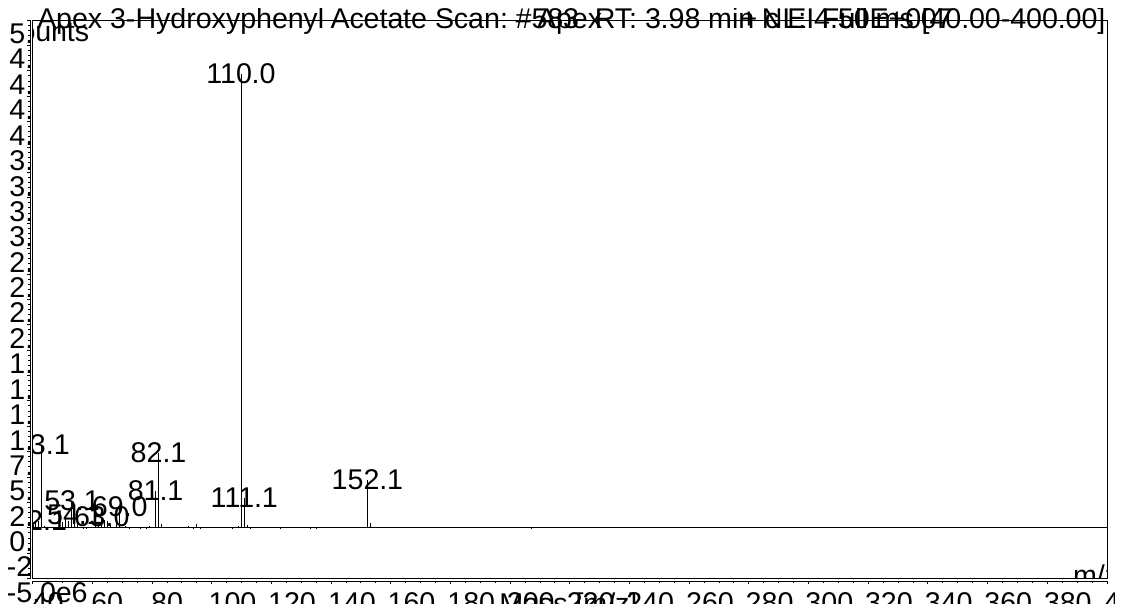

**Figure S58.** Fragmentation pattern for 3-hydroxylphenyl acetate.

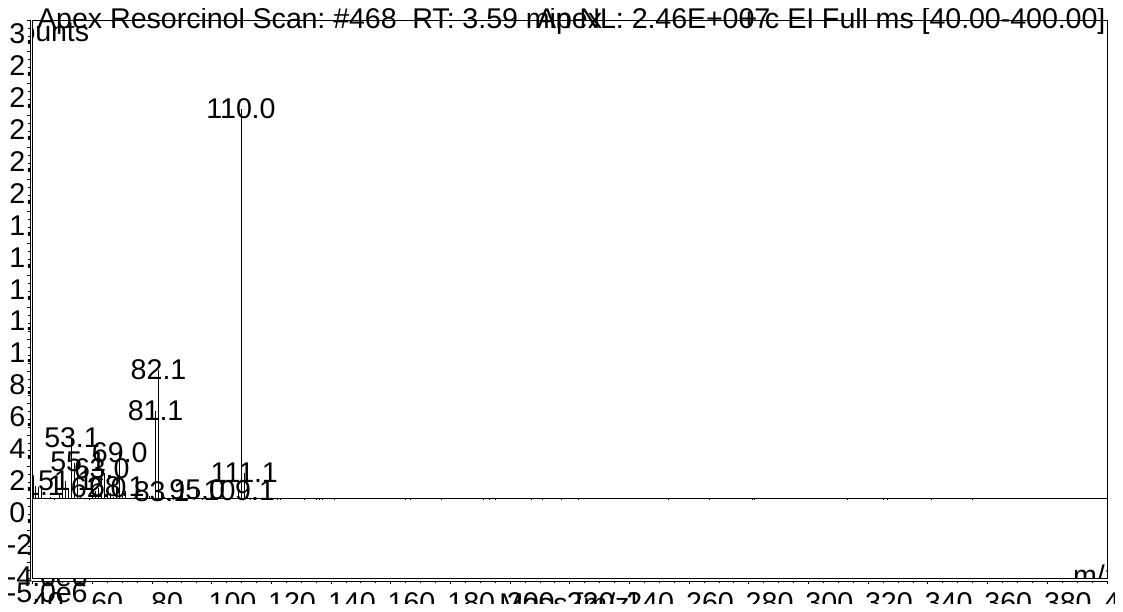

**Figure S59.** Fragmentation pattern for resorcinol.

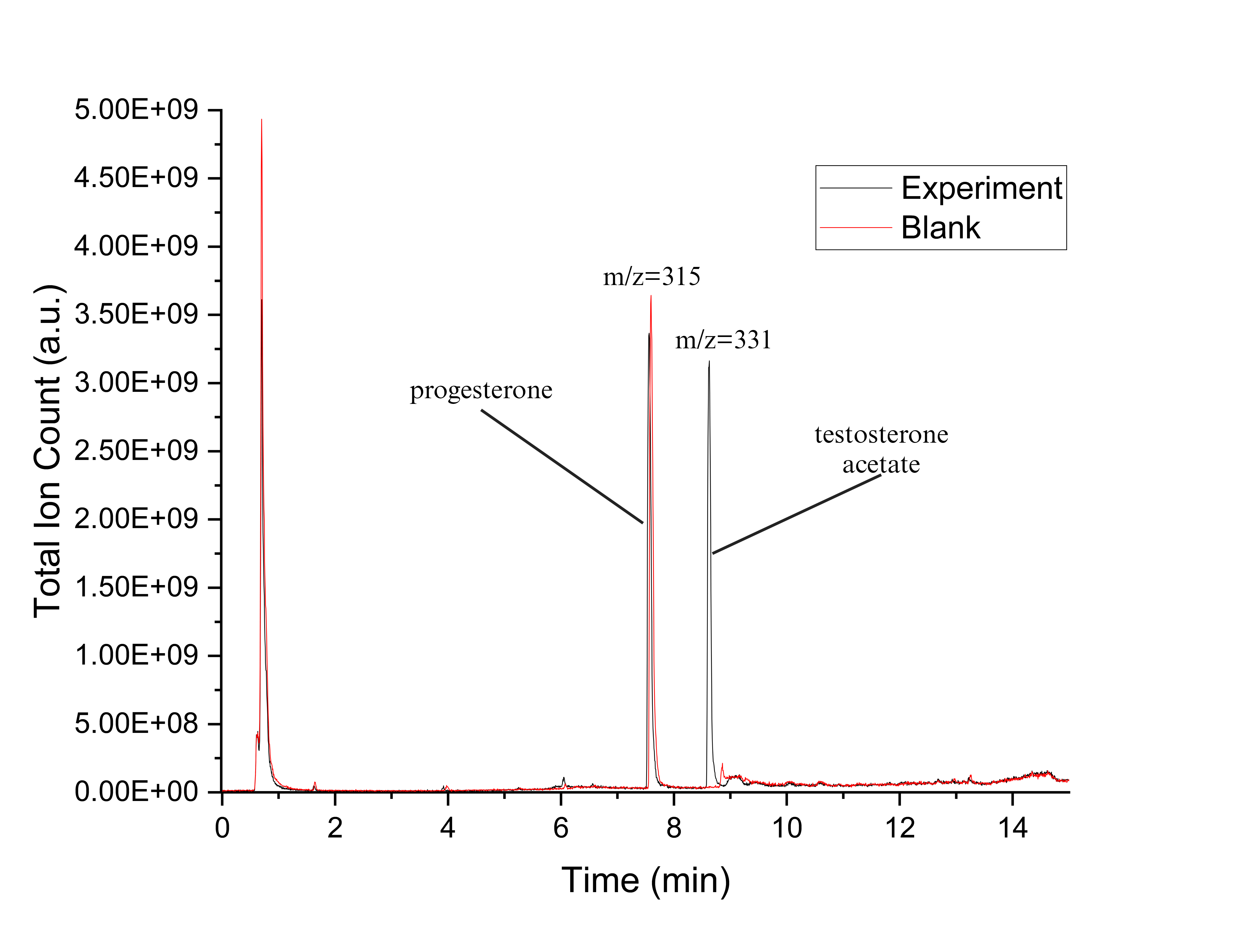

**Figure S60.** UPLC chromatogram for *ssn*BVMO reaction with progesterone. ESI positive ion mode was used.

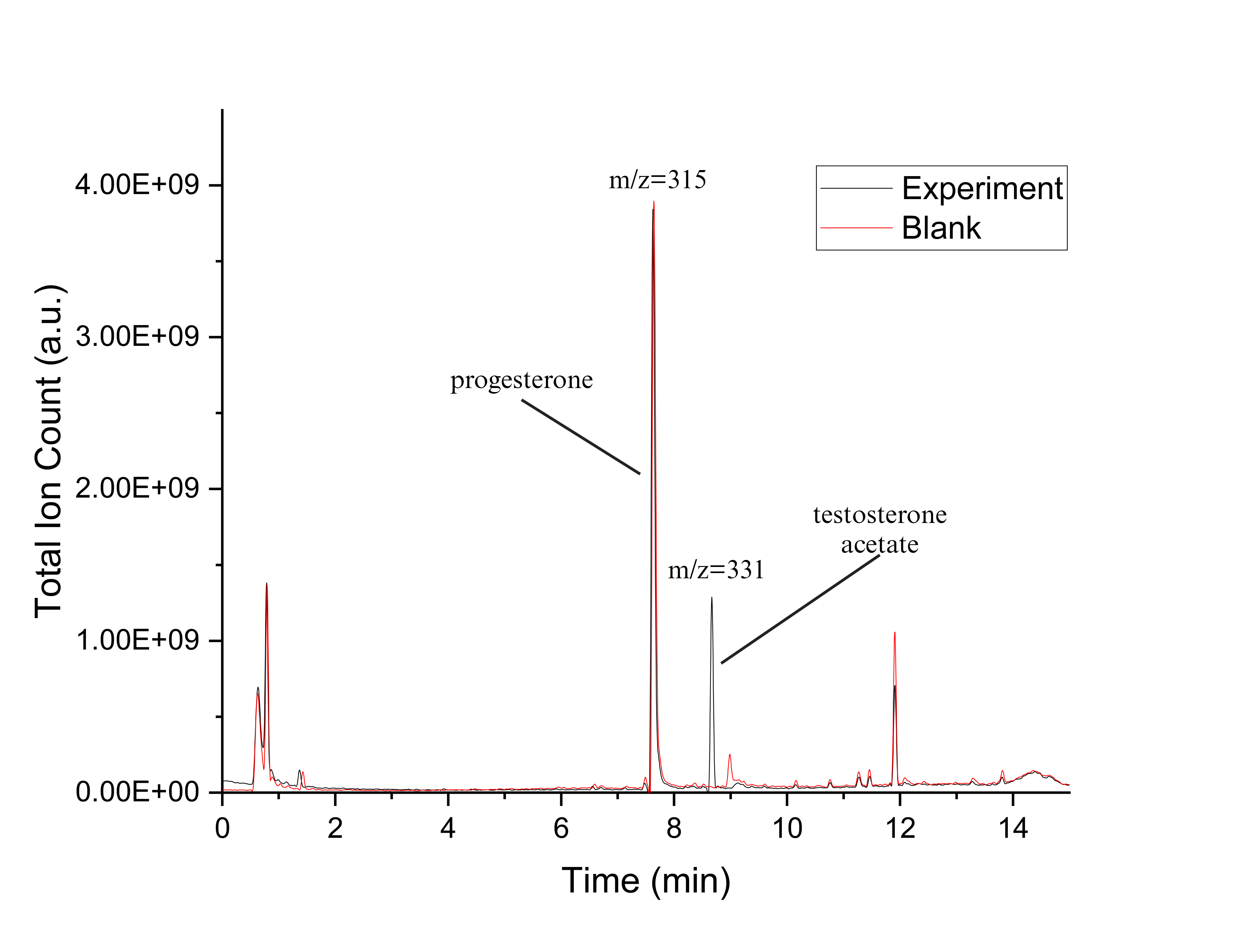

**Figure S61.** UPLC chromatogram for PTDH-L1-*ssn*BVMO reaction with progesterone. ESI positive ion mode was used.

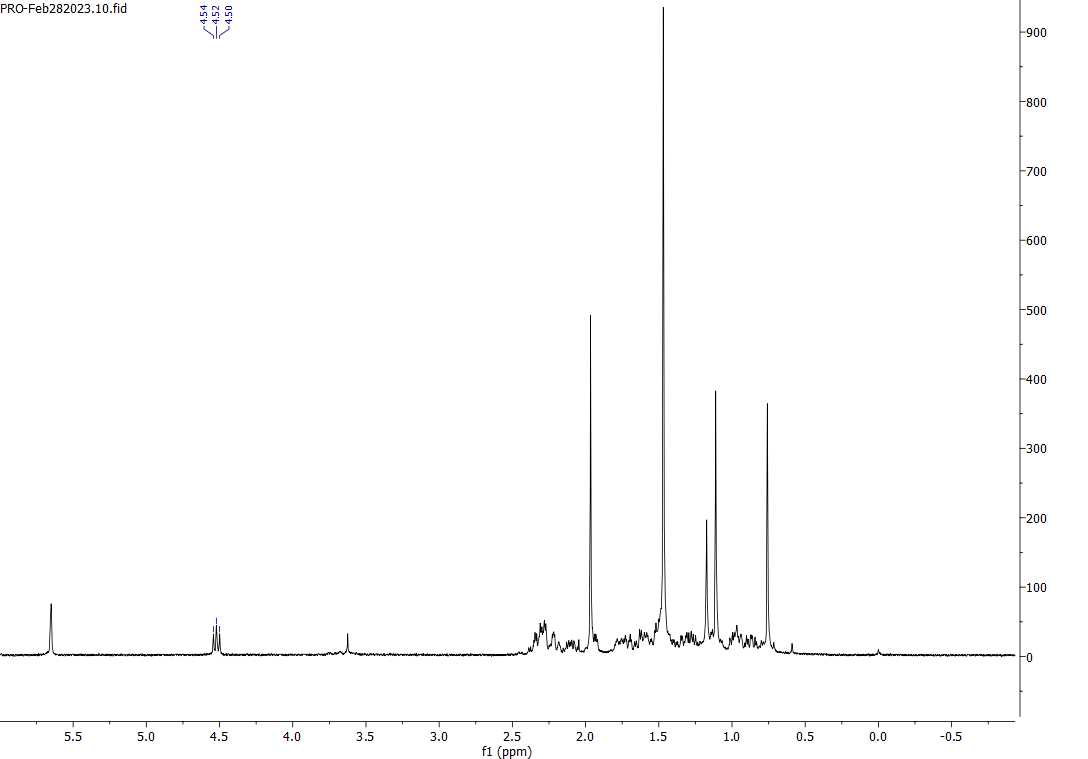

**Figure S62.** ^1^H NMR analysis for the product of *ssn*BVMO reaction with progesterone, showing the triplet de-shielding, indicative of oxygen insertion at the alkyl ketone of progesterone.

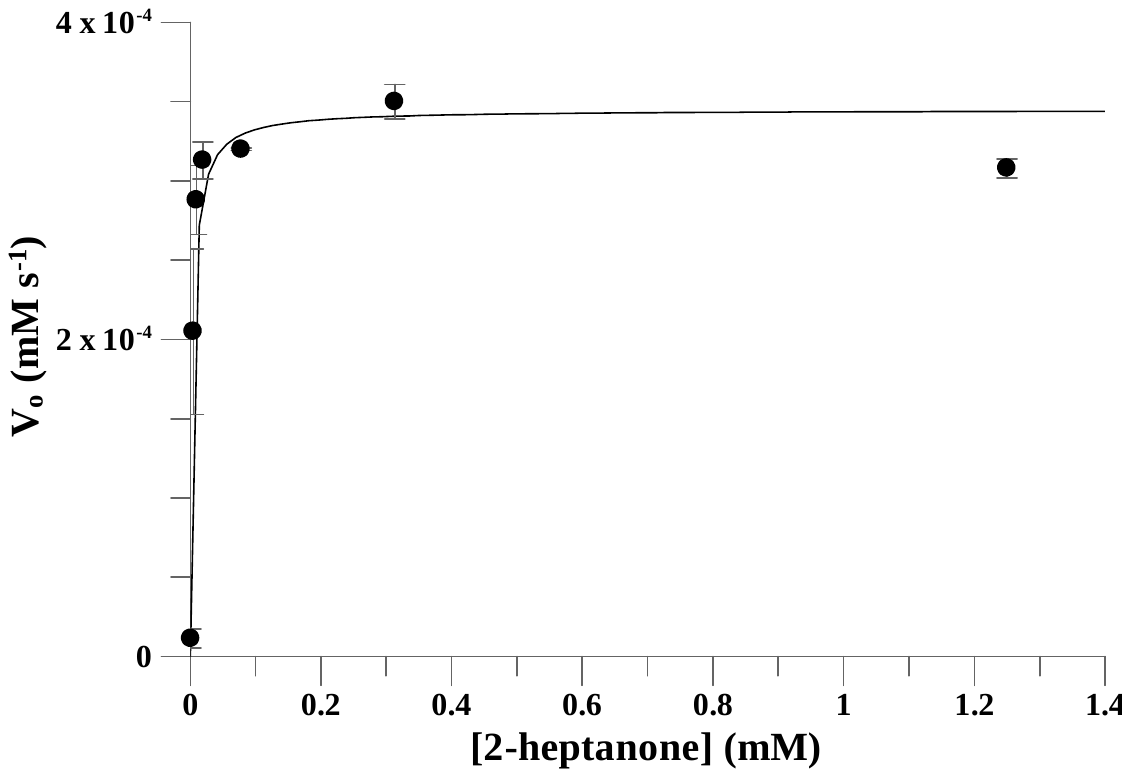

**Figure S63.** Fitting rate data for 2-heptanone to Michaelis-Menten equation used for determination of *V*_max_ (with corresponding *k*_cat_) and *K*_M_. Kinetic measurements were carried out using the following concentrations, [*ssn*BVMO] = 0.5 µM, and [NADPH] = 0.1 mM, and were carried out in triplicates.

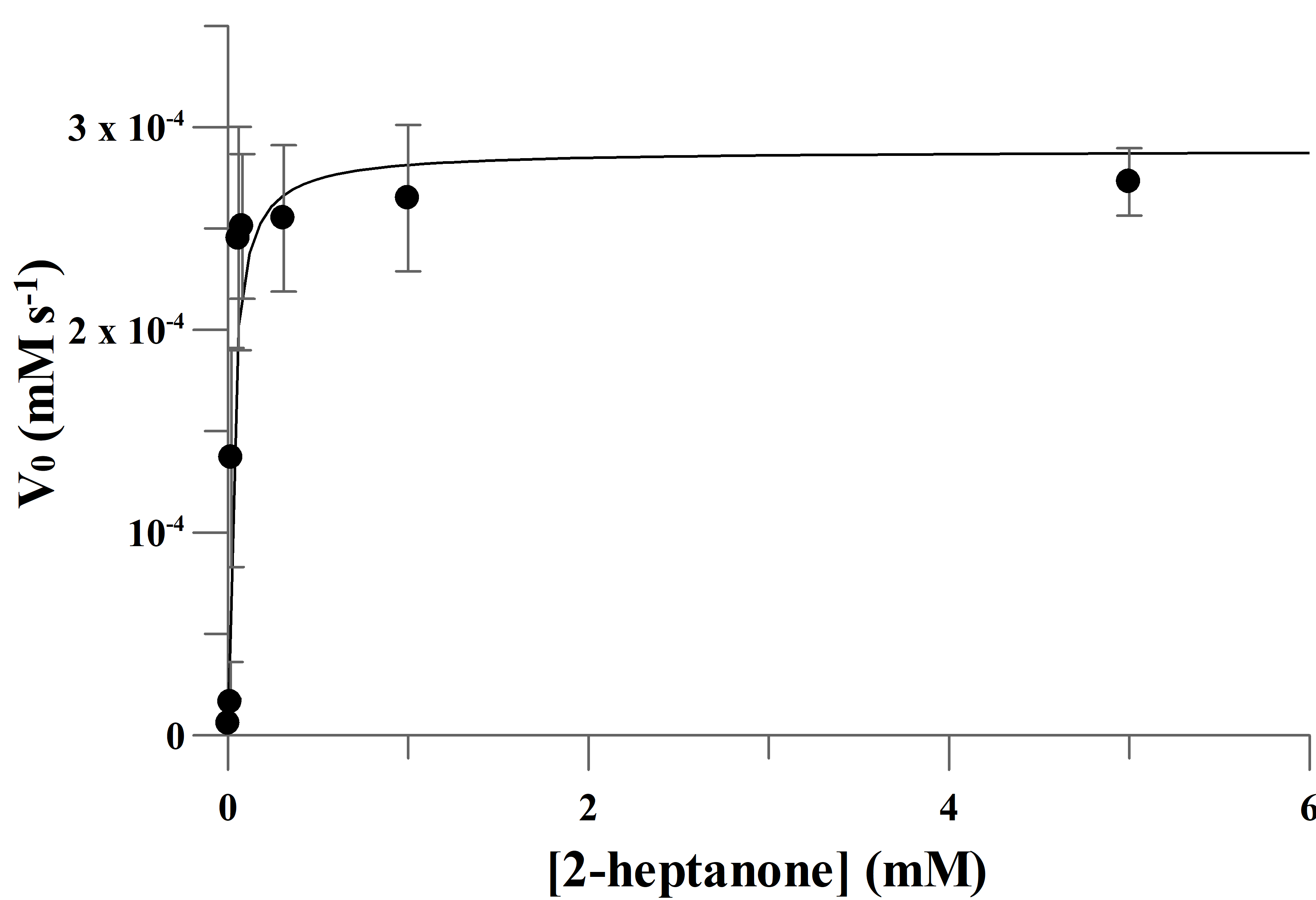

**Figure S64.** Fitting rate data for 2-heptanone to Michaelis-Menten equation used for determination of *V*_max_ (with corresponding *k*_cat_) and *K*_M_. Kinetic measurements were carried out using the following concentrations, [PTDH-L1-*ssn*BVMO] = 0.5 µM, and [NADPH] = 0.1 mM, and were carried out in triplicates.

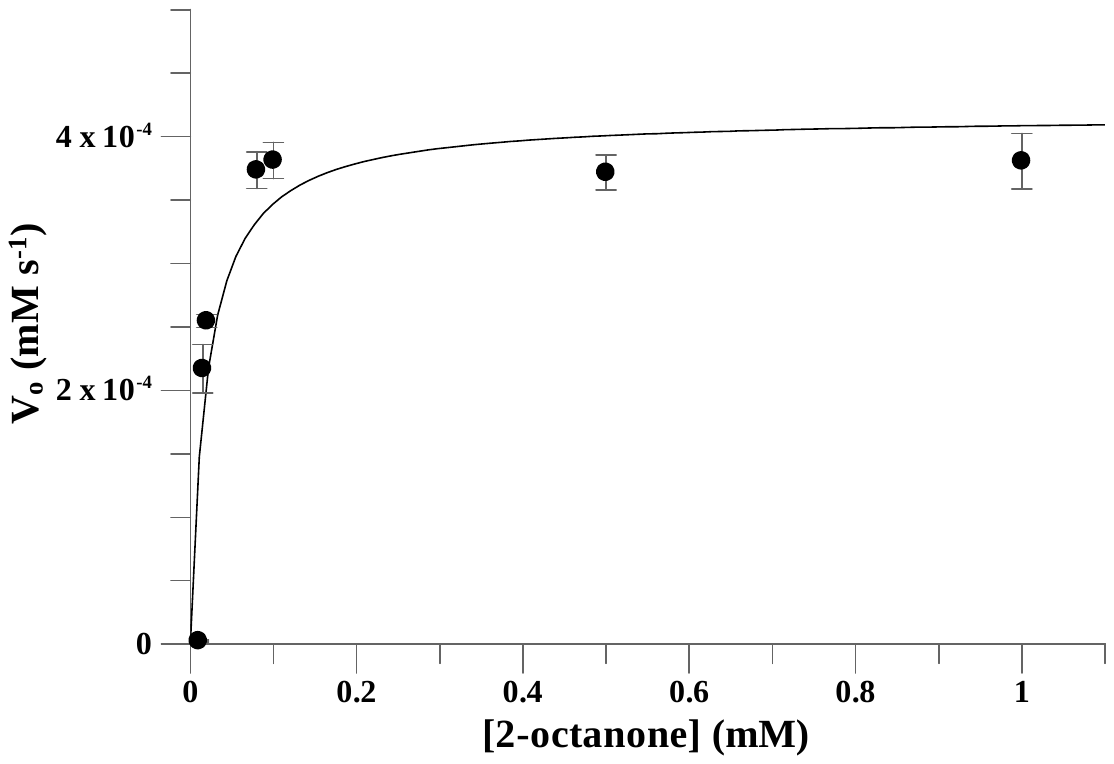

**Figure S65.** Fitting rate data for 2-octanone to Michaelis-Menten equation used for determination of *V*_max_ (with corresponding *k*_cat_) and *K*_M_. Kinetic measurements were carried out using the following concentrations, [*ssn*BVMO] = 0.5 µM, and [NADPH] = 0.1 mM, and were carried out in triplicates.

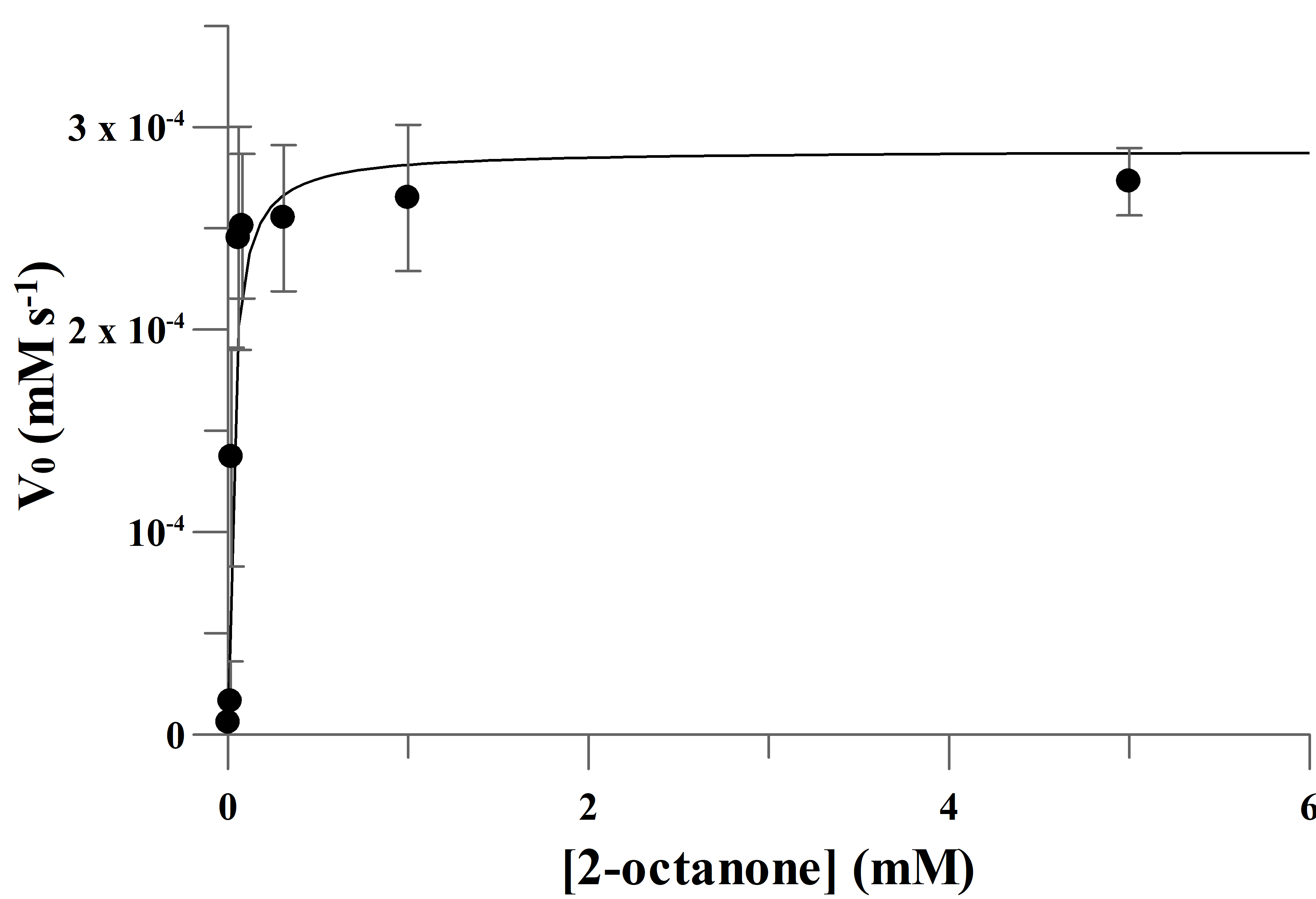

**Figure S66.** Fitting rate data for 2-octanone to Michaelis-Menten equation used for determination of *V*_max_ (with corresponding *k*_cat_) and *K*_M_. Kinetic measurements were carried out using the following concentrations, [PTDH-L1-*ssn*BVMO] = 0.5 µM, and [NADPH] = 0.1 mM, and were carried out in triplicates.

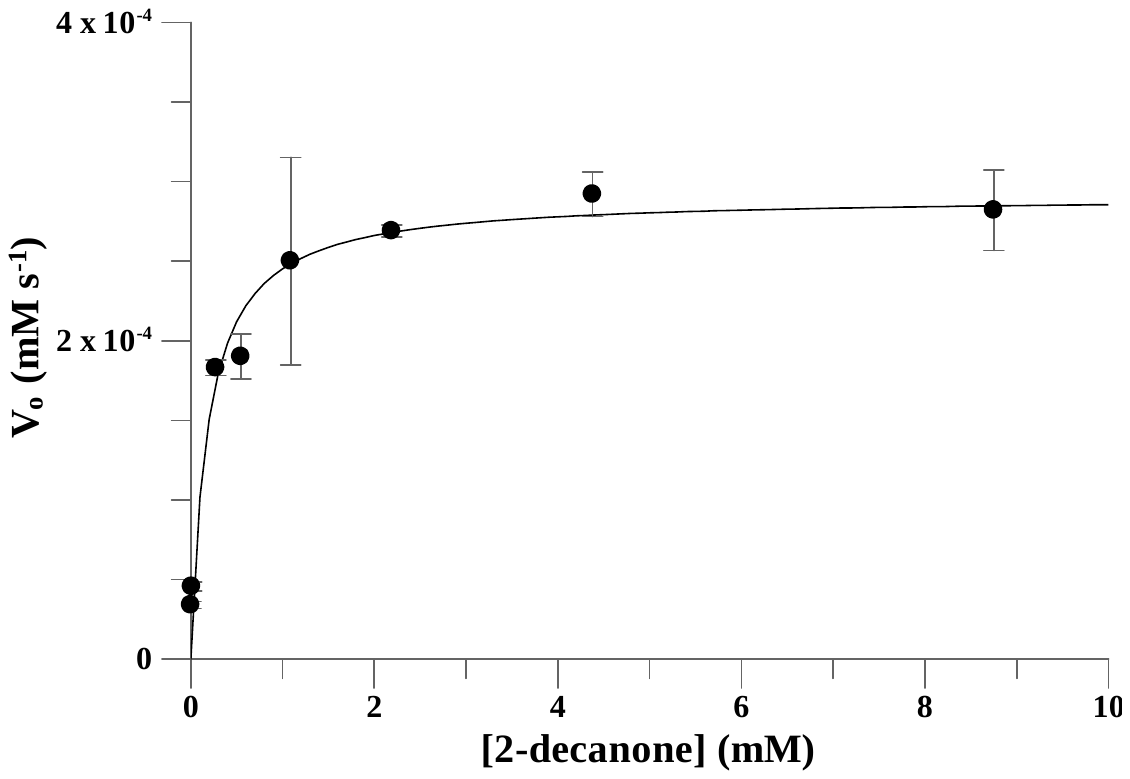

**Figure S67.** Fitting rate data for 2-decanone to Michaelis-Menten equation used for determination of *V*_max_ (with corresponding *k*_cat_) and *K*_M_. Kinetic measurements were carried out using the following concentrations, [*ssn*BVMO] = 0.5 µM, and [NADPH] = 0.1 mM, and were carried out in triplicates.

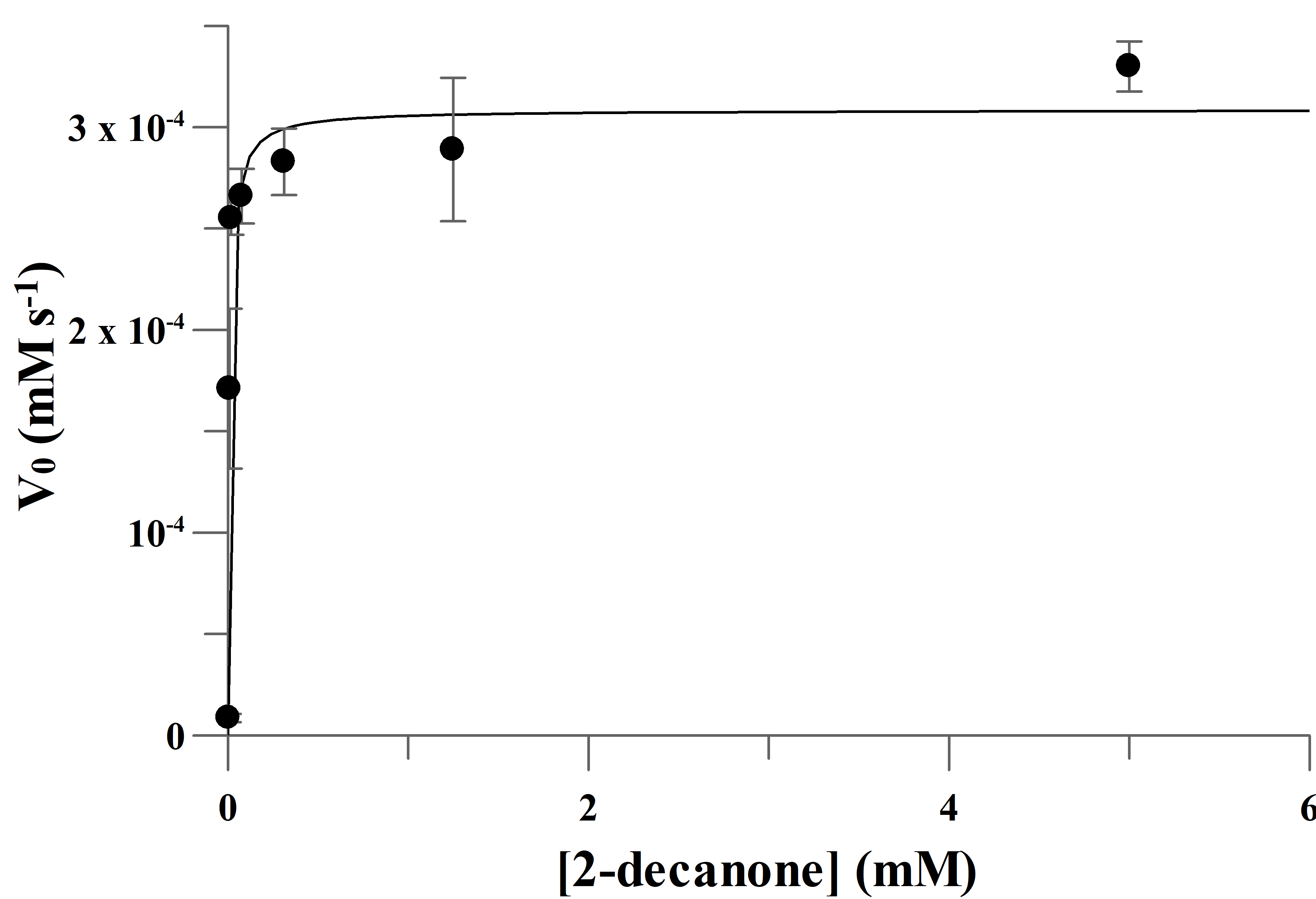

**Figure S68.** Fitting rate data for 2-decanone to Michaelis-Menten equation used for determination of *V*_max_ (with corresponding *k*_cat_) and *K*_M_. Kinetic measurements were carried out using the following concentrations, [PTDH-L1-*ssn*BVMO] = 0.5 µM, and [NADPH] = 0.1 mM, and were carried out in triplicates.

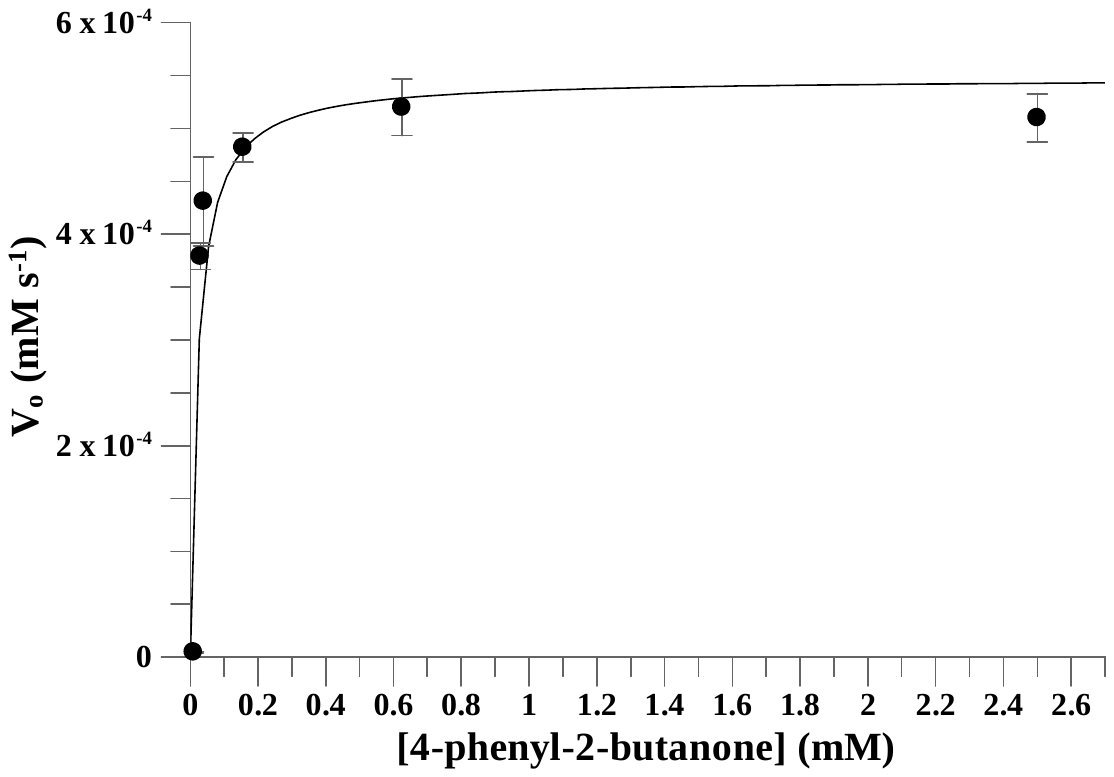

**Figure S69.** Fitting rate data for 4-phenyl-2-butanone to Michaelis-Menten equation used for determination of *V*_max_ (with corresponding *k*_cat_) and *K*_M_. Kinetic measurements were carried out using the following concentrations, [*ssn*BVMO] = 0.5 µM, and [NADPH] = 0.1 mM, and were carried out in triplicates.

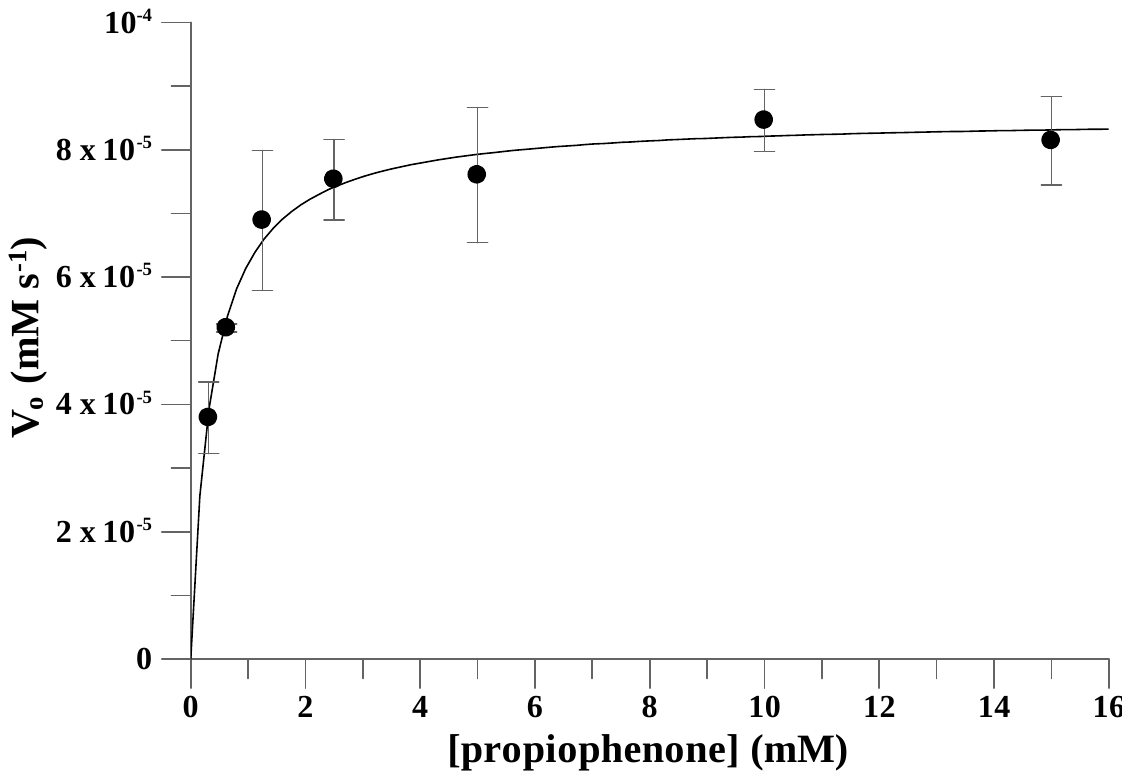

**Figure S70.** Fitting rate data for propiophenone to Michaelis-Menten equation used for determination of *V*_max_ (with corresponding *k*_cat_) and *K*_M_. Kinetic measurements were carried out using the following concentrations, [*ssn*BVMO] = 0.5 µM, and [NADPH] = 0.1 mM, and were carried out in triplicates.

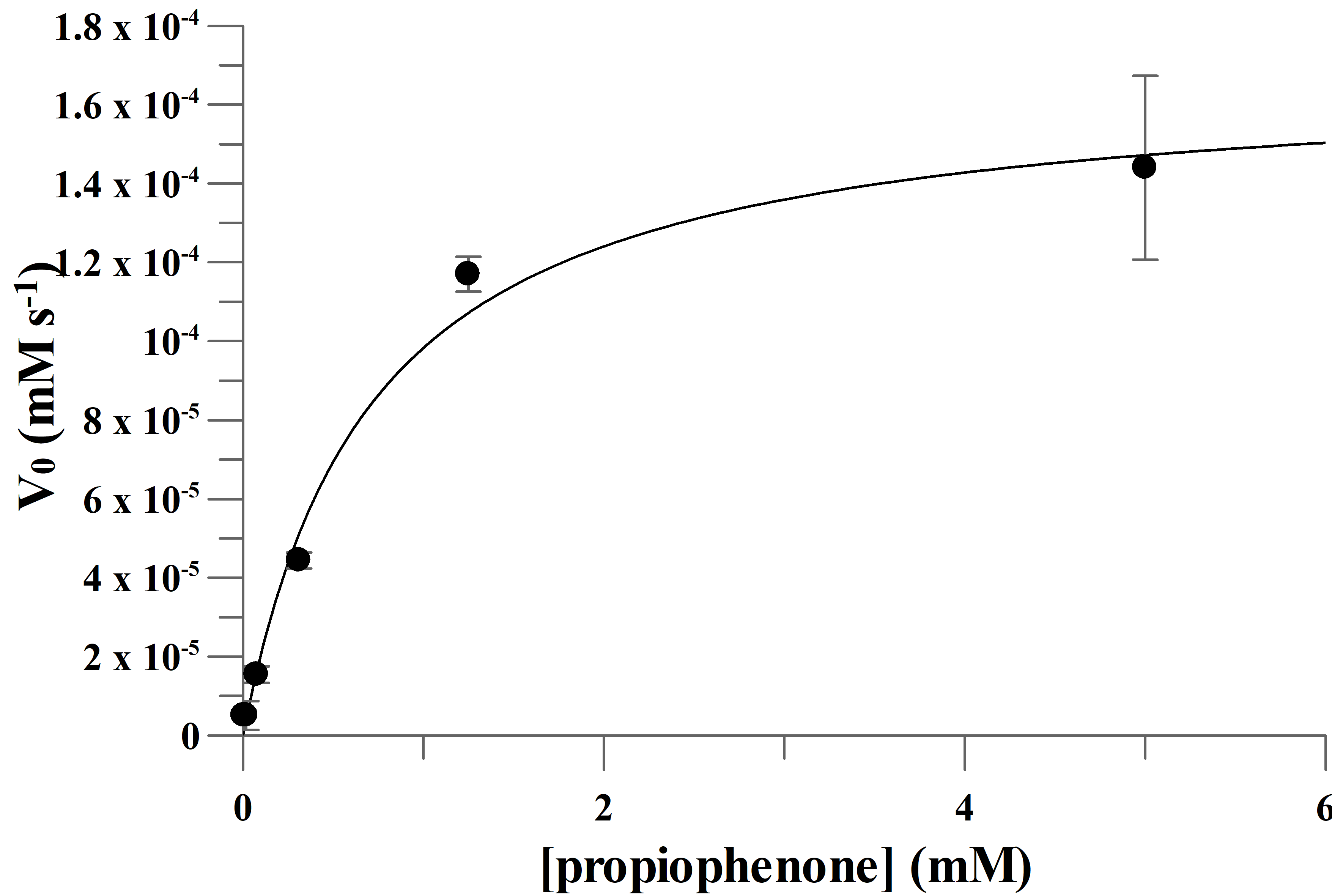

**Figure S71.** Fitting rate data for propiophenone to Michaelis-Menten equation used for determination of *V*_max_ (with corresponding *k*_cat_) and *K*_M_. Kinetic measurements were carried out using the following concentrations, [PTDH-L1-*ssn*BVMO] = 0.5 µM, and [NADPH] = 0.1 mM, and were carried out in triplicates.

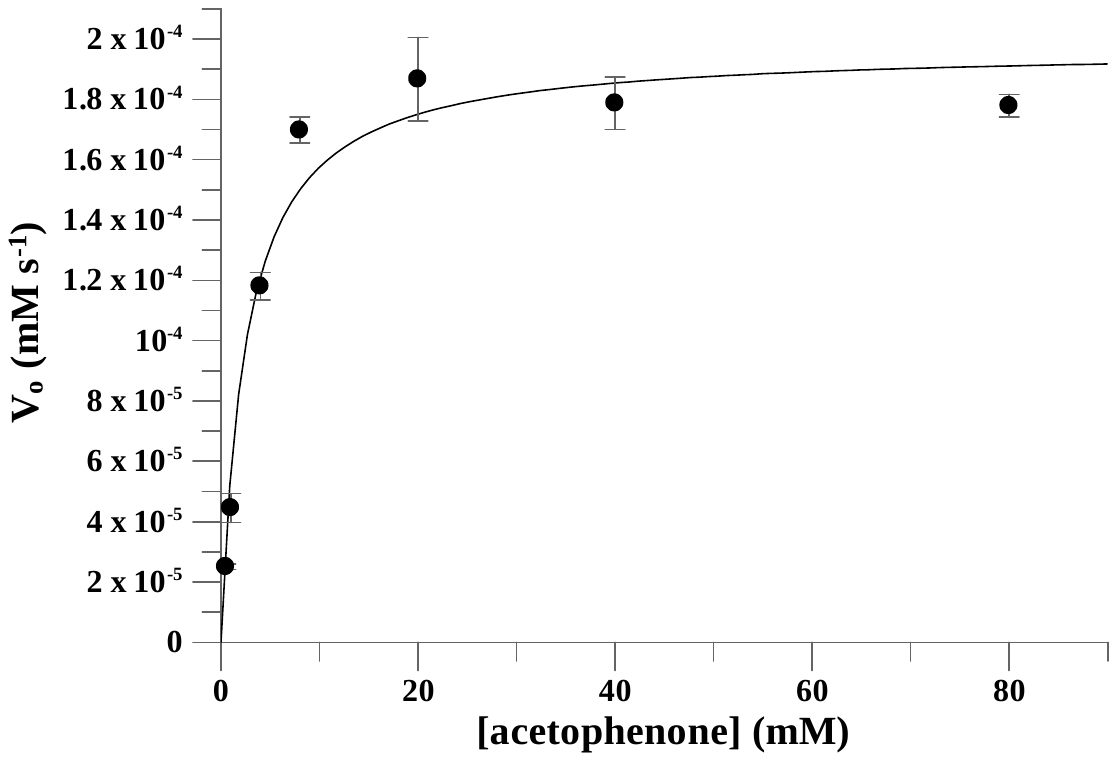

**Figure S72.** Fitting rate data for acetophenone to Michaelis-Menten equation used for determination of *V*_max_ (with corresponding *k*_cat_) and *K*_M_. Kinetic measurements were carried out using the following concentrations, [*ssn*BVMO] = 0.5 µM, and [NADPH] = 0.1 mM, and were carried out in triplicates.

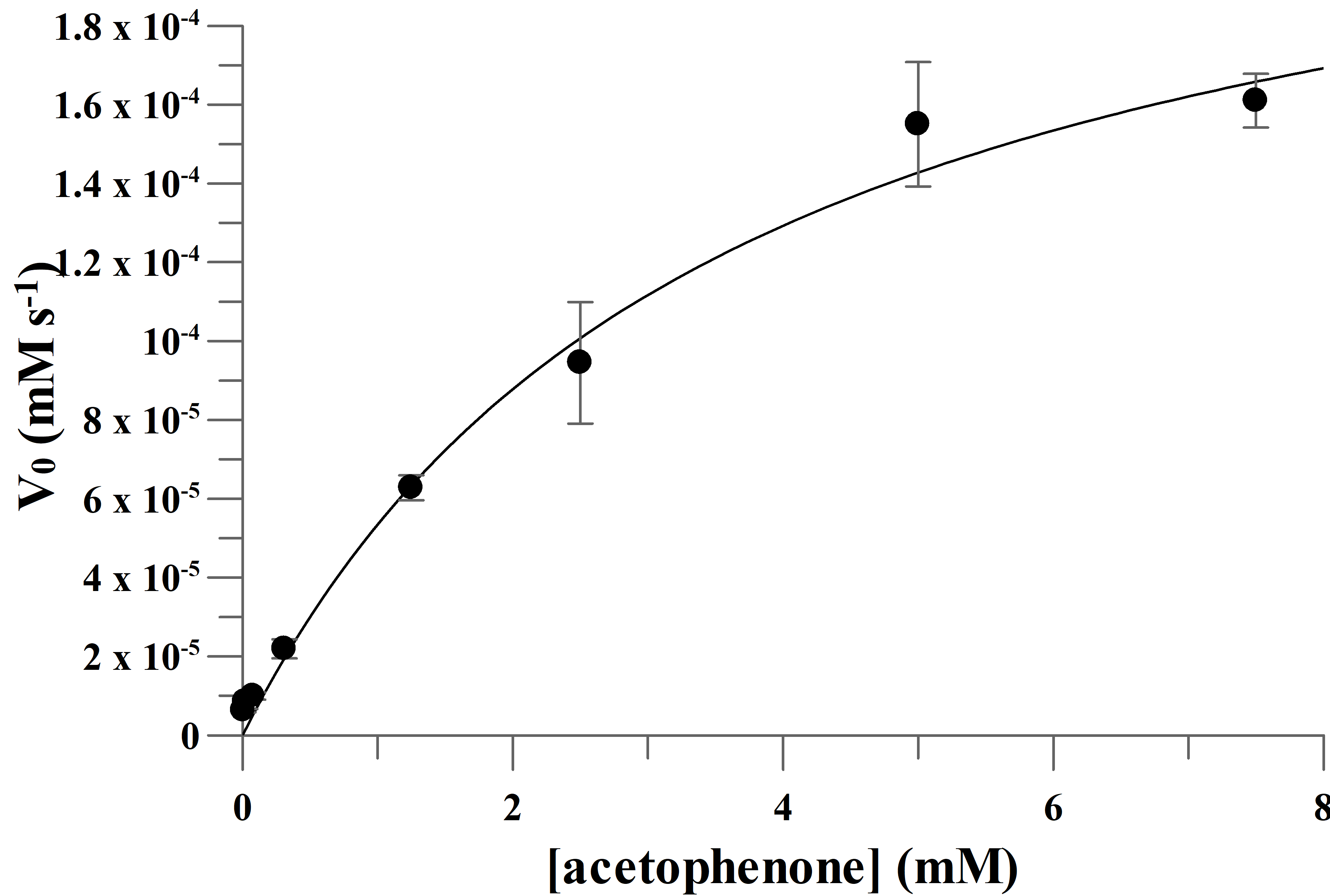

**Figure S73.** Fitting rate data for acetophenone to Michaelis-Menten equation used for determination of *V*_max_ (with corresponding *k*_cat_) and *K*_M_. Kinetic measurements were carried out using the following concentrations, [PTDH-L1-*ssn*BVMO] = 0.5 µM, and [NADPH] = 0.1 mM, and were carried out in triplicates.

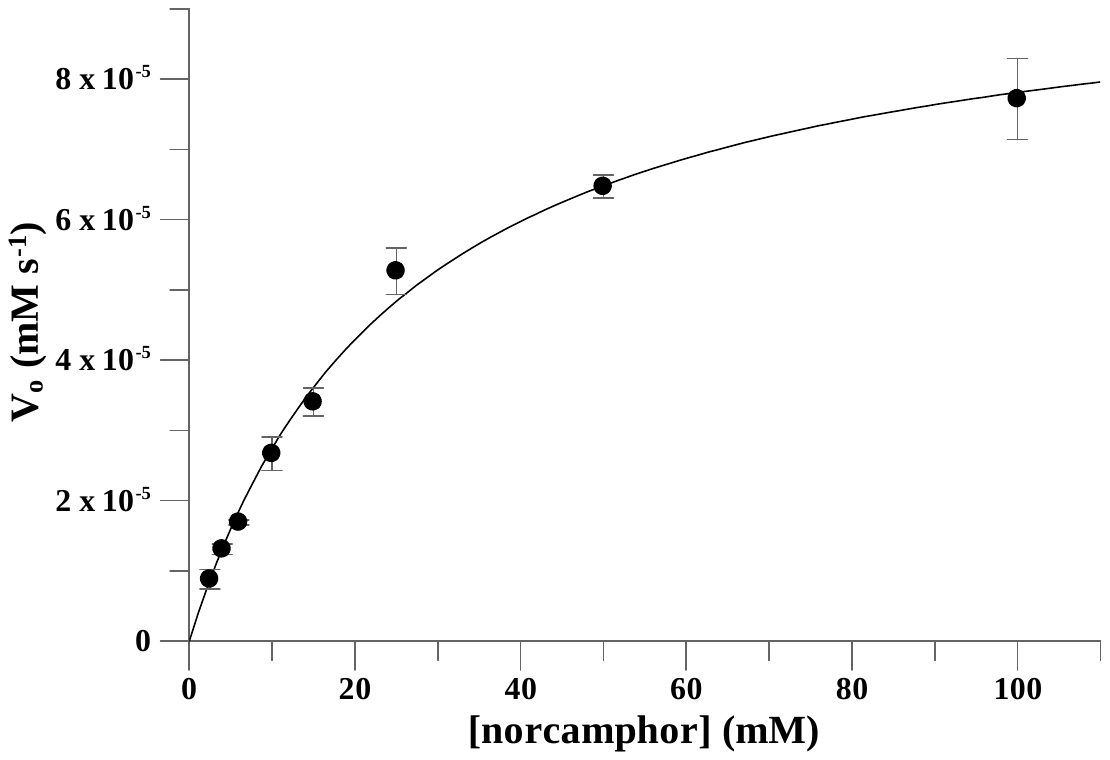

**Figure S74.** Fitting rate data for norcamphor to Michaelis-Menten equation used for determination of *V*_max_ (with corresponding *k*_cat_) and *K*_M_. Kinetic measurements were carried out using the following concentrations, [*ssn*BVMO] = 0.5 µM, and [NADPH] = 0.1 mM, and were carried out in triplicates.

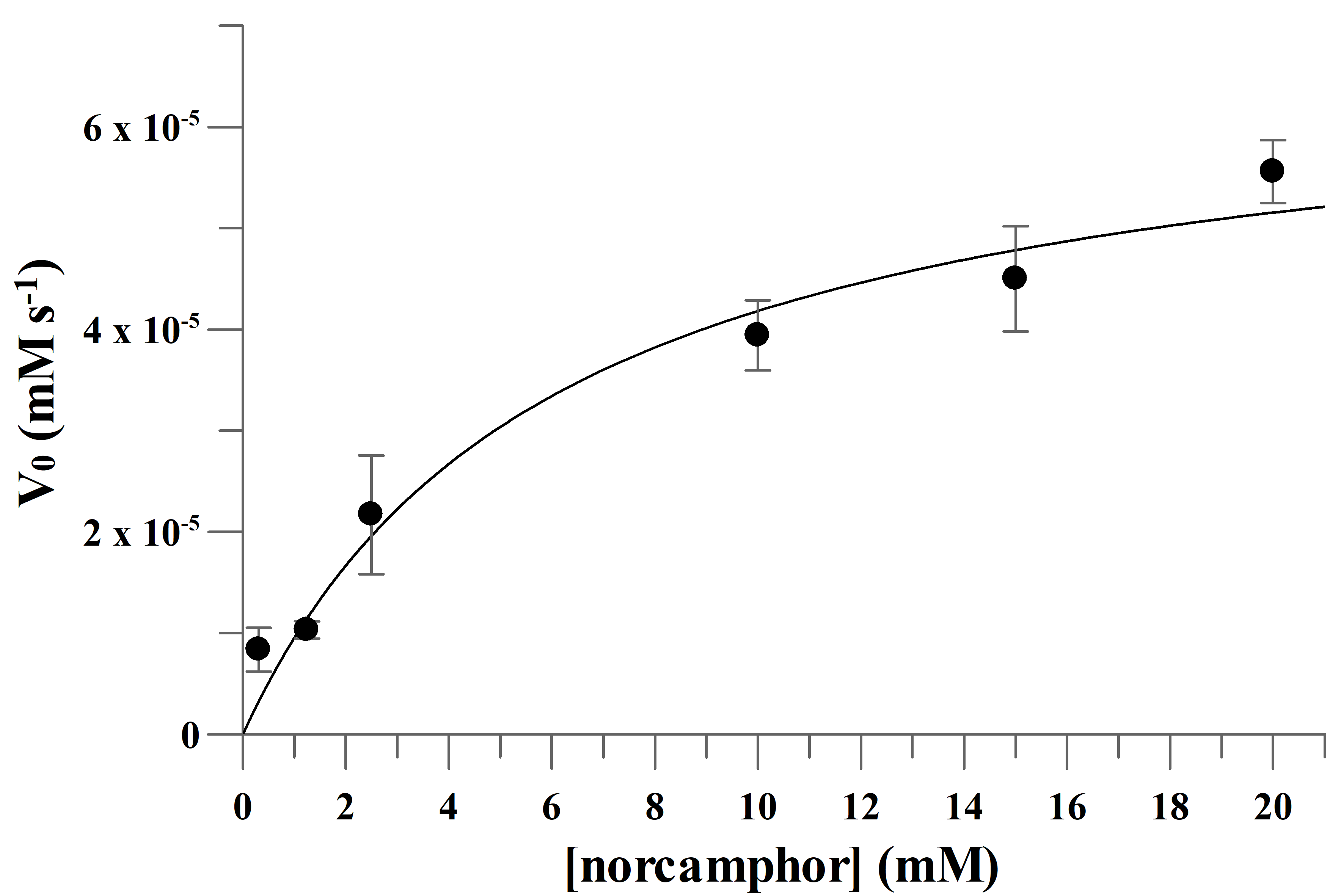

**Figure S75.** Fitting rate data for norcamphor to Michaelis-Menten equation used for determination of *V*_max_ (with corresponding *k*_cat_) and *K*_M_. Kinetic measurements were carried out using the following concentrations, [PTDH-L1-*ssn*BVMO] = 0.5 µM, and [NADPH] = 0.1 mM, and were carried out in triplicates.

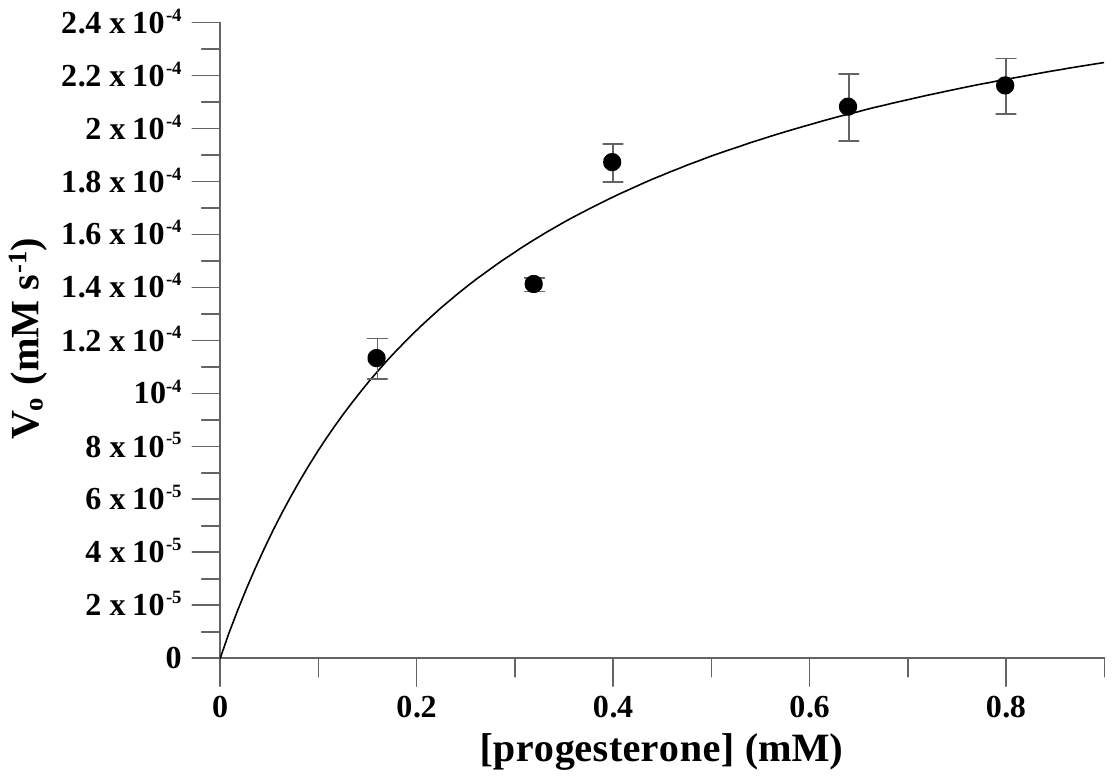

**Figure S76.** Fitting rate data for progesterone to Michaelis-Menten equation used for determination of *V*_max_ (with corresponding *k*_cat_) and *K*_M_. Kinetic measurements were carried out using the following concentrations, [*ssn*BVMO] = 0.5 µM, and [NADPH] = 0.1 mM, and were carried out in triplicates.

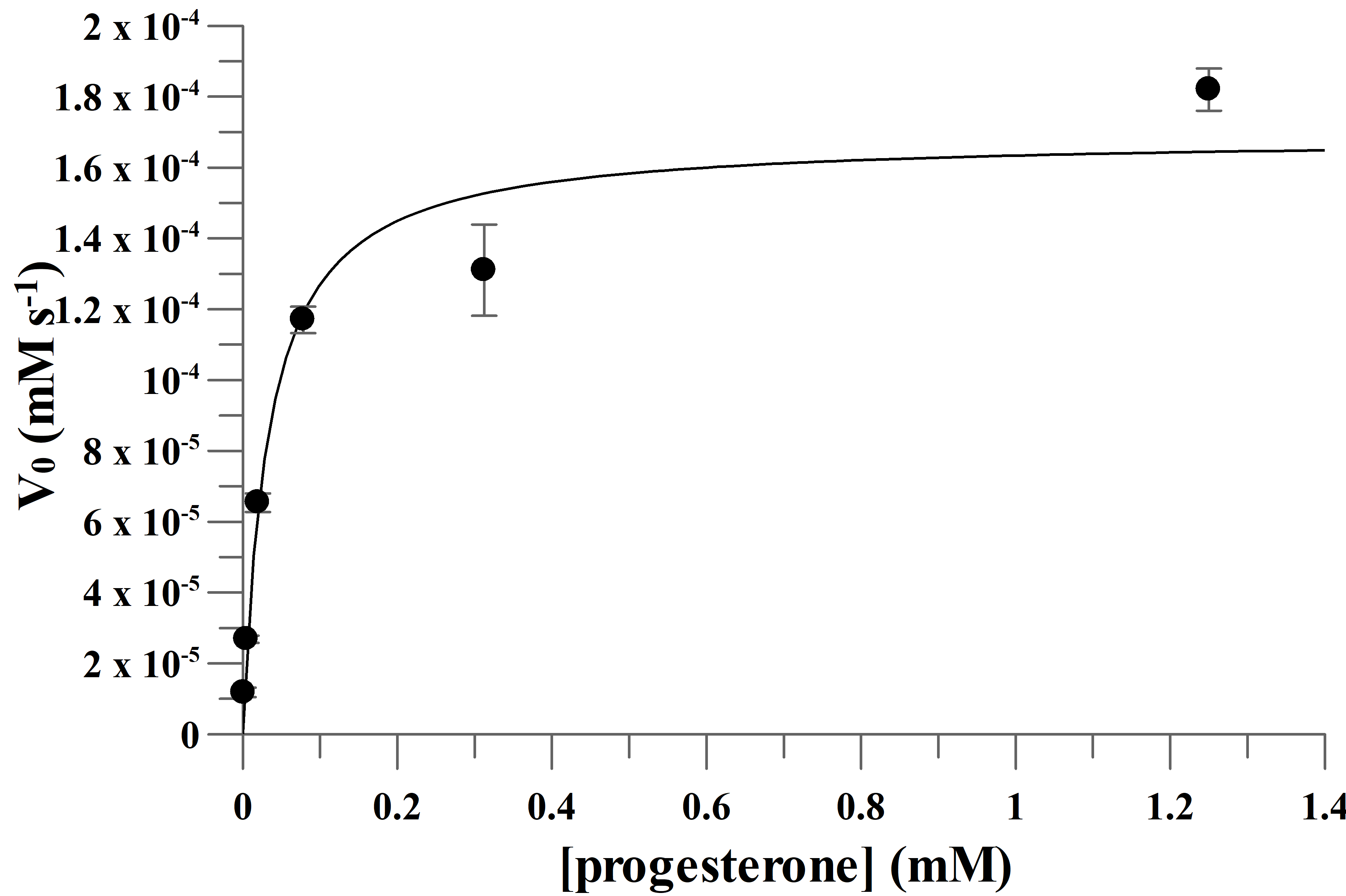

**Figure S77.** Fitting rate data for progesterone to Michaelis-Menten equation used for determination of *V*_max_ (with corresponding *k*_cat_) and *K*_M_. Kinetic measurements were carried out using the following concentrations, [PTDH-L1-*ssn*BVMO] = 0.5 µM, and [NADPH] = 0.1 mM, and were carried out in triplicates.

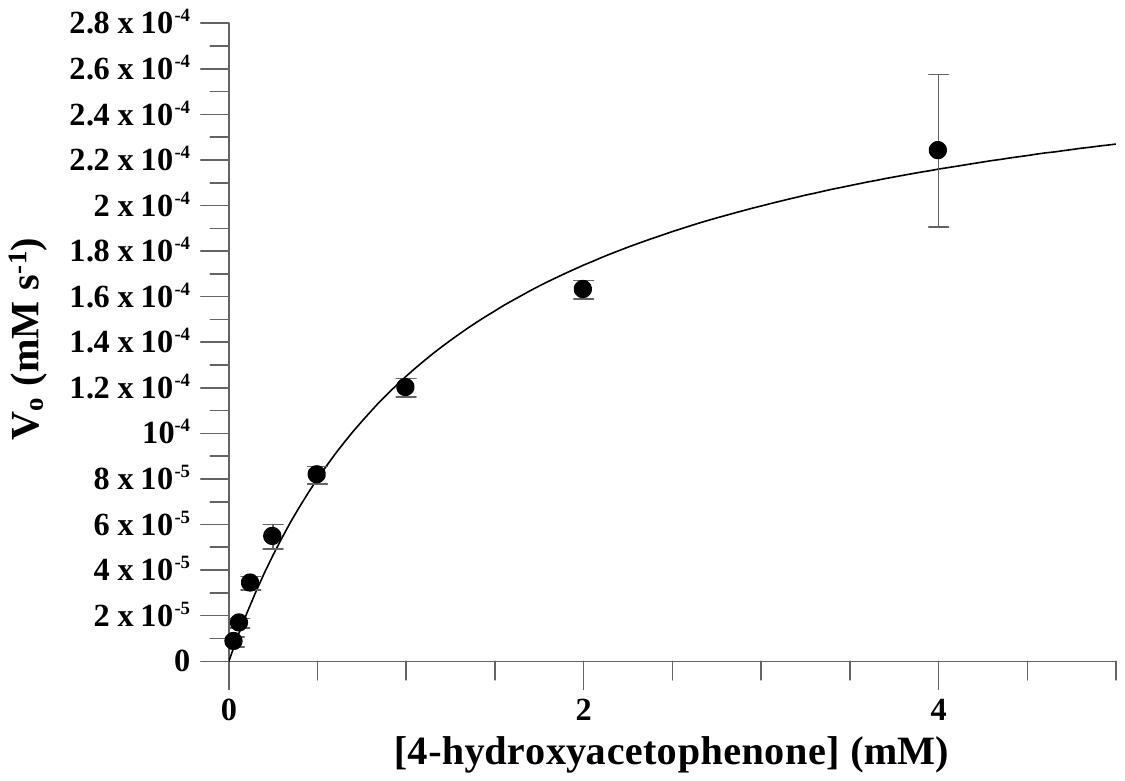

**Figure S78.** Fitting rate data for 4-hydroxyacetophenone to Michaelis-Menten equation used for determination of *V*_max_ (with corresponding *k*_cat_) and *K*_M_. Kinetic measurements were carried out using the following concentrations, [*ssn*BVMO] = 0.5 µM, and [NADPH] = 0.1 mM, and were carried out in triplicates.

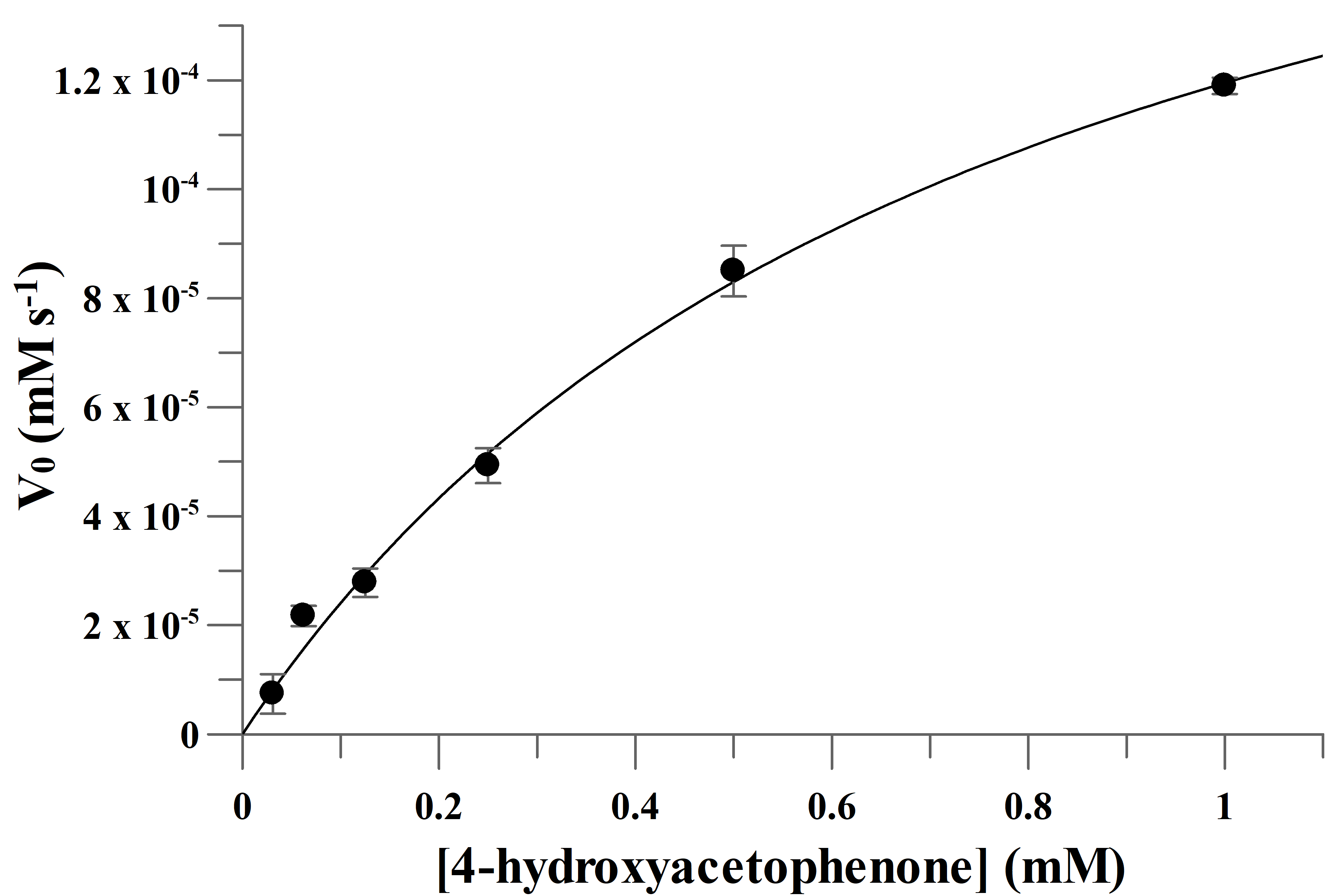

**Figure S79.** Fitting rate data for 4-hydroxyacetophenone to Michaelis-Menten equation used for determination of *V*_max_ (with corresponding *k*_cat_) and *K*_M_. Kinetic measurements were carried out using the following concentrations, [PTDH-L1-*ssn*BVMO] = 0.5 µM, and [NADPH] = 0.1 mM, and were carried out in triplicates.

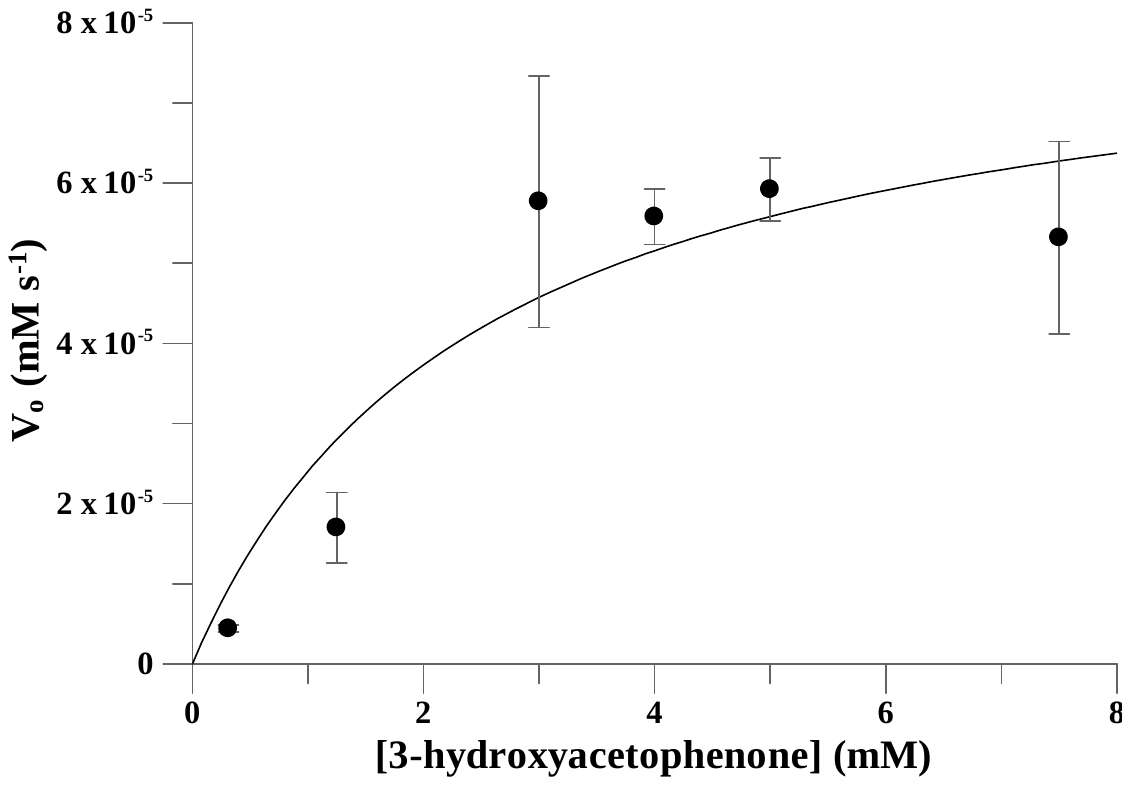

**Figure S80.** Fitting rate data for 3-hydroxyacetophenone to Michaelis-Menten equation used for determination of *V*_max_ (with corresponding *k*_cat_) and *K*_M_. Kinetic measurements were carried out using the following concentrations, [*ssn*BVMO] = 0.5 µM, and [NADPH] = 0.1 mM, and were carried out in triplicates.

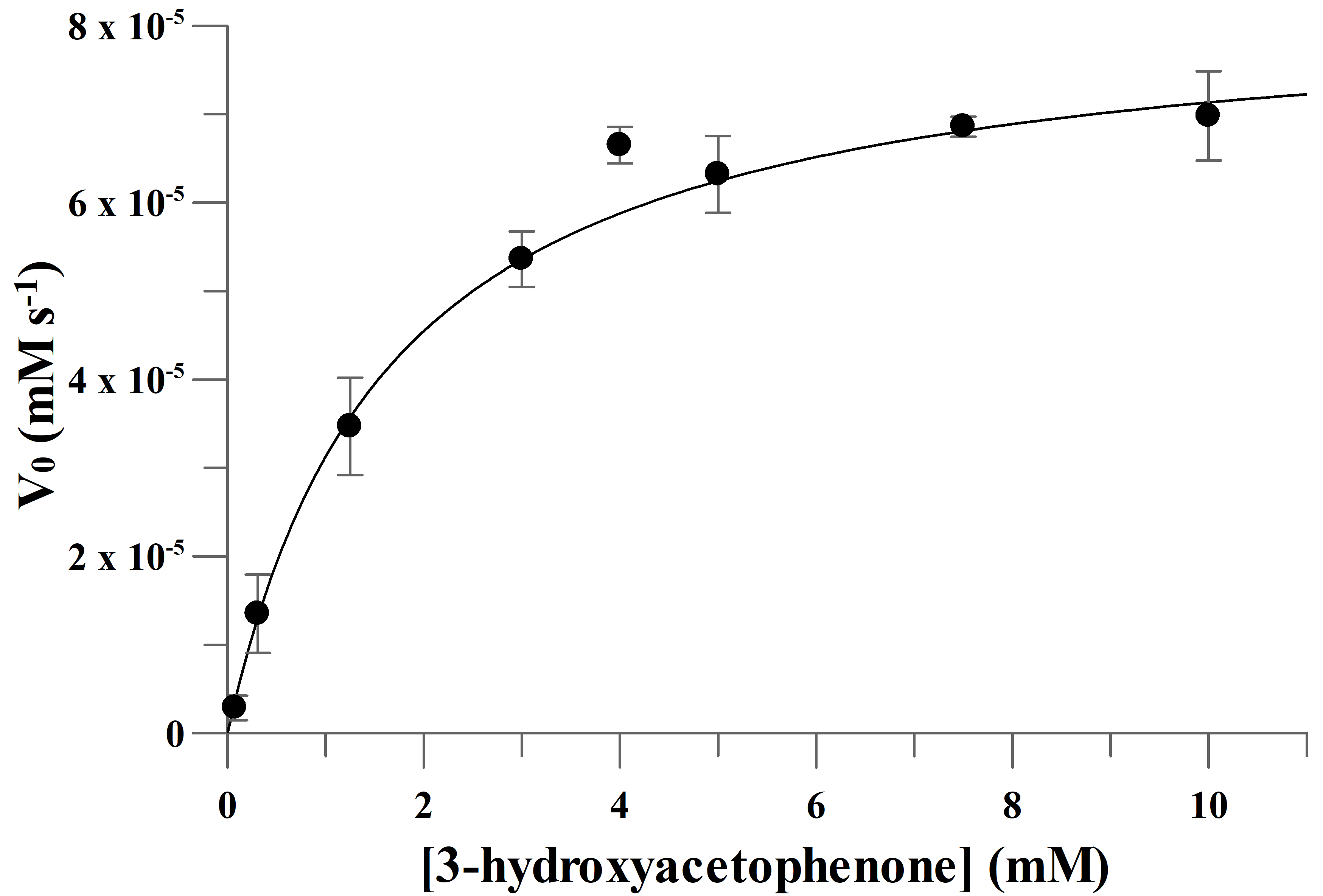

**Figure S81.** Fitting rate data for 3-hydroxyacetophenone to Michaelis-Menten equation used for determination of *V*_max_ (with corresponding *k*_cat_) and *K*_M_. Kinetic measurements were carried out using the following concentrations, [PTDH-L1-*ssn*BVMO] = 0.5 µM, and [NADPH] = 0.1 mM, and were carried out in triplicates.

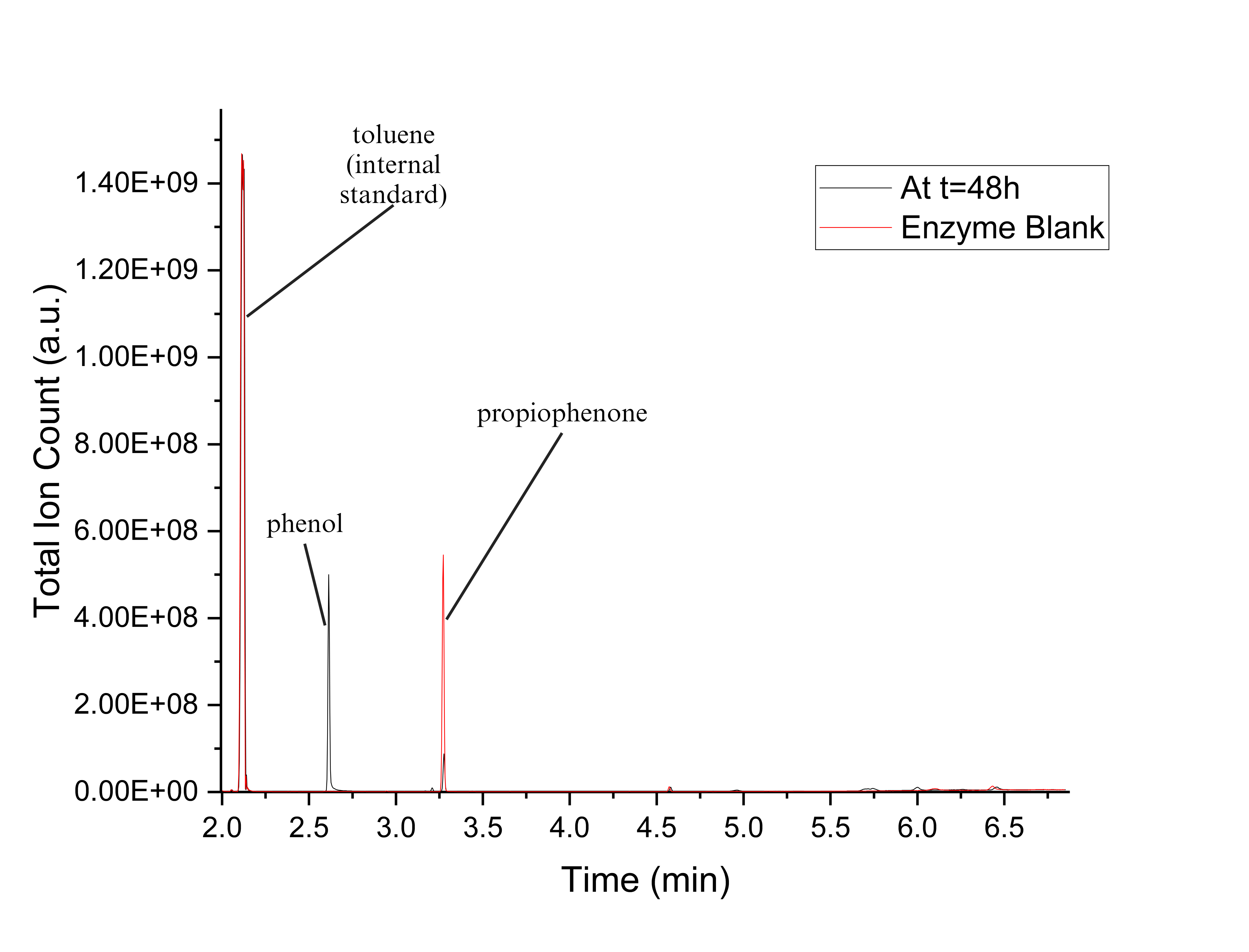

**Figure S82.** GCMS chromatogram for one of the replicates for propiophenone conversion experiments with *ssn*BVMO, and a corresponding enzyme blank overlaid.

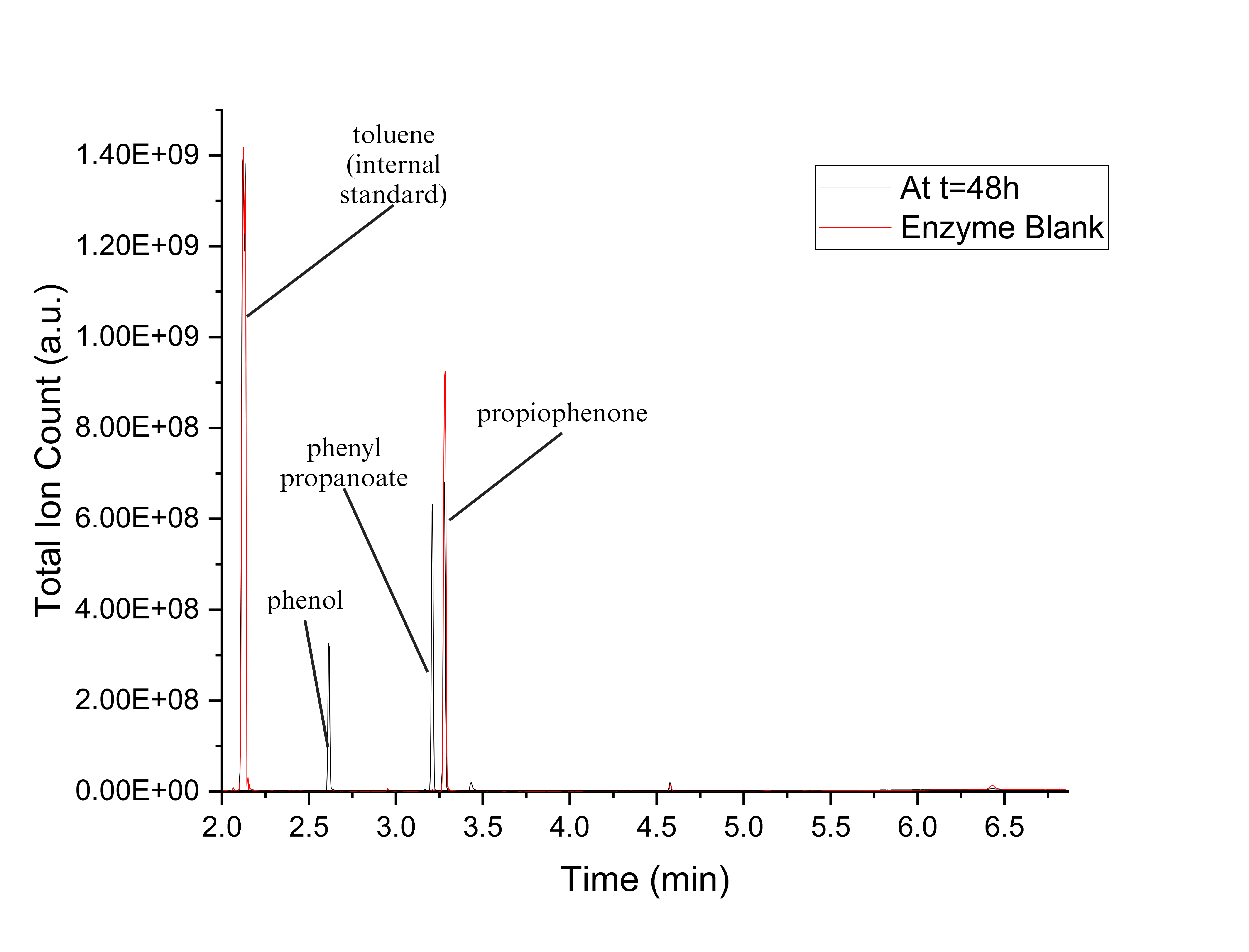

**Figure S83.** GCMS chromatogram for one of the replicates for propiophenone conversion experiments with PTDH-L1-*ssn*BVMO, and a corresponding enzyme blank overlaid.

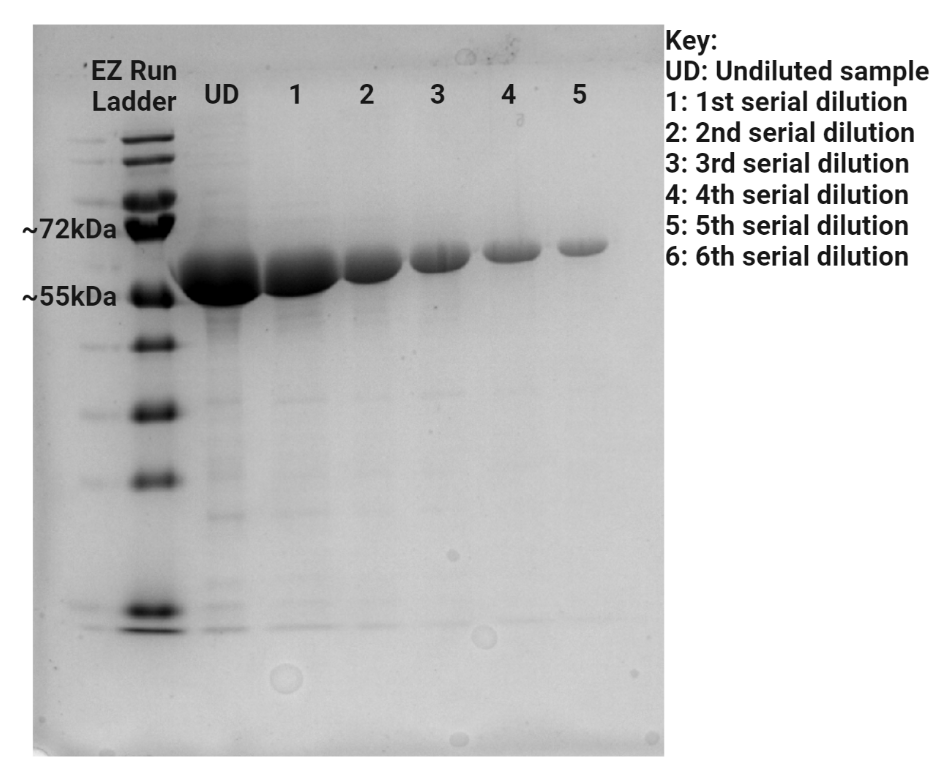

**Figure S84.** SDS-PAGE gel of purified *ssn*BVMO. The expected size of *ssn*BVMO is approximately 61 kDa.

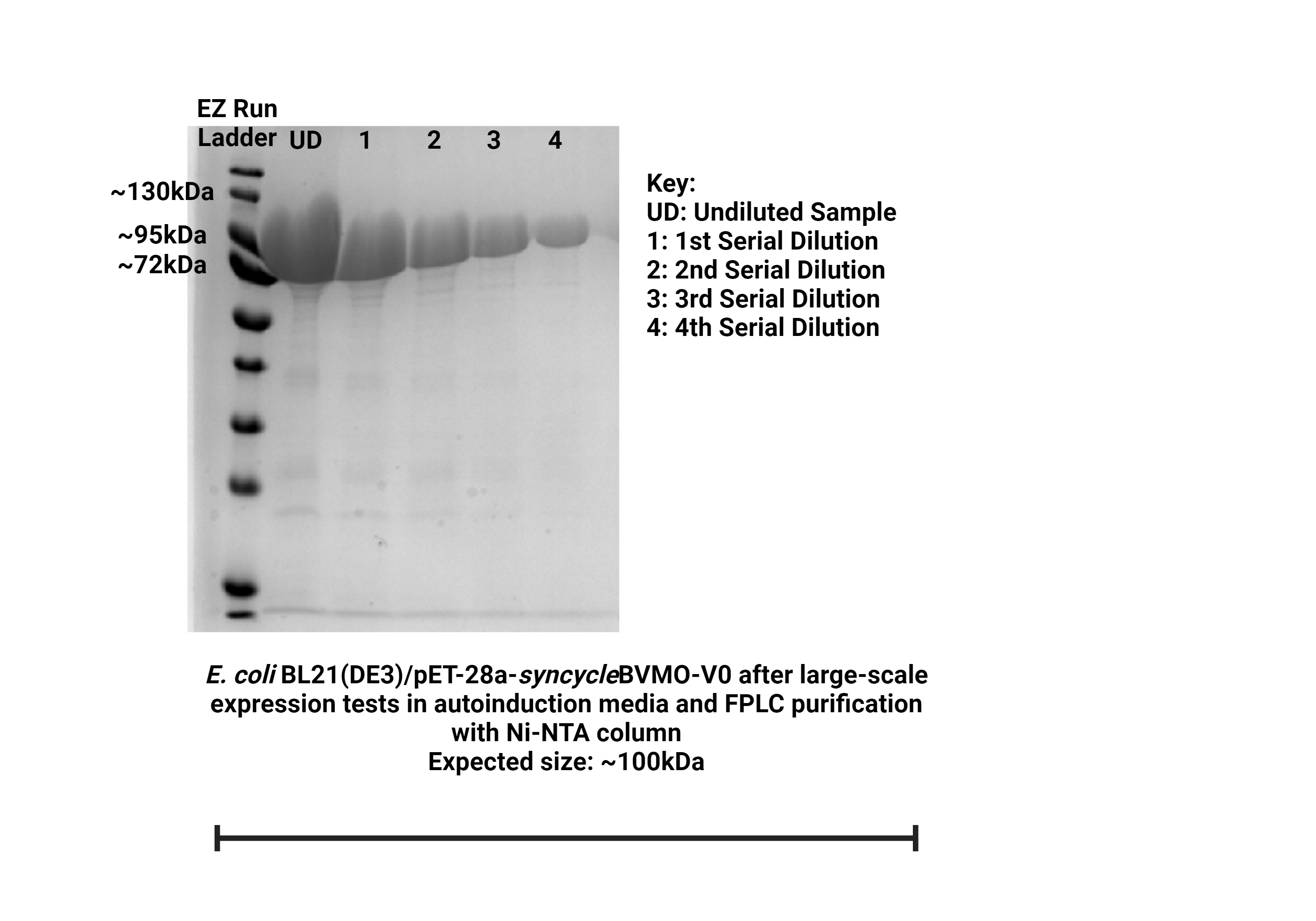

**Figure S85.** SDS-PAGE gel of purified PTDH-L1-*ssn*BVMO. The expected size of PTDH-L1-*ssn*BVMO is approximately 100 kDa.

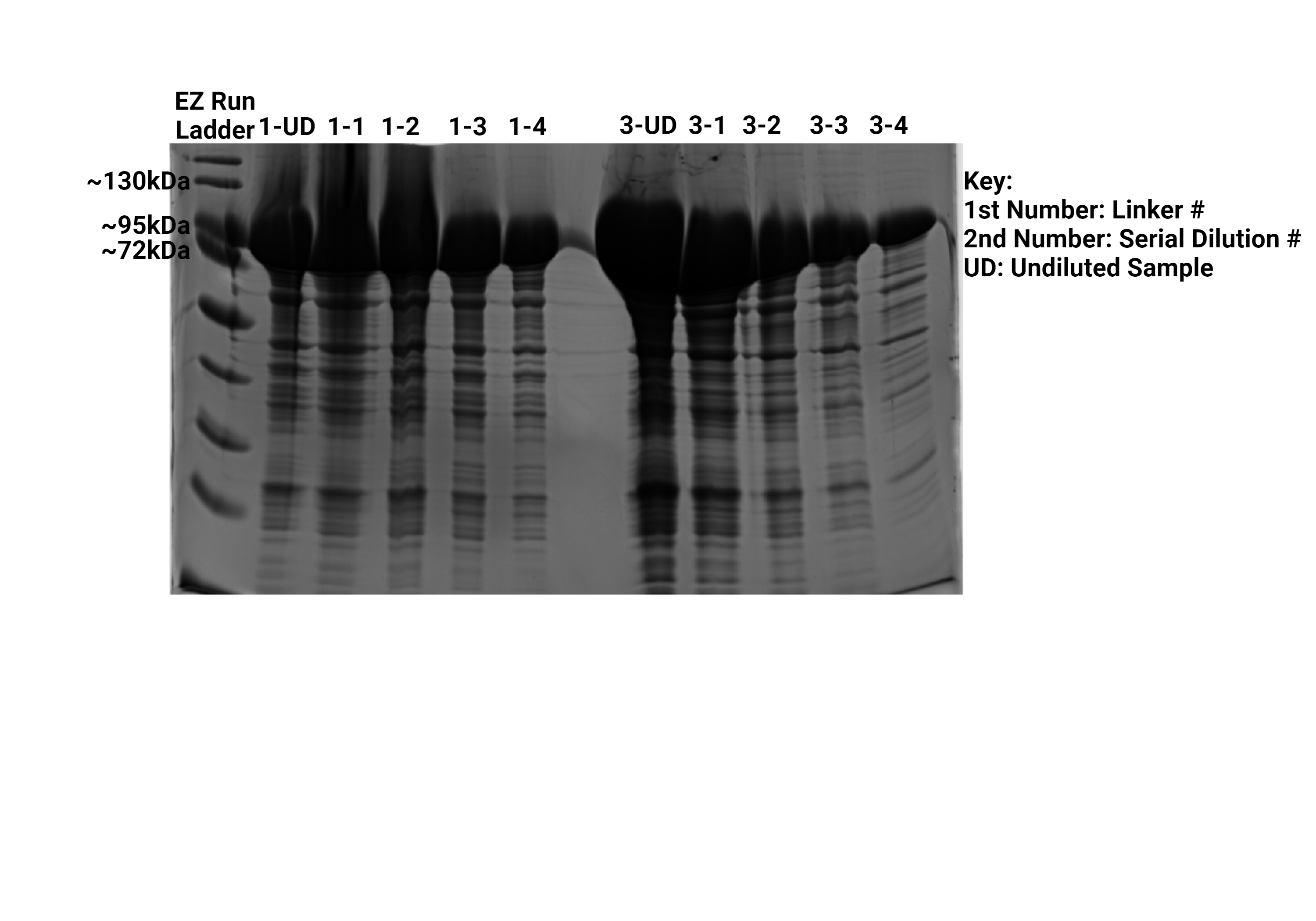

**Figure S86.** SDS-PAGE for preliminary purification of PTDH-L1-*ssn*BVMO (left 5 lanes), and PTDH-L3-*ssn*BVMO (right 5 lanes). The expected size of all PTDH-L-*ssn*BVMO variants is approximately 100 kDa.

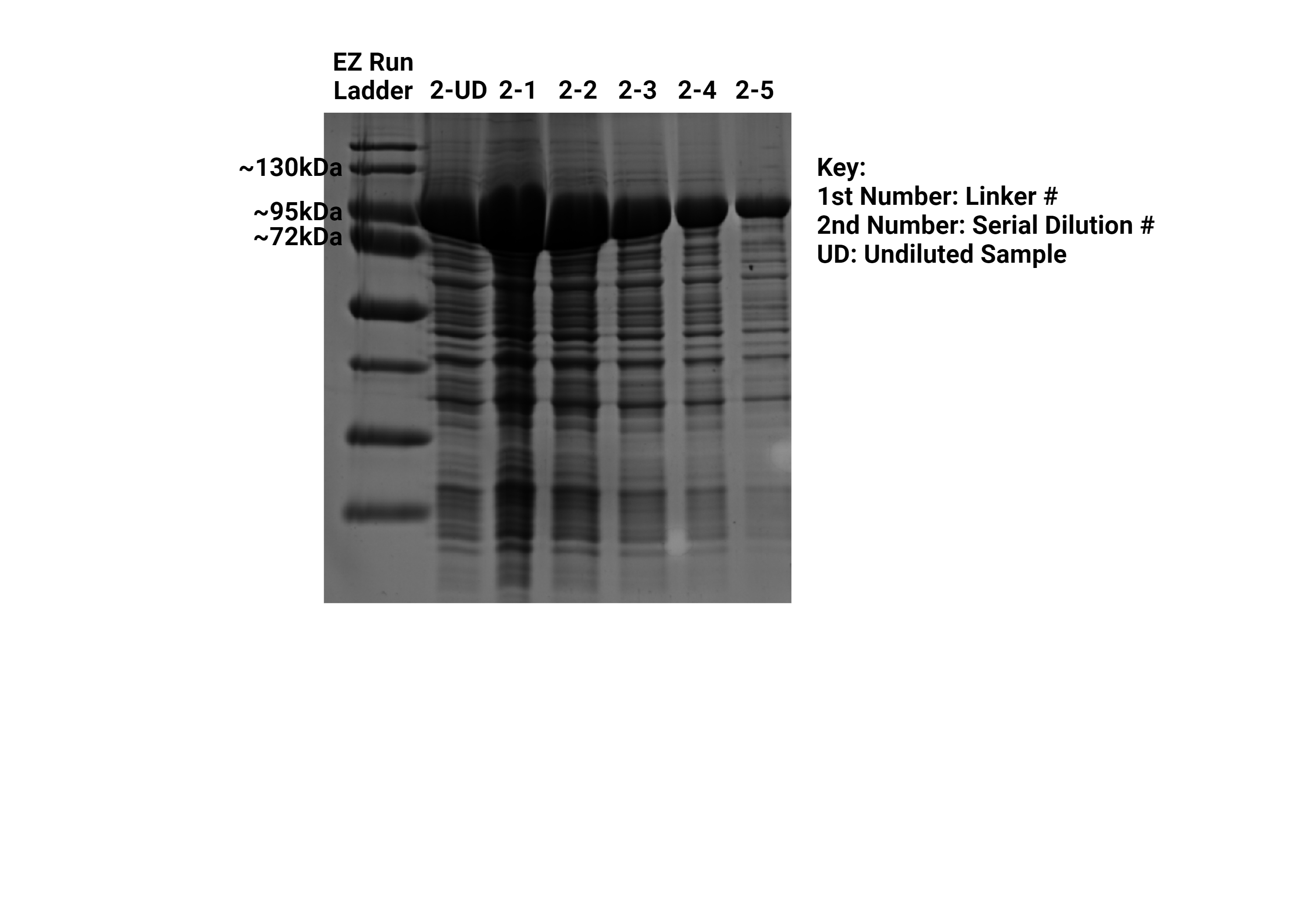

**Figure S87.** SDS-PAGE for preliminary purification of PTDH-L2-*ssn*BVMO. The expected size of all PTDH-L-*ssn*BVMO variants is approximately 100 kDa.

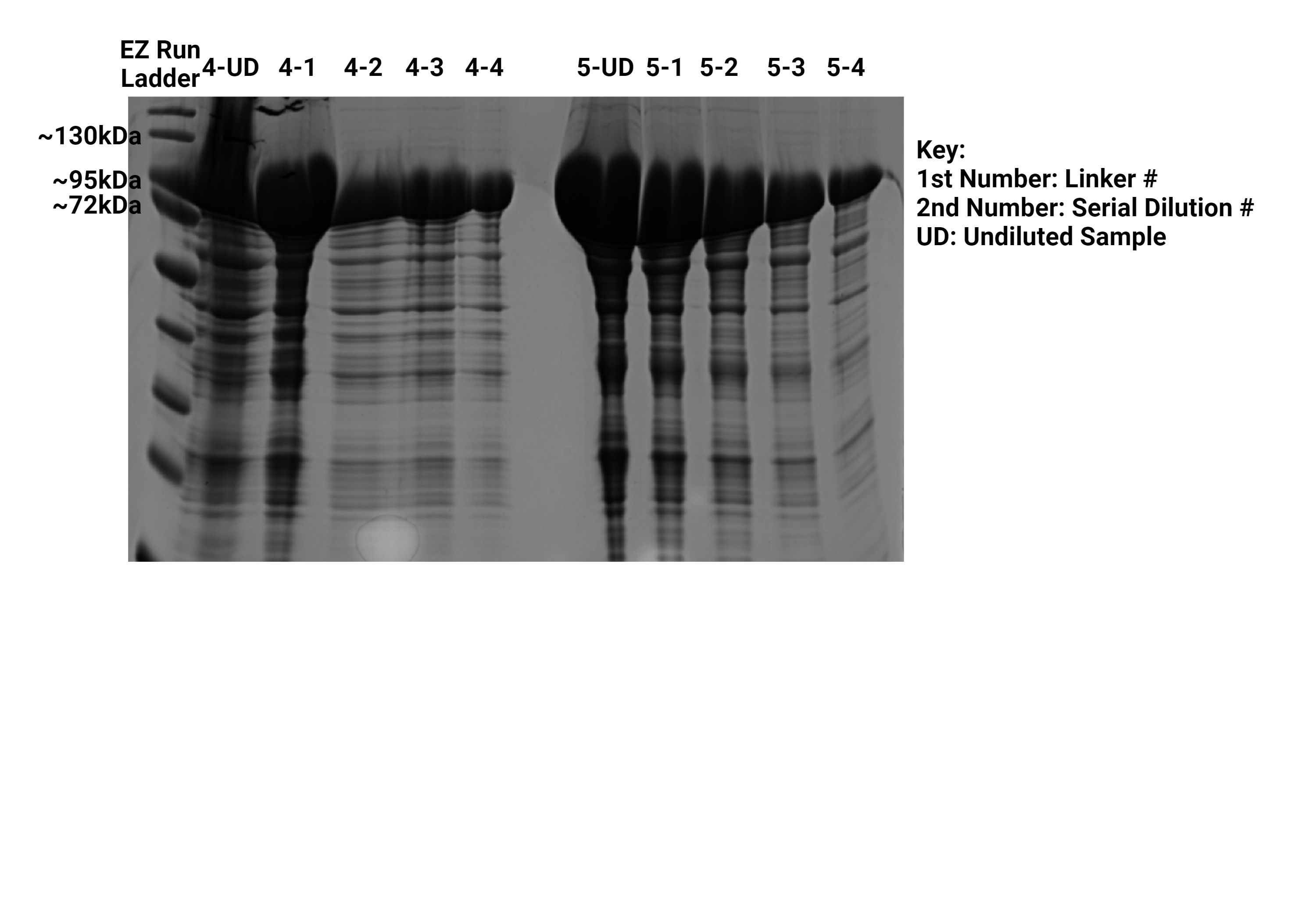

**Figure S88.** SDS-PAGE for preliminary purification of PTDH-L4-*ssn*BVMO (left 5 lanes), and PTDH-L5-*ssn*BVMO (right 5 lanes). The expected size of all PTDH-L-*ssn*BVMO variants is approximately 100 kDa.

**Table S5.** GC-MS settings used during analysis of *ssn*BVMO and PTDH-L1-*ssn*BVMO-catalyzed reactions.

| Parameter | Setting |
| --- | --- |
| Injection temperature | 250 °C |
| Split flow | 6.0 mL/min |
| Split ratio | 3 |
| Temperature program | Hold at 60 °C for 0.10 min, ramp to 200 °C at a rate of 65 °C/min and hold for 1.00 min. Then, ramp to 300 °C at a rate of 40 °C/min and hold for 1.10 min |

**Table S6.** UPLC-DAD-MS settings used during analysis of *ssn*BVMO- and PTDH-L1-*ssn*BVMO -catalyzed progesterone oxidations. Solvent A is deionized water, solvent B is HPLC-grade acetonitrile, and solvent C is 1% formic acid in water. Total run time is 15 minutes.

| Time (min) | Type | Flow (mL/min) | %A | %B | %C |
| --- | --- | --- | --- | --- | --- |
| 0-2 | Hold | 0.350 | 55 | 35 | 10 |
| 2-12 | Gradient | 0.350 | 55-0 | 35-90 | 10 |
| 12-13 | Hold | 0.350 | 0 | 90 | 10 |
| 13-14 | Gradient | 0.350 | 0-55 | 90-35 | 10 |
| 14-15 | Hold | 0.350 | 55 | 35 | 10 |
